# Metabolic plasticity in cancer activates apocryphal pathways for lipid desaturation

**DOI:** 10.1101/2020.06.07.139089

**Authors:** Reuben S.E. Young, Andrew P. Bowman, Elizabeth D. Williams, Kaylyn D. Tousignant, Charles L. Bidgood, Venkateswara R. Narreddula, Rajesh Gupta, David L. Marshall, Berwyck L.J. Poad, Colleen C. Nelson, Shane R. Ellis, Ron M.A. Heeren, Martin C. Sadowski, Stephen J. Blanksby

## Abstract

Fatty acid (FA) modifications, such as enzymatic desaturation and elongation, have long been thought to involve sequential and highly specific enzyme-substrate interactions, which result in canonical products that are well-defined in their chain lengths, degree of unsaturation and double bond positions.^1^ These products act as a supply of building blocks for the synthesis of complex lipids supporting a symphony of lipid signals and membrane macrostructure. Recently, it was brought to light that differences in substrate availability due to enzyme inhibition can activate alternative pathways in a range of cancers, potentially altering the total species repertoire of FA metabolism.^2,3^ We have used isomer-resolved lipidomics to analyse human prostate tumours and cancer cell lines and reveal, for the first-time, the full extent of metabolic plasticity in cancer. Assigning the double bond position(s) in simple and complex lipids allows mapping of fatty acid desaturation and elongation via hitherto apocryphal metabolic pathways that generate FAs with unusual sites of unsaturation. Downstream utilisation of these FAs is demonstrated by their incorporation into complex structural lipids. The unsaturation profiles of different phospholipids reveal substantive structural variation between classes that will, necessarily, modulate lipid-centred biological processes in cancer cells including membrane fluidity^3-5^ and signal transduction.^6-8^

---

Fatty acid (FA) metabolism is significantly altered within cancer cells, with increased FA unsaturation being pivotal in cell transformation, accelerated rates of proliferation, and augmented invasiveness.^6,9-13^ The introduction of carbon-carbon double bonds (DB) to specific sites along the FA chain is catalysed by three distinct desaturase enzymes, and results in structures with distinct physical properties and cellular functions.^5^ Alongside desaturation, elongation – a process that facilitates a two-carbon-unit chain extension through the elongase isoforms (ELOVL1-7) – can also have profound effects on FA properties and function.^1^ Together, these now-modified FAs are usually incorporated into various complex lipids, such as phospholipids, whereby molecular properties are imparted to the functional role of the lipid (*e.g*., structural membrane packing^4^ and fluidity^3^ or specific signal transduction^7,14,15^).

Previously, human prostate cancer (PCa) was shown to be characterised by increased ratios of monosaturated to saturated fatty acids relative to normal prostate tissue.^16^ These metabolic expressions were supported by transcriptomics showing elevated mRNA levels for both stearoyl-CoA desaturase-1 (SCD-1) and ELOVL7.^16,17^ Others have then focused on discerning the position of the double bond in monounsaturated FAs and found that within malignant PCa cells, the abundance of FA 18:1*n*-9 (*i.e*., an 18-carbon monounsaturated FA with the DB in the 9^th^ position from the methyl terminus) increases relative to the FA 18:1*n*-7 isomer compared with non-cancerous prostate cells.^18^

Notably, it was recently shown that a secondary desaturation mechanism can be activated to meet the metabolic needs of cancer.^2^ The canonical mechanism by which FAs undergo primary desaturation is through the oxygen-dependant SCD-1 enzyme introducing a DB at the Δ9 position (*i.e*., the 9^th^ carbon from the FA carboxylate terminus). However, under cellular stress fatty acyl desaturase 2 (FADS2) – a Δ6 desaturase – was also shown to possess primary desaturation capabilities, catalysing *n*-10 double bond formation in palmitic acid; a phenomenon usually observed only within lipids from hair and skin.^19^ Having two independent mechanisms for primary desaturation creates the opportunity for plasticity in cancer cells, which carries potential functional consequences for membrane structure and signalling. In order to elucidate if these changes are widespread, or confined to specific functions, it is imperative to identify the unsaturation profiles of complex lipids and not just the fatty acid building blocks. Unfortunately, technological limitations within conventional analysis, most notably gas-chromatography, first require FAs to be released from complex lipids by alkaline hydrolysis. The resulting analysis provides a FA profile integrated across the entire lipid pool, leading to a reduction of molecular-level information of the lipidome and, moreover, dilution of low abundant FAs that may have a particular association with a specific lipid class or composition. To overcome these limitations, next generation technologies, such as Paternò-Büchi derivatisation^20^ and ozone-induced dissociation (OzID),^21-23^ are instead able to discern DB positions within complex lipids. Such analyses not only provide links between unsaturation profiles and subcellular location or function, but also creates a platform in which lipid DBs can be imaged directly within a cellular-tissue matrix, revealing spatial distinctions between enzyme activities.

## Lipid unsaturation profiles in human prostate tissue

Lipid unsaturation distributions in human prostate sections were mapped using OzID coupled with matrix-assisted laser desorption ionisation mass spectrometry imaging (MALDI-MSI). Fig. 1A presents a spatial distribution between the monounsaturated phosphatidylcholine isomers PC 34:1*n-*9 (yellow) and PC 34:1*n-*7 (magenta), both metabolites of SCD-1, and the polyunsaturated PC 36:4*n-*6 (cyan), a metabolite formed through dietary FA substrates and FADS2 interactions. Comparison to the haematoxylin and eosin (H&E) stains of adjacent tissue sections reveal an increase of PC 34:1*n*-9 and a corresponding decrease of PC 36:4*n*-6 specific to the tumourous regions of the tissue. As these two metabolites are representative of two different desaturase enzyme activities, they are indicative of a change in substrate-enzyme interactions between tumour and adjacent non-tumour cells. The two PC 34:1 isomers are also found to have distinct distributions (Fig. 1A). In this instance however, both *n-*9 and *n-*7 forms arise from SCD-1 activity but require different FA substrates for their synthesis. In some regions across the tissue, signals consistent with a third DB isomer, the FADS2 metabolite PC 34:1*n*-10, were detected but abundances were insufficient to map distribution. Challenges in characterising less abundant metabolites are inherent to the MALDI-MSI technique (due to small sampling volumes) and therefore to increase the signal relative to the instrument background-noise, lipids were extracted from homogenised PCa tissues and subjected to direct infusion electrospray ionisation (ESI)-OzID analysis. Examination of lipid standards under identical conditions provided benchmarks for false-positive signals, thus confirming the presence of PC 34:1*n*-10 at low abundance in PCa tissues (Fig. 1B, green). Analogous double bond analysis of selected monounsaturated phosphatidylserine (PS) and phosphatidylethanolamine (PE) lipids also unveiled them as carriers of the unusual *n*-10 isomer within PS 36:1 and PE 36:1 (Fig. 2B, red and orange). Interestingly, signals arising from PE 34:1*n*-10 (blue) were insignificantly differentiated from background, indicating that incorporation of the *n-*10 FA may vary between lipid class and composition.

**Figure 1:**
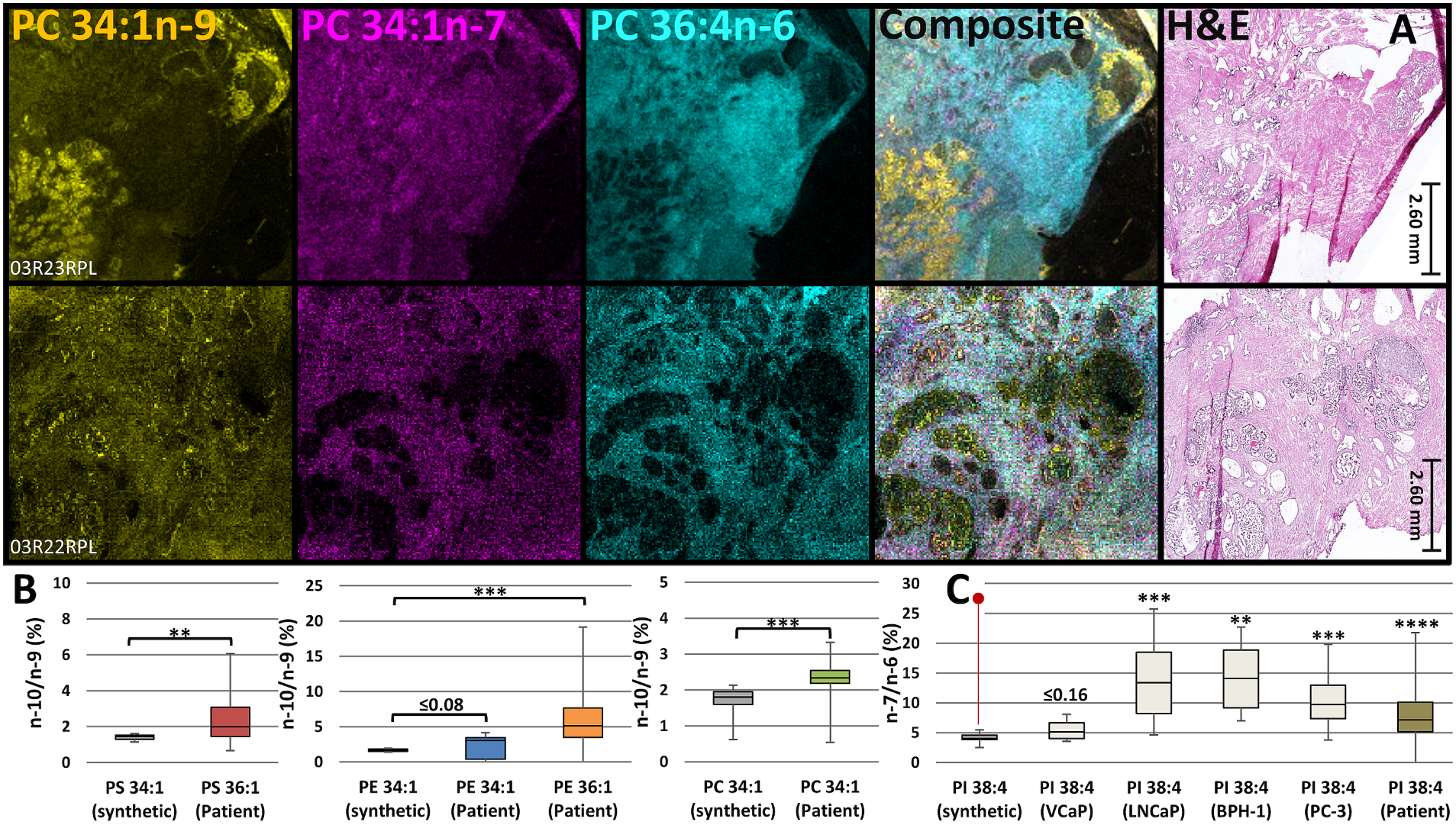
Metabolic differentiation of tumour in human prostate tissue. A) Two separate patient prostate lobe sections showing (left to right) MALDI-MSI-OzID distribution of PC 34:1*n*-9 (yellow), PC 34:1*n*-7 (magenta) and PC 36:4*n*-6 (cyan), a composite image of these lipids, and the adjacent tissue section H&E stained. (Magnification of tumour regions is shown, full images and spectral comparisons found in supplementary Fig. S1A-C). B) False positive *n*-10 signal arising from an *n*-9 standards (grey) compared to tissue lipid extracts from both prostate lobes of 8 patients (n=16); comparing PS 36:1*n*-10 (red), PE 34:1*n*-10 (blue) and PE 36:1*n*-10 (orange) and PC 34:1*n*-10 (green). C) *n*-7 Signal arising from PI 38:4*n*-6 standard (grey), four PCa cell lines (white) and tissue lipid extracts from both prostate lobes of 8 patients (n=16; gold). P-values compare biologically occurring PI 38:4*n*-7 against false positive *n-*7 signal from synthetic PI 38:4*n*-6 standard (red line marker). (Displayed p-values: **≤0.01, ***≤0.005, ****≤0.0001).

**Figure 2:**
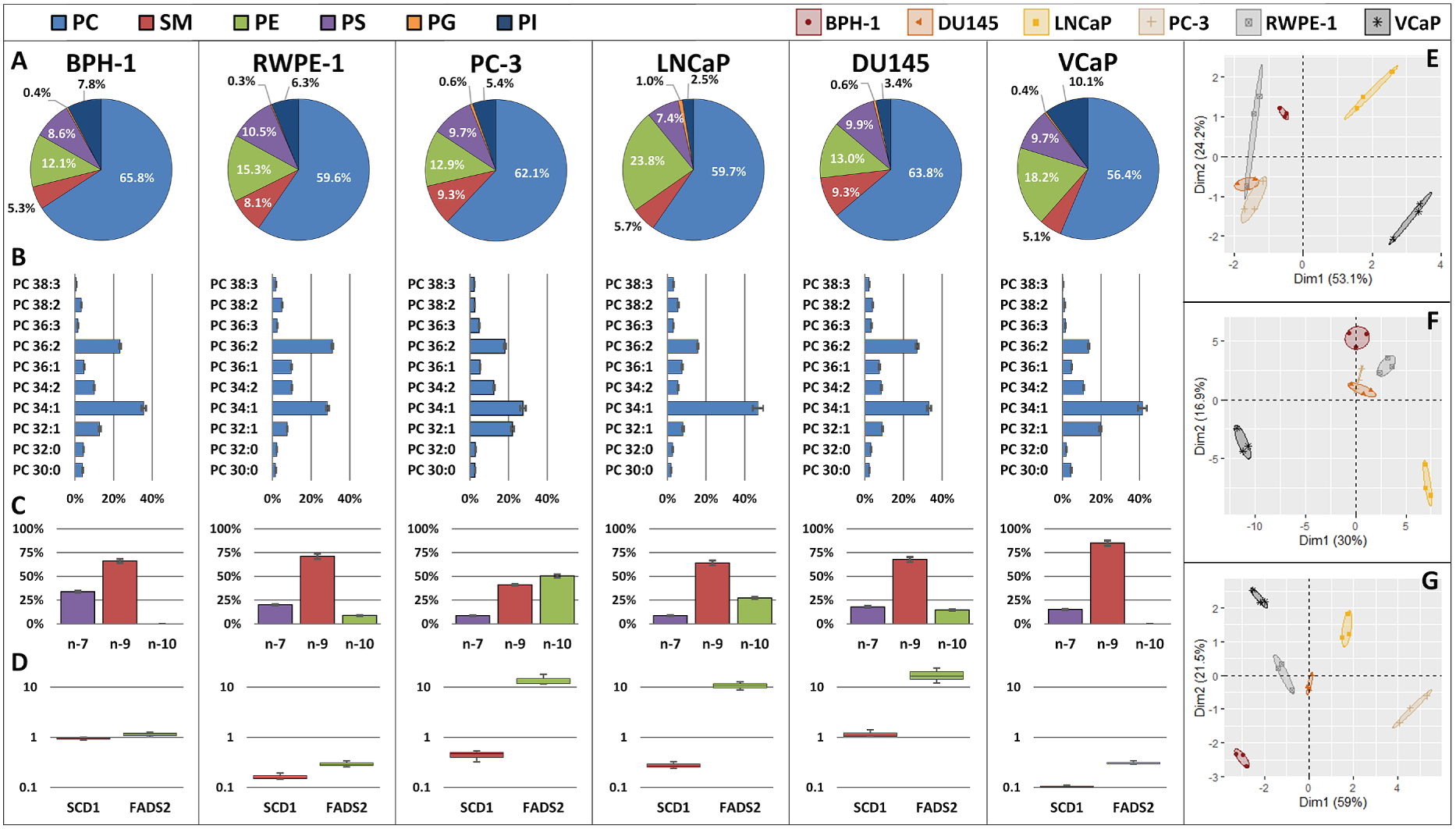
Lipid composition and targeted transcripts of normal prostate (BPH-1, RWPE-1) and cancer (PC-3, LNCaP, DU145, VCaP) cell lines (n=3). A) phospholipid profiles across 6 major classes (PC, SM, PE, PS, PG, PI). B). Normalised abundance of major PC lipids quantified at the sum composition level. C) Normalised OzID signal intensity from PC 34:1 isomers with double bonds at *n*-7, *n*-9 and *n*-10. D) qRT-PCR derived transcript expression of two desaturases (SCD-1, FADS2) as a fold-change relative to the normal prostate cell (BPH-1) mRNA expression. E-G) PCA dimensionality reduction for cell line differentiation based on (E) 6 phospholipid classes, (F) sum composition of 10 PCs and (G) double bond isomer distributions from PC 32:1, 34:1 and 36:1. Ellipses display 95% confidence interval, with the percentage of data represented by the dimensionality reduction being displayed as axes. A-C are displayed in relative totals for inter-cell line normalisation. (Mean and 95% confidence interval displayed).

Examination of double bond profiles from polyunsaturated lipids in PCa tissue extracts revealed that a previously unreported *n-*7 isomer was present alongside the highly-active signalling lipid, phosphatidylinositol (PI) 38:4*n*-6.^7^ The use of high-resolution mass spectrometry and OzID allowed for the unambiguous assignment of PI 38:4*n*-7 (see supplementary Fig. S11A-B). The ratio of signals arising from this lipid and the canonical PI 38:4*n*-6 are displayed in Fig. 1C, which reveals that tissue and PCa cell line extracts display PI 38:4*n*-7 around 10-15% above any background signal arising from the *n*-6 synthetic standard. Further evidence for apocryphal desaturation in PI 38:4 was obtained via comparison to (and between) PCa cell line lipid extracts (Fig. 1C) and PCa cell line fatty acyl double bond analysis (*vide infra* – Fig. 5B). These findings present for the first time the existence of the *n-*10 monounsaturated FAs and *n-*7 polyunsaturated FA within human primary prostate tumours, and implies distinctive changes to desaturase enzyme activity or expression across tumour tissues, which in turn is impacting lipids associated with intracellular membrane structures and signals.^14,24,25^

**Figure 3:**
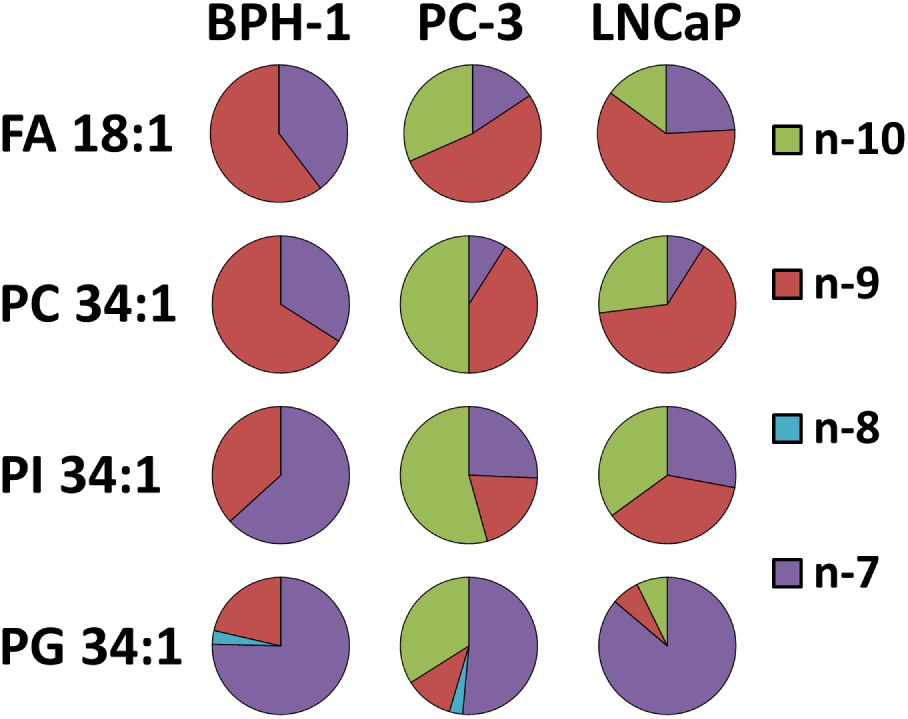
Variation of DBs across the hydrolysed fatty acyl pool (GC-MS) and PC, PI and PG (OzID) complex lipids for normal prostate BPH-1 cells and PC-3 & LNCaP cancer cells. (mean values displayed (n=3))

**Figure 4:**
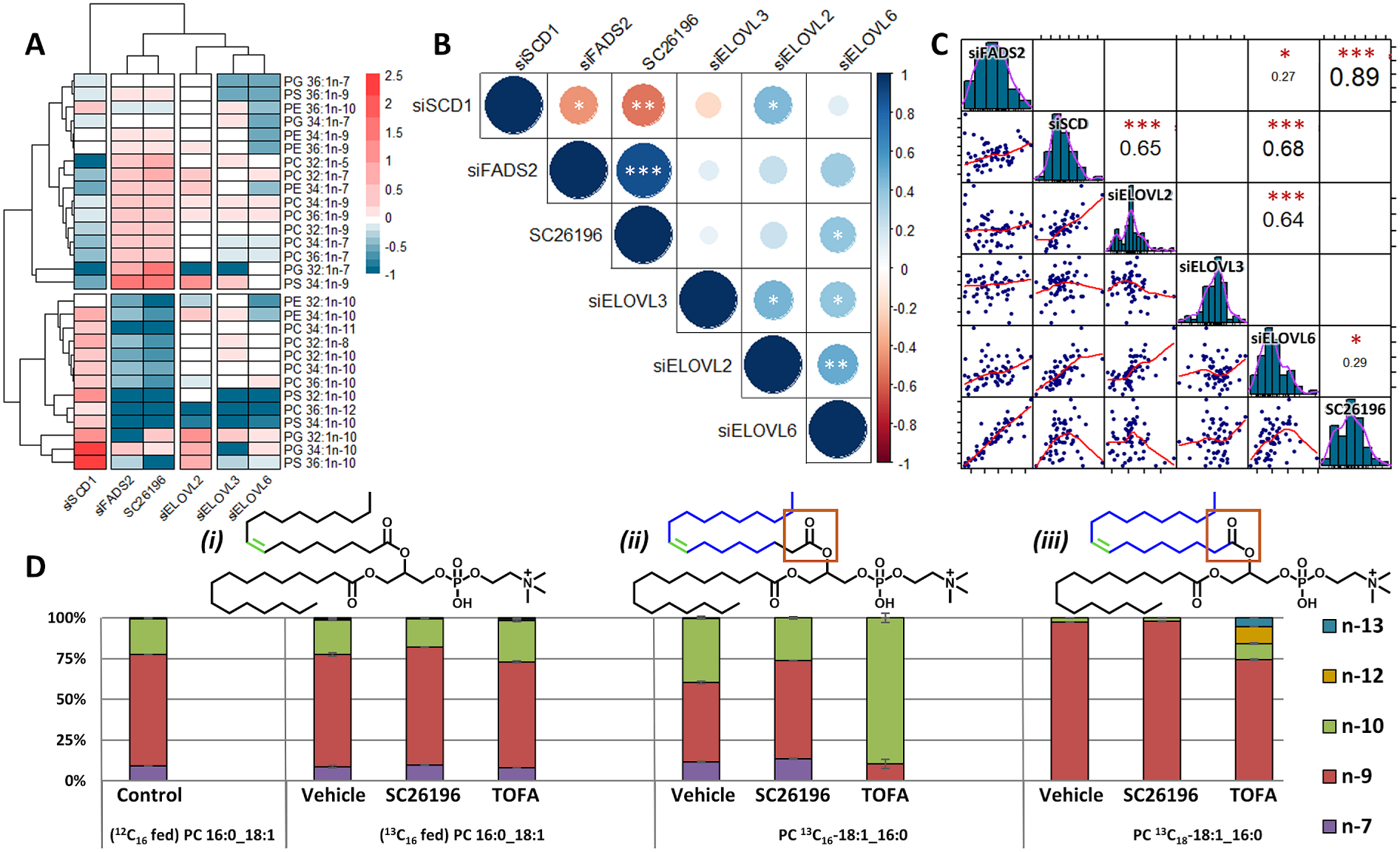
Enzyme/metabolite correlations and increased enzyme plasticity. **A**) Heat map of treated LNCaP cells showing change in monounsaturated phospholipid double bond isomers relative to control (untreated-LNCaP). **B**) Correlation matrix using numeric data from A to show positive (blue) and negative (red) correlations (n=29). **C**) Bivariate analysis, histograms and correlation matrices for treated-LNCaP lipid sum compositional analysis compared to control (n=60). **D**) PC 34:1 double bond fractional distribution profile from *de novo* lipogenisis (left and mid-left),^13^C_16_-palmitate tracing with de novo ^12^C-acyl-CoA modifications (mid-right) and ^13^C_18_-stearate tracing (right). Lipid structures are displayed above, with isotopic carbons in blue. (n=2, 95% confidence interval error). (Displayed p-values: *≤0.05, **≤0.01, ***≤0.001).

**Figure 5:**
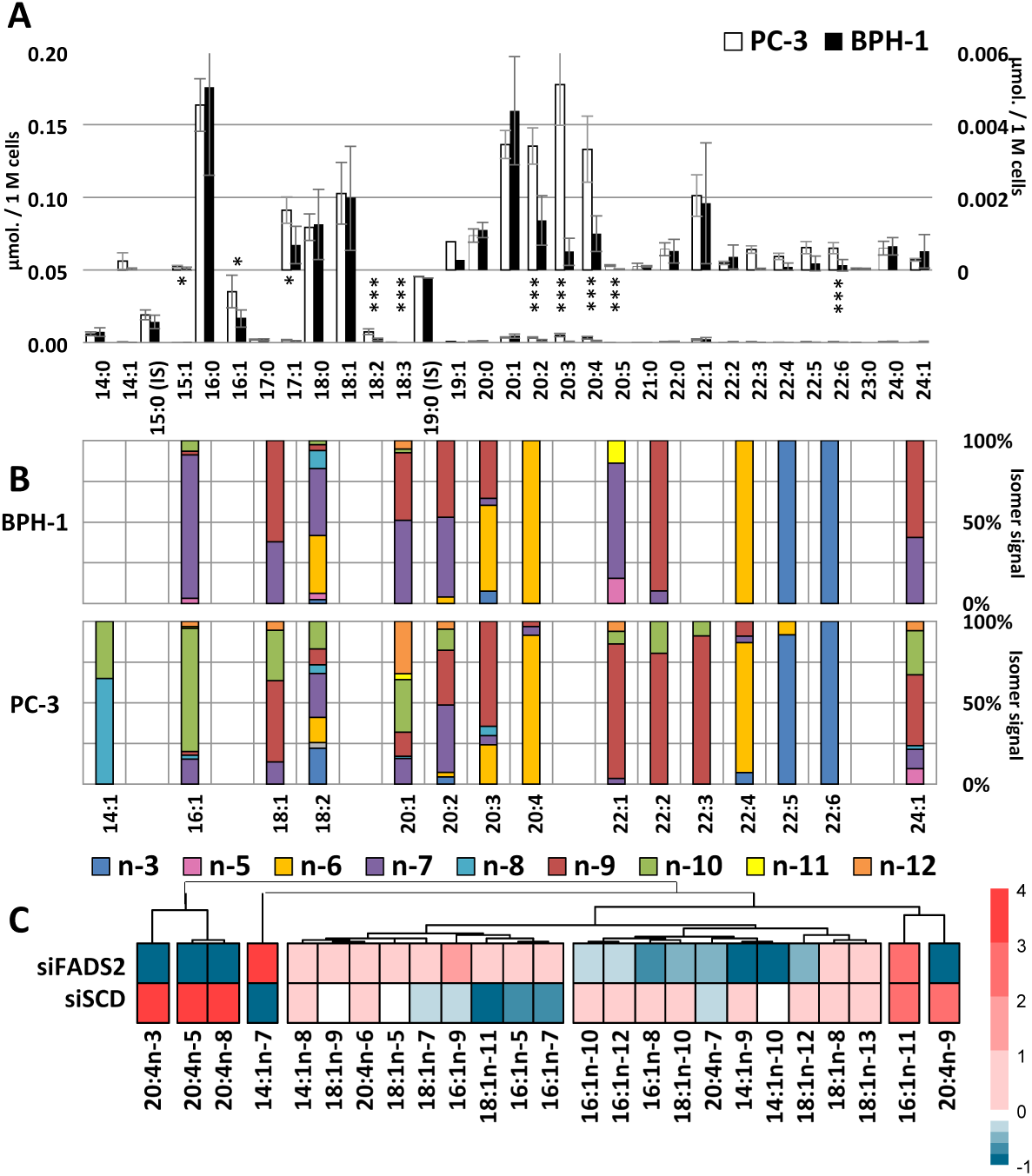
Pooled fatty acyl quantitation and double bond analysis of normal prostate cells (BPH-1), cancer cells (PC-3) and genetic silencing of desaturases in cancer cells (LNCaP). **A**) Cell count normalised quantitative sum composition fatty acyl analysis with magnification insert (n=3; 95% confidence interval error; p-values: *≤0.05, **≤0.01, ***≤0.005). **B**) Double bond fractional distribution profiles for 15 mono and polyunsaturated fatty acids from the hydrolysed lipid pool (n=3; mean fractional distribution displayed). **C**) Heat map of treated LNCaP cells showing change in mono and polyunsaturated fatty acid double bond isomers from the hydrolysed lipid

## Widespread remodelling of unsaturation in cellular lipidomes

Although patient samples provide powerful insight into biologically relevant responses to disease, they also carry with them a large number of uncontrolled variables. In order to minimise these variables and directly explore the impact desaturase and elongase enzymes have on the FA metabolic repertoire of cells, four prostate cancer cell lines from metastatic deposits (PC-3, LNCaP, DU145 and VCaP; hereinafter referred to as cancer cells) and two benign immortalised prostate epithelial cell lines (BPH-1 and RWPE-1; hereinafter referred to as normal prostate cells) were cultured under identical conditions. The cells were comprehensively characterised by: conventional lipidomics to obtain full profiles of molecular lipids (supplementary Fig. S2A-C) allowing comparison between phospholipid classes (Fig. 2A) and sum compositions, as shown for the dominant phosphatidylcholine class in Fig. 2B; ozone-induced dissociation to reveal the contributions of double bond isomers to the populations of monounsaturated lipids isomers (supplementary Fig. S4A-D), as shown for the most abundant lipid (PC 34:1 Fig. 2C); gas chromatography-mass spectrometry of the saponified extracts yielding the FA profile integrated over the lipid pool (supplementary, Fig. S3); and transcriptomics for fatty acyl desaturase expression (Fig. 2D). Data represented in Fig. 2A-C were summarised using an unsupervised multivariate analysis in the form of principal component analysis (Fig. 2E-G) to differentiate cell line profiles.

Principal component analysis (PCA) of the phospholipid profiles (Fig. 2E) and the abundance of major PC lipids (Fig. 2F) both reveal poor sample cluster separation including cluster intermingling, and mixed adequacy in terms of the explanation of variance. This is seen where the phospholipid profile data (Fig. 2E) can be reasonably explained by the first two principle components (∼77%), whereas the variation in PC sum composition (Fig. 2F) is weakly explained (∼47%). Combined, this indicates that conventional lipidomics contains unexplained factors of variation that contribute to an inability to distinguish cancer and normal prostate cell lines. Conversely, PCA of the DB isomer profiles from major monounsaturated PC species (Fig. 2G) presents independent sample clustering, with the variation being well captured by the first two principle components (∼81%). The clustering of PC-3, LNCaP and VCaP cancer cell lines also displays strong negative association across both principal dimension axes against the normal prostate cell line BPH-1. Overall, multivariate analysis based on double bond profiles clearly distinguishes between cancer and normal prostate cell lines (Fig. 2G), while analogous analyses based on conventional lipidomics data are unable to do so (Fig. 2E-F). Differentiation of cell lines based upon PCA of double bond isomers is influenced strongly by the presence of unusual *n*-10 lipids. As evidenced in Fig. 2C, PC 34:1*n*-10 contributions range from being absent in BPH-1 and VCaP cells to 50% of the isomer fraction in PC-3 – remarkably representing up to 8.5 mol% of total phospholipid without impacting cell viability.

The high proportion of PC 34:1*n*-10 in some cell lines (Fig. 2C, green) combined with the significant variation between unsaturation profiles (Fig. 2G) implies change in the relative activity and/or expression of the two desaturase enzymes, SCD-1 and FADS2. Noting the absence of *n*-10, the BPH-1 normal prostate cell line was used as a reference to compare the transcription of SCD-1 and FADS2 enzymes across all cell lines. Fig. 2D shows a 10-fold increase in the transcription of FADS2 for the cancer cell lines PC-3, LNCaP and DU145, where the contribution of *n-*10 unsaturation is most pronounced (Fig 2C, green). In contrast, RWPE-1 and VCaP cell lines present lower transcription of FADS2 (relative to BHP-1) and are characterised by a decreased abundance and complete absence, respectively, of the n-10 lipid isomer. Interestingly, these two cell lines also share a similar transcript expression of SCD-1 and FADS2 despite the clear difference in PC 34:1 DB isomer profiles (notably the absence of *n*-10 in VCaP). This apparent discrepancy between desaturase transcript expression and resulting metabolites points to competition between desaturase enzymes for common substrates with the potential for additional competition through substrate elongation. Such competition is also evident across the cell lines in the association between increased PC 34:1*n*-10 (Fig. 2C green) and decreased abundance of PC 34:1*n*-7 (Fig. 2C purple). Both isomers share 16:0 as a common substrate and thus competition between the independent SCD-1 or FADS2 reactions is apparent.^26,27^ Similarly, observing the VCaP unsaturation profile relative to BPH-1 (Fig. 2C), the decreased presence of PC 34:1*n*-7 (Fig. 2C purple) is matched by an increased abundance of PC 34:1*n*-9 (Fig. 2C red). As SCD-1 desaturation can yield either *n*-7 or *n*-9 fatty acids depending on the substrate (16:0 or 18:0, respectively), and ELOVL6 is known to elongate 16:0 to 18:0,^28,29^ the relative increase of VCaP PC 34:1*n*-9 suggests competitive 16:0 metabolism by either direct-desaturation or a combination of elongation/desaturation reactions.

Given the potential for competition between desaturation and elongation pathways, the association between sites of unsaturation and specific fatty acyl chain length(s) becomes critical for mapping metabolism. Fatty acyl compositional analysis, based on collision-induced dissociation (CID) combined with OzID, reveals that cell line PC 34:1 is comprised of >95% PC 16:0_18:1 (supplementary Fig. S5A-C) inferring that the vast majority of the *n*-7, *n*-9 and *n*-10 double bonds shown in Fig. 2C are carried by 18:1 fatty acyl chains. Although somewhat varied, fatty acyl compositional analysis (based on negative polarity CID) of other lipid classes also revealed a majority contribution of 16:0_18:1 to the 34:1 species of PI, PS, PE and phosphatidylglycerol (PG), with average contributions of 78%, 58%, 77% and 96%, respectively (supplementary Fig. S5A-C). When exploring the double bond positions tentatively assigned to the 18:1 chain of these phospholipids however, the results show a large perturbation between cancer and normal prostate cells. Fig. 3 displays the fractional contributions of DB isomers within three lipid subclasses across BPH-1 cells and PC-3 and LNCaP cancer cells. Because the majority of these monounsaturated lipids carry the 18:1 acyl chain, the isomer distribution of FA 18:1 from the hydrolysed lipid pool was also derived from GC-MS (representative chromatograms shown in supplementary Fig. S6A). The results summarised in Fig. 3 highlight a significant contribution of FA 18:1*n*-10 to the total FA 18:1 pool in PC-3 (32%) and LNCaP (15%) cancer cell lines while it is completely absent in the BPH-1 normal prostate cells. Any contribution of FA 18:1*n-*10 (or FA 16:1*n-*10) from exogenous sources was explicitly excluded by rigorous analysis of cell culture media and other controls (supplementary Fig. S6A-B). Comparing the isomer factions of FA 18:1 with the major 18:1 bearing lipids (*i.e*., PC 34:1, PI 34:1, PG 34:1), two things become acutely apparent: (i) DB isomer proportions vary between phospholipid class (*i.e*., Fig. 3: PC vs. PI vs. PG), and (ii) the PG 34:1 of BPH-1 and PC-3 carries an additional n-8 isomer. These observations, which would be overlooked by pooled FA analysis alone, indicate a degree of specificity in the unsaturation profile across different phospholipid classes and the potential for greater diversity in lipid unsaturation than previously considered.

## Expanded substrate accommodation of elongase and desaturase enzymes

While being a major contributor to the lipid pool for cancer cell lines, the biosynthetic origin of the FA 18:1*n*-10 building block remains to be clarified. The association of 18:1*n*-10 with the FADS2 desaturase is apparent (*cf*. Fig. 2) and the canonical pathway would proceed via Δ6-desaturation of 16:0 to 16:1*n*-10 with subsequent elongation by a hitherto unassigned enzyme. To identify this elongase (hereinafter referred to as ELOVLx) along with other mechanisms for cancer cell plasticity, the LNCaP cancer cell line was chosen as a model based on its high expression of FADS2 and *n*-10 abundance. Cells were subject to gene silencing by small interfering RNA (siRNA; Western blot, qRT-PCR and cell confluence in supplementary Fig. S7 and primer sequences in supplementary Fig. S12) and enzyme inhibition before monounsaturated phospholipids were characterised by OzID.

Relative to the untreated LNCaP control, the heatmap of Fig. 4A displays 29 monounsaturated phospholipids for which double bond positions are assigned and their abundance shift in response to treatment with desaturase gene silencing (siSCD-1, siFADS2), elongase gene silencing (siELOVL3, siELOVL6, siELOVL2; to broadly represent canonical elongation activity for saturated, monounsaturated and polyunsaturated fatty acyl substrates, respectively)^1,29,30^ and a FADS2 inhibitor (SC26196 – a potent FADS2 inhibitor that was found to prevent the conversion of linoleic acid to arachidonic acid by ≥95%).^31^ Horizontal hierarchical clusters in Fig. 4A identify that siSCD-1 is distinct from siFADS2 and SC26196 which, unsurprisingly due to their activity, cluster together. Similarly, all siELOVL treatments are clustered, with siELOVL3 and siELOVL6 showing a closer relationship to each other than to siELOVL2. Vertical hierarchical cluster analysis identifies of two main groups that correspond to SCD-1 related (*i.e*., predominately *n-*7 & *n*-9) and FADS2 related (*i.e*., predominately *n*-10) lipid products. Amongst the three desaturase treatments, inhibition or gene silencing of one enzyme causes an apparent increase in metabolic products from the alternative enzyme *e.g*., the relative abundance of PG 32:1*n*-7 and *n*-10 are observed to have an inverse response during SCD-1 and FADS2 gene silencing. In addition to the establishment of these inverse metabolite relationships, gene silencing (or inhibition) of FADS2 and SCD-1 also creates further augmentation of monounsaturated DB profiles with sites of unsaturation identified at *n*-5, *n*-8, *n*-11 and *n*-12.

To observe statistical correlations between the monounsaturated lipid profiles generated in response to treatments (Fig. 4A), Pearson’s correlation was implemented, and the resulting correlation matrix is presented in Fig. 4B. The circle size and colour intensity display the magnitude of the positive (blue) and negative (red) correlations, with statistical significance indicated. For example, the inverse relationship between siFADS2/SC26196 and siSCD-1 identified previously in Fig. 4A, is observed in Fig. 4B to be statistically significant negative correlations (p<0.05 / p<0.01, respectively). Similar to the horizontal hierarchical clusters of Fig. 4A, the desaturase and elongase treatments group separately at the top and bottom matrix vertices, indicating similarity in response to like-enzyme treatments. Within the matrix region corresponding to the correlation of desaturase treatment profiles to elongase enzyme treatment profiles, only two statistically significant positive correlations can be observed, siSCD-1/siELOVL2 and SC26196/siELOVL6.

As previously mentioned ELOVL6 is mainly responsible for the elongation of 16:0 to 18:0,^29^ therefore downregulation of this activity by siELOVL6 treatment would cause accumulation of 16:0, palmitic acid. To prevent palmitic acid lipotoxicity, canonical activity of SCD-1 would drive desaturation to FA 16:1*n*-7. Likewise, inhibition of the FADS2 enzyme by SC26196 would prevent palmitic acid from undergoing FADS2 desaturation to FA 16:1*n*-10 and instead consolidate desaturation activity through SCD-1 to FA 16:1*n*-7. This would in turn present a positive correlation to siELOVL6 (to view the metabolic impact that treatments have on fatty acyl modification networks, *cf*. supplementary Fig. S8A-E).The silencing of SCD-1 desaturation (and hence increased abundance of FADS2 related lipid products) however, has no reported reason to display positive correlation to siELOVL2. Given that the silencing of ELOVLx would prevent the elongation of FA 16:1*n*-10 to FA 18:1*n*-10, the accumulation FA 16:1*n*-10 would display positive correlation to a profile dominated by FADS2 related products, such as with siSCD-1. Therefore, it is suggested that ELOVL2 is indeed ELOVLx and is responsible for the elongation of FA 16:1*n*-10 to FA 18:1*n*-10.

Akin to the major role of the FADS2 enzyme in polyunsaturated lipid synthesis, ELOVL2 has only previously been implicated in polyunsaturated fatty acyl elongation, but never in the elongation of monounsaturated fatty acids.^29,32^ Hence, in order to validate the finding that ELOVL2 catalyses elongation of FA 16:1*n*-10 to FA 18:1*n*-10, exploration into the impact of treatment on the wider lipidome were conducted. Relative to the control, Fig. 4C displays the correlations between the lipid profiles of treatments, consisting of 60 saturated, monounsaturated and polyunsaturated sum composition phospholipids (PC, PE, PS, PG). As before in Fig. 4B, the siELOVL2 and siELOVL6 treatments of Fig. 4C display statistically significant positive correlations to desaturase silencing and inhibition, informing of moderate to high degrees of lipid profile similarity. Additionally, the profile from siELOVL6 also shows correlation with the siSCD-1 treatment profile. Similar to the logic used previously, accumulation of palmitic acid by way of silencing ELOVL6 would lead to an increase in shorter-chain monounsaturated lipids, which as substrates for further desaturation, would in turn increase the abundance of polyunsaturated lipids. Comparably, silencing SCD-1 would increase FADS2 activity, which would increase polyunsaturated lipid abundance due to its major canonical role in polyunsaturated lipid synthesis. Hence, the profiles of siELOVL6 and siSCD-1 would present similarly. Although slight skewing can be observed, all histograms present a normal distribution of the data, with the exception of siELOVL2. This bimodal distribution suggests that silencing of ELOVL2 has two distinctive impacts on the lipidome, which given its known role in the elongation of polyunsaturated lipids, provides further indication for an additional metabolic role by ELOVL2. Therefore, the combined data from Fig. 4B & 4C suggest that ELOVL2 is the most likely candidate for the unassigned elongase responsible for apocryphal elongation of 16:1*n*-10.

To explore the sequence of elongation and desaturation events, LNCaP cancer cells were supplemented with ^13^C-labelled fatty acids. Incorporation of ^13^C_16_-palmitate and ^13^C_18_-stearate tracers into PC ^13^C_16_-18:1_16:0 (*m/z* 798) and PC ^13^C_18_-18:1_16:0 (*m/z* 800), respectively, was confirmed by high resolution tandem mass spectrometry (supplementary Fig. S13A-B). The presence of PC ^13^C_16_-18:1_16:0 (*m/z* 798) arising from ^13^C_16_-palmitate supplementation indicates intracellular elongation with installation of two unlabelled carbons (see chemical structures, Fig. 4Dii). This conclusion is supported by OzID fragmentation of the mass-selected isotopologue that assigns the location of the two unlabelled methylene units between the site of unsaturation and the carboxylate moiety. The PC ^13^C_16_-18:1_16:0 metabolite is also characterised by a distribution of sites of unsaturation in the labelled 18:1 chain showing contributions from *n-*7, *n-*9 and *n-*10 isomers (Fig 4Dii left). In contrast, the fully labelled PC ^13^C_18_-18:1_16:0 derived from ^13^C_18_-stearate is characterised by near exclusive *n-*9 unsaturation in the labelled chain (Fig 4Diii left). These tracer results infer direct desaturation of stearate giving rise to 18:1*n*-9, whilst 18:1*n*-10 follows a desaturation-elongation sequence analogous to the canonical formation of 18:1*n*-7. Within the same experimental system, the suppression of *n*-10 unsaturation in PC ^13^C_16_-18:1_16:0 was observed upon the inhibition of FADS2 (SC26196), further demonstrating Δ6-desaturation of palmitate prior to elongation (Fig. 4Dii mid). In contrast, inhibition of the same enzyme in the presence of the ^13^C_18_-stearate tracer yields no observable change to the unsaturation profile of PC ^13^C_18_-18:1_16:0 relative to vehicle (Fig. 4Diii mid) and corroborates direct desaturation by SCD-1 to form *n*-9. For comparison, the inhibition of SCD-1 (TOFA – originally an ACC1 inhibitor found to have potent SCD-1 inhibition)^33^ in the presence of the ^13^C_16_-palmitate tracer led to complete depletion of *n*-7 and a significant reduction of *n*-9 within PC ^13^C_16_-18:1_16:0, indicating the cessation of Δ9-desaturation of palmitate and stearate, respectively – an effect that appears to trigger compensation through desaturation by FADS2 yielding an unsaturation profile dominated by *n*-10 (Fig. 4Dii right). Such compensation is also observed when inhibiting SCD-1 in the presence of the ^13^C_18_-stearate tracer, which noticeably the reduces *n*-9 unsaturation within PC ^13^C_18_-18:1_16:0 and promotes formation of an unusual suite of *n*-13, *n*-12 and *n*-10 isomers indicative of direct Δ5-, Δ6-, Δ8-desaturation of the tracer to (Fig. 4Diii right). Given the well described Δ9-fidelity of SCD-1,^34^ these results represent a scientific first in providing evidence for direct desaturation of stearate by desaturases *other* than SCD-1. Specifically, the findings demonstrate that 18:1*n*-10 can be synthesised directly from stearate by the hitherto apocryphal Δ8-desaturation activity of FADS2. Although Δ8 desaturases do occur naturally in some marine micro algae,^35^ mammalian Δ8-desaturation of saturated fatty acids has only been observed exogenously on skin.^19^ From the isotope labelling experiments above, the presence of *n*-12 accompanying *n*-10 monounsaturation within PC ^13^C_18_-18:1_16:0 reveals that FADS2 may also be able to simultaneously exhibit Δ8-desaturation activity alongside canonical Δ6 activity. This result could indicate differential activities of the FADS2 enzyme arising from intracellular compartmentalisation, local macrostructural environment or substrates (*i.e*., carrier of the stearates). The proclivity of FADS1 (a third mammalian desaturase) towards Δ5 activity within polyunsaturated FAs, may help explain the curious observation of the *n*-13 monounsaturated PC ^13^C_18_-18:1_16:0. In this instance however, the desaturase would require exertion of apocryphal metabolic behaviour to instead act upon saturated substrates at the Δ5 position to initiate primary desaturation. This same Δ5 motif is observable in Fig. 4A, wherein gene silencing of SCD-1 or FADS2 causes respective amplification or elimination of an unusual PC 34:1*n*-11. This variable presence may be explainable by Δ5-desaturation of palmitic acid substrates by FADS1 during times of lesser competition for substrate by SCD-1.

## Evidence for alternate pathways in lipid desaturation

Analysis of monounsaturated phospholipids within PCa cell lines revealed an unexpected diversity of lipid isomers (*vide supra*). To visualise this diversity across the entire fatty acyl pool, cell line extracts were hydrolysed, derivatised with 1-[4-(aminomethyl)phenyl]pyridinium (AMPP+) and subjected to OzID. This direct-infusion MS approach was found to be more sensitive to low abundant FAs and enabled unambiguous assignment of double bond positions in the absence of reference standards.^36^ Comparison of AMPP+ against GCMS results can be found in supplementary Fig. S9. Fig. 5A displays molar abundance of sum composition FAs for the two representative cell lines, PC-3 (cancer) and BPH-1 (normal prostate), across three orders of magnitude. When isomeric contributions are ignored, the abundance of the saturated and monounsaturated fatty acids are well conserved between cell lines. In contrast, several low abundant polyunsaturated fatty acids show significant changes in cellular abundances. *e.g*., FA 20:3, FA 20:4 and FA 22:6 are more abundant in PC-3 compared to BPH-1.

As was observed within phospholipids, ostensibly similar sum composition profiles can mask significant isomeric differences (*cf*. Fig. 2). Fatty acyl isomer profiles were thus extracted from OzID analysis and are presented in Fig. 5B. Among the monounsaturated even-chain fatty acids of PC-3, the apocryphal elongation of FA 16:1*n*-10 to FA 18:1*n*-10 that has been shown throughout appears not to be a terminal step in the elongation of the *n*-10 family. Instead, this family includes longer-chain fatty acids such as 20:1, 22:1 and 24:1, representing successive chain elongation (*vide infra* Fig. 6*vi*). Alongside further elongation of FA 18:1*n*-10 substrates, the presence of FA 18:2*n*-10, the skin lipid sebaleic acid, indicates the possibility for additional desaturation using this same FA 18:1 substrate and demonstrates a point of divergence in enzymatic activity surrounding the FA 18:1*n*-10. Further combinations of elongation and desaturation of FA 18:2*n*-10 subsequently lead to additional *n*-10 polyunsaturated fatty acyl species observed within FA 20:2, FA 22:2, FA 22:3 (Fig. 5B; *vide infra* Fig. 6*vii*). A similar enzyme branch point can be observed with FA 18:1*n*-7, whereby the presence of *n*-7 isomers within FA 20:1, FA 22:1 and FA 24:1 (*vide infra* Fig. 6*iv*) indicates elongation, while *n-*7 contributions to polyunsaturated FA 18:2, FA 20:2, FA 20:3, FA 20:4 and FA 22:4 demonstrate coalescent elongation and desaturation (Fig. 5B; *vide infra* Fig. 6*iv*). These polyunsaturated fatty acids are remarkable not only because they are formed exclusively through *de novo* synthesis, but because they bear structural similarity with the biologically active FAs such as arachidonic acid or adrenic acid.

**Figure 6:**
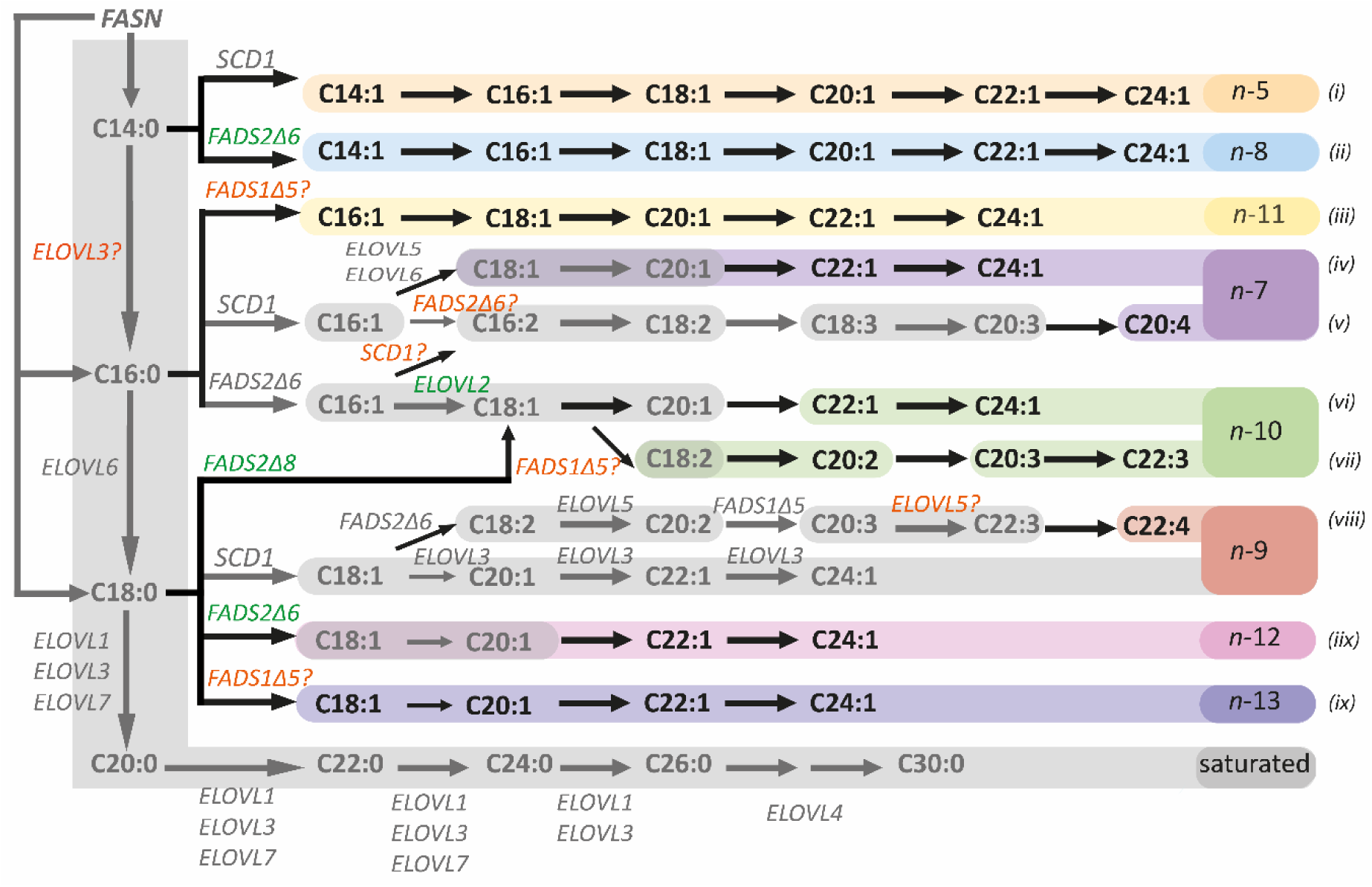
Fatty acyl desaturase and elongase pathways rationalising the fatty acyl species observed within this study. Greyed-out shows previously defined pathway^6^ and coloured presents new FA species found in human prostate cell lines. Grey text represents known possibilities for mammalian enzyme activity, green text shows newly confirmed enzyme results for human prostate cell lines, and orange text displays enzymes requiring further confirmation.

The multiple isomeric species observed within the FA 20:4 profile of PC-3 in Fig. 5B, including FA 20:4*n*-7 – an isomer also seen within PI 38:4*n*-7 of PCa tissues and cell lines (*cf*. Fig. 1C) – and their absence from normal prostate BPH-1 cells, represents an overall increase in the isomer diversity of polyunsaturated fatty acids in PC-3 and other cancer cell lines (supplementary Fig. S10A-C). This increased isomeric complexity of polyunsaturated fatty acyl profiles appears to correspond with an increase in isomer speciation created during the primary desaturation of shorter chain saturated fatty acids (*i.e*., FA 14:0, FA 16:0 and FA 18:0) and can be clearly observed with desaturase gene silencing (Fig. 5C). Relative to the untreated LNCaP control, gene silencing of cellular FADS2 decreases the isomeric complexity of monounsaturated FA 14:1, FA 16:1 and FA 18:1 by heavily reducing the presence of *n*-8, *n*-10 and *n*-12 isomers while simultaneously eliminating of FA 20:4*n*-7 and consolidating the FA 20:4 profile in FA 20:4*n*-6, arachidonic acid. Conversely, the FA profile from SCD-1 gene silencing appears to have greater isomeric speciation within FA 14:1, FA 16:1 and FA 18:1, which in turn has no impact on polyunsaturated FA 20:4 isomer complexity as both FA 20:4*n*-7 and FA 20:4*n*-6 are found to be present.

The expanded isomeric complexity afforded by apocryphal primary desaturation also creates an array of unusual monounsaturated fatty acids, which individually, infer explicit enzyme-substrate activity that can be organised into a network of metabolites (Fig. 6). For example, FA 24:1*n*-5, which is present in the fatty acids of PC-3 in Fig. 5B, is consistent with canonical SCD-1 Δ9-desaturation of FA 14:0 to initially yield FA 14:1*n*-5, which along with other chain length intermediates in this *n*-5 family (Fig. 6*i*), can be observed in supplementary Fig. S10A-E. In another example of divergent enzyme action, this same FA 14:0 substrate is seen to undergo Δ6-desaturation by FADS2 to form FA 14:1*n*-8, a product that is observed within PC-3 in Fig. 5B and is highly sensitive to the gene silencing of FADS2 (Fig. 5C). The presence of this DB position in varying chain lengths (supplementary Fig. S10A-E) indicates that this family of fatty acids has the ability to extend to at least FA 24:1 (Fig. 6*ii*) and has the potential for incorporation into intact phospholipids (*cf*. Fig. 3). Shifting to FA 18:0 as a substrate, FADS2 Δ6-desaturation yields FA 18:1*n*-12, which along with the 20:1, 22:1 and 24:1, is a FA family that can be observed in the fatty acids of PC-3 (Fig. 5B; Fig. 6*iix*). The 18:1*n*-12, is also seen to form under SCD-1 inhibition of intact lipids with incorporated heavy-labelled FAs (*cf*. Fig. 4D) and shows sensitivity to FADS2 gene silencing with apparent decreases relative to the control (Fig. 5C). Seemingly beyond the reported Δ9 and Δ6 positional activities for primary desaturation via SCD-1 or FADS2, two additional monounsaturated FA families can be observed in Fig. 5B-C, *n*-11 and *n*-13 (*cf*. Fig. 4A & D). The emergence of both these families is consistent with Δ5 primary desaturation of either FA 16:0 or FA 18:0 substrates to initially yield FA 16:1*n*-11 and FA 18:1 *n*-13, respectively, before chain elongation with intermediates observed out to FA 24:1 (supplementary Fig. S10A-E; Fig. 6*iii* & 6*ix*). Although no mammalian desaturase has been reported to facilitate Δ5 primary desaturation for the synthesis of monounsaturated fatty acids, Δ5-desaturation is commonplace during the synthesis of polyunsaturated fatty acids and undertaken by FADS1. Given the evidence shown throughout that FADS2 has the ability to exhibit a high degree of plasticity, and its near-identical genetic similarity to FADS1,^37^ it is speculated that FADS1 can also exhibit plasticity and is responsible for apocryphal Δ5 primary desaturation to form the *n*-11 and *n*-13 fatty acid families.

## Origin and consequence of desaturation plasticity

Exploration of the double bonds within cancer cell lipids in combination with gene silencing and isotope tracing experiments has revealed that mammalian cellular fatty acyl modification is far more dynamic than previously considered. It is important to note that the flow chart represented in Fig. 6 presents the metabolic capabilities of mammalian cells, with some FA families arising due to extreme metabolic stress (*e.g*., SCD-1 gene silencing allowing FADS1 *n*-13 formation). Because cancer is known to be severely disruptive to cellular fatty acyl metabolism however, it stands to reason that this expanded network of possibilities for FA modification provides plasticity for changes to lipid uptake or FA enzyme expression in PCa cells and tumour tissue.

Although SCD-1 and FADS2 largely show preference for canonical Δ9 and Δ6 sites during primary desaturation, a change in substrate availability and/or enzyme expression can coax a plastic response from the FADS2 enzyme – potentially yielding a much wider array of mono and polyunsaturated fatty acid isomers. For example, relative to the normal prostate BPH-1 cells, PC-3 cancer cells display a significant change in expression of SCD-1 and FADS2 (*cf*. Fig. 2D), which in turn has a large downstream effect on the isomeric complexity expressed within the mono- and polyunsaturated fatty acids of PC-3 compared to BPH-1 (*cf*. Fig. 5B). Furthermore, if substrate availability is altered in combination with these enzyme expression/activity differences, FADS2 (and possibly FADS1) plasticity allows for substrate acceptance usually observed in sebocytes (*e.g*., FADS2 Δ6-desaturation of 14:0, 16:0 and 18:0 to yield *n*-8, *n*-10 and *n*-12, respectively) or even changes to the enzyme-substrate complex to result in an alternative site of desaturation (*e.g*., FADS2 Δ8-desaturation of 18:0 to yield 18:1*n*-10) (*cf*. Fig. 4D). The metabolic similarity to sebum is an interesting point to ponder; sebum only contains these unusual FADS2 metabolites because sebocytes lack the expression of SCD-1.^27^ One reason for this may be because SCD-1 desaturation is an oxygen dependant reaction, meaning the oxygen-independent FADS2 can more finely regulate desaturation reactions for lipid homeostasis in the oxygen rich environment of the skin. Given the hypoxic conditions of some tumorous tissues, it is logical to think that the SCD-1 reaction would become limited^12^ and thereby cause the upregulation of FADS2 to allow adoption of primary desaturation behaviours alongside its role in polyunsaturation events. This would account for: the increase in FADS2 expression seen in the transcripts of cancer cell lines (Fig. 2 D), the unusual desaturation products observed (Fig. 2C and Fig. 5B), and the increase in polyunsaturated FA products observed (Fig. 5A). Similarly, under metabolic stress, SCD-1 and FADS1 also displayed plasticity towards substrate preference to allow for the formation of *n*-5, *n*-11 and *n*-13 FA families, further demonstrating cancers propensity to adapt.

As has been shown in previous studies into cancer and normal prostate cell lines, a strong distinction is created when observing the differences of FA 18:1 DB isomers from the hydrolysed FA pool.^18^ These differences pertain to a change in either substrate availability or substrate preference by the SCD-1 enzyme, which in turn causes a statistically significant increase of the FA 18:1*n*-9 isomeric fraction (relative to the FA 18:1*n*-7) within cancer cells. More recently, in a study of human breast cancer and adjacent non-tumorous tissues, it was shown that a small number of sum composition phospholipids displayed a minor but statistically significant change in abundance.^38^ The authors were then able to show that: (i) discerning the double bond location greatly increased the significance of the phenotypic distinction, and (ii) the phospholipids from the cancer tissues contained a much higher proportion of the 18:1*n*-9 fatty acyl chain compared to the FA 18:1*n*-7. Herein, we were able to show that the elucidation of double bond positions in the context of complex lipids served to further improve phenotypic distinction while simultaneously providing a rich fingerprint for metabolic activity in prostate cancer tissues. Furthermore, similar to the findings in breast cancer tissue, we were able to show that epithelial cancer cells in prostate tissue also contained higher proportions of *n*-9 in PC 34:1 (*cf*. Fig. 1A). This feature of aberrant SCD-1 behaviour however, was also marked by the absence of usual FADS2 polyunsaturated FA metabolites and perhaps indicates FADS2 plasticity with a change towards primary desaturation. The *n*-10 FA metabolites associated with this enzyme plasticity were not observed in the imaging of PC 34:1, but instead were found in other phospholipid classes suggesting that, similar to PCa cell lines, these unusual fatty acids are unevenly distributed among phospholipid subclasses. *n*-10 Phospholipids and other metabolites generated via FADS2 plasticity were detected in all clinical prostate specimens, however no linear relationship was observed between the presence of *n*-10 phospholipids and percentage tumour content. As all tissues examined (Fig. 1 A&C) were derived from radical prostatectomy specimens containing prostate cancer, it is not possible to disentangle if unusual FADS2 activity is a general feature of prostate epithelial cells or specific to unhealthy (*i.e*., cancer, field-cancerisation, or premalignant aberrations in histologically benign tissue) prostate cell function. Further experimentation and comparison to tumour-free prostate tissue derived from healthy prostate (obtained at cystoprostatectomy) and benign prostatic hyperplasia will provide further insight into this distinction and hence clinical utility.

From cell line studies, we were able to reveal that all FA metabolites formed from apocryphal desaturation activity were active substrates for elongation, and ELOVL2 was shown as a candidate in further diversifying the *n*-10 family. This is perhaps an ode to the canonical role of ELOVL2 in elongating FADS2-modified dietary fatty acyl metabolites,^1^ which may suggest co-localisation or co-activity between enzymes. Instead of these unusual FA metabolites being regarded by cellular machinery as malformed or unusable, they are functionalised through incorporation into known membrane and signalling phospholipid classes. As further indication towards a functional role, these unusual FA metabolites are found to be unevenly associated with certain phospholipid subclasses compared to others (*cf*. Fig. 3, supplementary Fig. S4A-D). Given that double bond location will likely affect inter-molecular forces and membrane fluidity,^3,5^ here it is speculated that the change in isomeric DB composition will impact independent organelle membrane fluidity unevenly. In turn, this could aid increased mitosis for cancer cell stemness^8^ or even hinder apoptotic processes, such as PS outer-leaflet exposure^14^ or peroxidation and ferroptosis.^39^ Similarly, the synthesis of entirely *de novo* synthesised polyunsaturated fatty acids, such as FA 20:4*n*-7, bear remarkable similarity to biologically active FA metabolites and are being incorporated into known signalling phospholipids. Consequently, this may disrupt homeostatic lipid signalling, or indeed, fulfil usual signalling roles in the absence of dietary FA uptake or other chemical environment changes. Although further work is required to specify the biological impact that unusual lipid unsaturation has on cells, the experimental workflows and findings presented throughout this body of work serve as a roadmap toward future discovery.

## Acknowledgements

This study was supported by the Movember Foundation and the Prostate Cancer Foundation of Australia through a Movember Revolutionary Team Award and the Australian Government Department of Health. Part of this work was conducted at the Translational Research Institute which is supported by a grant from the Australian Government. Some of data reported in this paper were obtained at the Central Analytical Research Facility operated by the Institute for Future Environments (QUT, Brisbane, Australia). Access to CARF is supported by generous funding from the Science and Engineering Faculty (QUT). This work was supported through funding from the Australian Research Council (DP150101715, LP180100238, DP190101486). A.P.B, R.M.A.H and S.R.E are grateful for funding from Interreg V EMR and the Netherlands Ministry of Economic Affairs within the “EURLIPIDS” project (EMR23). RSEY would like to acknowledge support through an Australian Government RTP scholarship and support through the award of a QUT Supervisor scholarship top-up.

## Author contributions

### Investigation and quantification

MCS cultured all cells and was responsible for cell counting, tracing and treatments and biological quality control. KDT was responsible for cell line qRT-PCR for mRNA data acquisition and assisted with cell maintenance. CLB conducted western blot analyses of FADS2 antibodies. EDW acquired human prostate tumour tissues, ensured ethical responsibilities were upheld and was responsible for sectioning, slide mounting and H&E staining of all tissue images displayed. APB prepared samples and acquired data for MALDI-MSI-OzID. RSEY was responsible for all remaining data acquisitions and analyses, including cell line lipid bi-phasic extraction, normalisation and quantification, tissue lipid extraction, mRNA expression data analysis, MALDI-MSI-OzID data analysis, complex lipidomics acquisition and analysis, GCMS FAMEs acquisitions and analysis, OzID double bond acquisition and analysis (for all positive/negative intact lipids from tissue and cell lines, lipid standards, AMPP fatty acid, carbon tracing lipids and gene silencing treatment lipids), AMPP derivatisation wet chemistry procedure and all statistical modelling throughout.

### Informal analysis

DLM, BLJP and SJB all contributed to mass spectrometry understanding. MSC and KDT both contributed and assisted with biological and biochemical understanding. EDW contributed to histopathology understanding of prostate tissue and cancer. APB SRE and RMAH all contributed with mass spectrometry imaging understanding. RG contributed to gas chromatography understanding.

### Resources

SJB, CCN, MCS, RMAH and SRE all provided resources to the experiments undertaken throughout.

### Supported the development of methods

BLJP, SJB, DLM supported OzID-MS instrumentation. VRN supported AMPP+ method development. RG supported GCMS FAMEs method development. BLJP, RG, RSEY supported automated complex lipidomics. SRE, APB supported MALDI-MSI-OzID. RSEY, DLM supported automated OzID.

### Conceptualised

The project was conceptualised by RSEY, SJB and MCS.

### Data visualisation

Data visualisation was performed by RSEY.

### Writing/editing

RESY wrote the first draft, SJB reviewed and all authors contributed to editing.

### Supervision/funding/PI

SJB and MCS supervised the direction of this project and SJB, BLJP, CCN, SRE and RMAH all provided project funding.

### Competing interest declaration

MALDI-MSI OzID technology was developed through an industry linkage supported by Waters Corporation and the Australian Research Council (LP180100238).

## Additional information

### Data Availability

The datasets generated during and/or analysed during the current study are available in the QUT Research Data Finder repository, https://researchdatafinder.qut.edu.au/display/n8324

### Correspondence

should be addressed to SJB and MCS

## Methods

### 1.0 Lipid nomenclature

Lipid nomenclature was based on previously defined shorthand naming systems that only state the level of molecular detail that is known.^1,2^ In short, the lipid category is defined by a two letter abbreviation (*e.g*., PC; phosphatidylcholine) followed by the number of carbons and double bonds in the fatty acids separated by a colon (*e.g*., a PC with 34 carbons and 2 double bonds; PC 34:2). If further analysis has been undertaken to reveal individual fatty acid composition but not the stereospecific number on the glycerol backbone (*i.e*., *sn-*position), this can be indicated with an underscore (*i.e*., PC 16:0_18:2). Instead, if *sn*-position is known fatty acids can be separated by a forward-slash (*i.e*., *sn-1/sn-2*; PC 16:0/18:2). For established double bond positions, the digit indicating the number of double bonds is directly followed by the *n*-number when referencing the number of carbons distal to the methyl terminus (*i.e*., PC 16:0/18:2*n*-6) or the bracketed Δ-number when referencing the carboxylate terminus (*i.e*., PC 16:0/18:2(*Δ9)*). It should be noted that a single double bond position given in polyunsaturated series implies methylene interruption of subsequent double bonds (*i.e*., PC 16:0/18:2*n*-6 = PC 16:0/18:2*n*-6,9).

### 2.0 Materials

200 µL glass vial-inserts, 2 mL and 4 mL glass vials and ultraclean Silicone/PTFE septa caps (Thermo Fisher, Waltham, MA, USA) were used for biology, analytical and wet-chemistry methods. LCMS grade methanol, Optima^®^ grade acetonitrile (ACN), N,N-dimethyl formamide (DMF) and water and HPLC grade methyl tert-butyl ether (MTBE) were obtained from Thermo Fisher used throughout. Hydrochloric acid (HCl; 37% in water), anhydrous sodium acetate (NaOAc), ammonium acetate (NH_4_OAc), trimethylsulfonium hydroxide (LiChropure™ TMSH, 0.25 M in MeOH), tetrabutylammonium hydroxide (TBAOH; 40 wt. % in water) and dibutylhydroxytoluene (BHT) solid were all obtained from Sigma Aldrich, Castle Hill, Australia. For MALDI-MS imaging chloroform (LC-MS grade), methanol (LC-MS grade), 2,5-dihydroxyacetophenone (DHA, ≥99.5%, Ultra pure), and anhydrous sodium acetate (NaOAc, ≥99%) were all purchased from Sigma Aldrich (Zwijndrecht, The Netherlands) and used without further purification.

### 3.0 Biological methods

#### 3.1 Tissue ethics and sectioning/mounting

Prostate tissues were collected with ethical approval of the St. Vincent’s Hospital Human Ethics Committee and in accordance with Australian National Health and Medical Research Council Guidelines. Tissues samples were collected from radical prostatectomy specimens by a pathologist, immediately snap frozen in liquid nitrogen, and stored at −80°C. Using a CM 1950 Cryostat (Leica Biosystems, Nussloch, Germany), tissue biopsies were sectioned at 10 µm thickness using a blade that was free from optimal cutting temperature (OCT) compound and fixed on standard glass slides (SuperFrost +, Menzel-Gläser, Braunschweig, Germany) using 10% neutral buffered formalin for 30 seconds prior to MALDI-MS imaging protocol (see method section 4.2) and the haematoxylin and eosin (H&E) staining protocol (see method section 3.2). Tissue slides for MALDI-MSI were placed into a sealed slide holder and purged with nitrogen gas before being stored on dry ice for inter-laboratory shipping.

#### 3.2 Haematoxylin and Eosin Staining

Tissue H&E staining took place after sectioning/mounting using an autostainer (Tissue-Tek Prisma, Sakura Finetek, Torrance, CA, USA) according to the following sequence: tissues were washed with water for 2 minutes before being exposed to haematoxylin (Harris Haematoxylin (PAH), Australian Biostain P/L, Traralgon, Australia) for 5 minutes. “Bluing” was achieved with water rinsing for 4 minutes and exposed for 10 seconds to ethanol (Chem Supply Gilman, Australia) acidified with 1% HCl before 5 minutes of further rinsing with water. Eosin staining (0.25% Eosin Y; certified C.C. # 45380, ProSciTech, Kirwan, Australia) took place for 2 minutes before 40 seconds of water rinsing. One 80% ethanol rinse followed by two 100% ethanol rinses then took place for 45 seconds, 30 seconds and 45 seconds, respectively. Triplicate xylene (Point Of Care Diagnostics, North Rock, Australia) washes were then conducted for 1 minute each. Coverslips (Tissue-Tek Glas, Sakura Finetek, Torrance, CA, USA) were then applied before imaging using a Pannoramic Digital slide scanner (3DHistech, Hungary) for histopathological analyses.

#### 3.3 Cell culturing

LNCaP (CVCL_0395), VCaP (CVCL_2235), PC-3 (CVCL_0035), DU145 (CVCL_0105) and RWPE-1 (CVCL_3791) cells were obtained from the American Type Cell Culture Collection (ATCC; Manassas, Virginia, USA), while BPH-1 (CVCL_1091) was gifted by P. J. Russell and J. Clements (Australian Prostate Cancer Centre-Queensland, Australia). All cell lines were cultured in Roswell Park Memorial Institute (RPMI) medium (Thermo Fisher, Waltham, MA, USA) supplemented with 5% foetal bovine serum (FBS, Invitrogen™, Waltham, MA, USA) and incubated at 37°C in 5% CO_2_. Medium was changed every 3 days, and cells were passaged at approximately 80% confluency by trypsinisation. Cell lines were authenticated using genotyping in March 2018 by Genomics Research Centre (Brisbane, Australia) and routinely tested to exclude mycoplasma infection. Cell number and viability was determined by trypan blue staining and using a TC20 Automated Cell Counter (Bio-Rad).

#### 3.4 Gene silencing by siRNA and enzyme inhibition

LNCaP cells were seeded at 1.2×10^5^ cells/well in 6-well plates. After 48 hours, cells were transfected with 10 nM siRNAs (Sigma-Aldrich) targeting SCD-1 (SASI_Hs01_00029617, SASI_Hs01_00181377, SASI_Hs01_00181371), FADS2 (SASI_Hs01_00029610, SASI_Hs01_00029612, SASI_Hs01_00029608), ELOV2 (SASI_Hs01_00018935), ELOVL3 (SASI_Hs01_00086480), and ELOVL6 (SASI_Hs01_00242071). Off-target effects were controlled using a scrambled siRNA control sequence (SIC001, Sigma-Aldrich) at 10 nM. Before forward transfection, growth medium was replaced with 1 mL serum free medium (RPMI 1640), and transfection solution using RNAiMAX lipofectamine reagent was prepared according to the manufacturer’s instructions (Thermo Fisher). Following six hours of transfection, FBS was added medium to a final concentration of 5%, and cells were grown for 72 hours. For direct inhibition of enzyme activity, cells were treated for 72 hours with the indicated concentrations of SC26196 (FADS2Δ6 inhibitor, Sigma-Aldrich) and 5-tetradecyloxy-2-furoic acid (TOFA; combination SCD-1Δ9 /ACC1 inhibitor, Sigma-Aldrich).

#### 3.5 ^13^C carbon tracing

For ^13^C tracing studies, LNCaP cells were chosen due to their heightened FADS2 expression. Cells were seeded and cultured under the conditions mentioned previously (*cf*. method section 3.3). After 48 hours of seeding, cells were switched to fresh media supplemented with either unlabelled (*i.e*., ^12^C_16_-palmitic acid) or labelled (*i.e*., ^13^C_16_-palmitic acid or ^13^C_18_-stearic acid) fatty acids conjugated to bovine serum albumin (BSA) at a final concentration of 20 µM. All fatty acids were purchased from Sigma Aldrich, Castle Hill, Australia. Cells were grown for a further 72 hours and washed twice with ice-cold phosphate buffered saline (PBS) before lipid extraction.

#### 3.6 RNA extraction and quantitative real-time polymerase chain reaction (qRT-PCR)

Cells seeded in 6 well plates were grown in 5% FBS to a confluency of 70% before RNA extraction using the RNEasy mini kit (Qiagen, Hilden, Germany) following the manufacturer’s instructions. RNA concentration was measured using a NanoDrop ND-1000 Spectrophotometer (Thermo Scientific, Waltham, MA, USA). 2 µg of total RNA was used to prepare cDNA with SensiFast cDNA synthesis kit (Bioline) according to the manufacturer’s instructions and diluted 1:6 with DNAse/RNAse-free water (Thermo Fisher, Waltham, MA, USA). qRT-PCR was performed with SYBR-Green Master Mix (Thermo Fisher Scientific, Waltham, MA, USA) using the ViiA-7 Real-Time PCR system (Applied Biosystems, Forster City, CA, USA). Determination of relative mRNA levels was calculated using the comparative ΔΔCt method,^*3*^ where expression levels were normalised relative to that of the housekeeping gene receptor-like protein 32 (RPL32) for each treatment and calculated as fold change relative to the expression levels of BPH-1 cells. All experiments were performed in triplicate and analysis and statistics were performed with Microsoft Excel. Primer sequences can be found in supplementary information (*cf*. supplementary Fig. S12).

#### 3.7 Western blot confirmation method

Cell seeding and gene silencing of FADS2 was carried out as described above. Protein extracts for Western blotting were generated from whole cell lysates prepared in lysis buffer [50 mM Tris, HCl pH 7.6, 150 mM sodium chloride, 1% Triton-X100, 0.5% sodium deoxycholate, 0.1% SDS, one cOmplete™ EDTA-free Protease Inhibitor Cocktail tablet (Roche) per 10 ml, phosphatase inhibitors sodium fluoride (30 mM), sodium pyrophosphate (20 mM), β-glycerophosphate (10 mM), and sodium orthovanadate (1 mM)]. Before lysis in 250 µL buffer on ice for 5 minutes, cells were washed twice with ice-cold PBS. Protein extracts were cleared by centrifugation for 10 minutes at 20,000 x g at 4°C and transferred into fresh tube. Protein concentration was measured using Pierce BCA Protein Assay kit according to manufacturer’s instructions (Thermo Fisher Scientific, Waltham, MA, USA). 20 µg of total protein/lane were separated by SDS-polyacrylamide gel electrophoresis (SDS-PAGE) using NuPAGETM 4-12% Bis-Tris SDS-PAGE Protein Gels (Thermo Fisher Scientific), and Western blot was completed using the Bolt Mini Blot Module (Thermo Fisher Scientific) according to the manufacturer’s instructions. After transfer and blocking of polyvinylidene fluoride (PVDF) membranes (Immobilon) in 5% BSA TBS Tween-20 buffer (Thermo Fisher Scientific), primary antibody directed against FADS2 (PA5-87765, Thermo Fisher Scientific) was applied overnight at 4ºC at a dilution of 1:1000 followed by probing with the appropriate Odyssey fluorophore-labelled secondary antibody and visualization on the LiCor^®^ Odyssey imaging system (LI-COR^®^ Biotechnology, NE, USA). Protein expression levels were quantified using Image Studio Lite (LI-COR^®^ Biotechnology), normalised relative to the indicated housekeeping protein, and expressed as fold-changes relative to the control treatment.

#### 3.8 Lipid extraction

Cell and homogenised tissue lipids were extracted using methods similar to those described by Matyash *et al*.^*4*^ and were quantifiable through the use of internal standards in the form of deuterated lipids (SPLASH Lipid-o-mix, Avanti Polar Lipids, Alabaster, USA) and an odd-chain fatty acid found to be not present within the samples (nonadecanoic acid, Sigma Aldrich, Munich, Germany). To minimise pipetting error, a stock internal standard solution was made in bulk using 720 μL MTBE (0.01% BHT), 40 μL SPLASH Lipid-o-mix^®^ and 20 μL nonadecanoic acid in MTBE (3.35mM) per 2 M cells. Cell pellets in 2 mL clear glass vials (∼2 M cells) were twice washed with PBS solution, before adding 220 μL of methanol and 780 μL of the prepared internal standard stock solution. Capped vials were vortexed for 20 s, before 1.5 h bench-top agitation. Phase separation was induced by adding 200 µL of aqueous ammonium acetate (150 mM) before samples were vortexed for 20 s and centrifuged for 5 min at 2,000 x g. The organic supernatant was pipetted off to a clean labelled 2 mL glass vial and stored at – 20°C before analyses.

#### 3.9 AMPP derivatisation

100 μL of the lipid extract was dried under nitrogen gas in 4 mL glass vials. Fatty acids were hydrolysed from lipids using 1:1 methanol:tetrabutylammonium hydroxide (40 wt. % in water) and heating at 85 °C for 2 h before allowing to cool to room temperature. 1.5 mL of water (Optima^®^) was added to each of the vials and acidified with 120 μL of 5 M aqueous hydrochloric acid to achieve a pH of 2. Biphasic extraction of the supernatant was then undertaken using two separate aliquots (1 mL) of n-hexane to optimise the recovery of fatty acids. The individual sample supernatant fractions were combined and dried under nitrogen to yield fatty acids. The obtained fatty acids were then functionalised with AMPP+ using an AMP+ Mass Spectrometry Kit (Cayman Chemical, Ann Arbor, MI). and following a similar method discussed by Bollinger et al.^*5*^ Briefly, 150 μL of 4:1 ACN:DMF was added to the hydrolysed lipid samples prepared above, followed by 10 μL of 1-ethyl-3-(3-dimethylaminopropyl)carbodiimide (EDC•HCl; 1 M in water), 20 μL of 1-hydroxybenzotriazole (5 mM in 99:1 ACN:DMF), and 20 μL of AMPP+ coupling reagent (15 mM in ACN). The resulting solution was sealed, vortexed for 1 min and heated at 65 °C for 30 min. After cooling to room temperature, the reaction mixtures were diluted with water (1 mL) and saturated aqueous NH_4_Cl (50 μL); mixtures were then twice extracted using MTBE (2×1 mL). The MTBE supernatants were combined and stored in sealed 2 mL vials at −20 °C before MS analysis. 5 μL of the Restek 37 mix of FAMEs and blanks were also derivatised through the above method and were used as experimental quality-controls.

### 4.0 Analytical methods

#### 4.1 Gas chromatography – pooled fatty acyl analysis

A reference standard of 38 fatty acid methyl ester standards was prepared by mixing a purchased 37 fatty acid methyl ester (FAME) standard (Restek, Bellefonte, PA, USA) and 450 μM methyl-nonadecanoate in MTBE (Sigma Aldrich) at a 1:9 ratio. Samples were prepared for analysis by mixing sample extracted lipids 5:1 with TMSH. Pooled batch quality controls (PBQC) and blanks were used throughout for quality control and data reliability.

Samples were analysed using a TQ8040 GC/MS (Shimadzu, Kyoto, Japan) with chromatographic separation being carried out through an RTX-2330 capillary column (cyanopropyl stationary phase, 60 m x 0.25 mm, 0.20 µm film thickness; Restek). GC/MS instrument conditions were then optimised for FAME separation (He carrier gas, column flow of 2.6 mL/min, 22:1 split ratio, 1 μL sample injections, injector temperature of 220°C, interface and ion source temperature of 250°C). To shorten the total experiment duration and assist with chromatographic separation, a 30 minute column oven temperature gradient was used (150°C initial temperature with a 10°C/min increase to 170°C, followed by a 2°C/min increase to 200°C and a further 1.3°C/min increase to 211°C where the temperature was held for the remaining 5 minutes of the experiment). Column eluents were then subject to 70 eV of source energy for electron ionisation and ions were detected by Q3 scan mode over a *m/z* 50-650 range.

#### 4.2 MALDI-MSI OzID for lipid double bond imaging

10 µm tissue sections mounted on standard glass slides were first thinly coated with 12 passes (45 mm spray height, 30 °C, 10 psi, 2 mm track spacing) of 100 mM sodium acetate (2:1 methanol/chloroform) via an HTX TM-Sprayer (HTXImaging, Chapel Hill, NC, USA). Sample slides were then coated with 2,5-DHA via sublimation (40 mg, 160 °C, 4 minutes) using an sublimator (HTXImaging, Chapel Hill, NC, USA). Coated sample slides were then loaded into a prototype µMALDI source^*6*^ (Nd:YAG laser operating at 1.5 kHz, producing 25 nJ pulses at 355 nm; Waters, Wilmslow, England) for sample desorption and ionisation. 50 µm^2^ pixels were sampled at a velocity of 2.0 mm/s with a 1.62 A laser diode current. Samples were analysed using a Synapt G2-*Si* HDMS mass spectrometer (Waters, Wilmslow, England) modified with a closed loop ozone generation system to deliver ozone (up to 18% w/w in oxygen) to the ion mobility gas inlet of the instrument, as described previously.^*7*^ Ozone was generated from oxygen feed gas (99.999% purity, Linde Gas Benelux BV, The Netherlands) using a high concentration ozone generator (TG-40 gen 2, Ozone Solutions, Hull, IA, USA) and the concentration measured online using a UV-absorption based ozone monitor (106-H, 2B Technologies, Boulder, CO). The mass spectrometer was operated in ion-mobility mode, resulting in a reaction time with ozone of ∼15 ms, corresponding to the ion-mobility drift time. The quadrupole mass filter was set to transmit m/z 782 (*i.e*., [PC 34:1+Na]^+^ and [PC 36:4+H]^+^) which was subsequently reacted with ozone in the ion mobility cell to yield OzID fragmentation. The resulting monoisotopic ions (precursors and products) were then mass analysed by time of flight (nominal resolution 15,000).

#### 4.3 Direct infusion ESI-OzID of lipid double bonds

The double bonds of intact glycerophospholipids were determined via mass spectrometry using a modified Orbitrap Elite high-resolution mass spectrometer (Thermo Scientific, Bremen, Germany) capable of ozone-induced dissociation. Briefly, ozone was produced via a high-concentration generator (Titan-30UHC Absolute Ozone, Edmonton, Canada) and was introduced into the helium buffer gas flow before conduction through to the high-pressure region of the linear ion trap (LIT).^*8*^ Similarly, a diverter valve was placed on the nitrogen gas inlet to the higher collisional dissociation (HCD) cell, and nitrogen was replaced with generated ozone gas.^*9*^

Operating in positive-ion mode for PC lipid OzID, cell line lipid extract samples were mixed 1:1 (v/v) with 500 µM methanolic sodium acetate solution and introduced to the mass spectrometer via a chip-based nano-electrospray source (TriVersa Nanomate, Advion, Ithaca, NY, USA) using 1.35 kV/0.35 psi spray parameters. Using the Thermo Xcalibur software package, a data independent acquisition sequence was created to perform sequential OzID (activation time (AT)= 2 s (HCD), collision energy (CE)= 1V) and CID/OzID (MS^2^: AT= 30 ms (LIT), normalised collision energy (NCE)= 40; MS^3^: AT= 1 s, NCE = 0) for 9 sodiated phosphatidylcholine precursor ion masses with a maximum injection time of 100 ms, isolation window of ±0.5 Da across a 175-1000 Da scan range. Included sodiated precursor ion *m/z* values were: 754.6, 780.6, 782.6, 804.6, 806.6, 808.6, 810.6, 830.6 and 832.6. This method was also used to assign double bonds within lipids that were labelled with stable-isotope fatty acids, with the only method modification being the precursor ion masses that were selected. For PCs labelled with heavy-palmitate or heavy-stearate three labelled and three unlabelled lipids were submitted for OzID and CID/OzID analysis (*m/z* 754, 770, 782, 798, 810 and 826; and *m/z* 754, 772, 782, 800, 810 and 826, respectively). The product ions from all fragmentation experiments were detected using the orbitrap mass analyser for high-resolution mass accuracy, allowing for unambiguous assignment of characteristic fragments to specific lipids and not isobaric lipids or isotopes. Intensity values obtained from OzID and CID/OzID mass spectrometric experiments were the average of 24 and 39 microscans, respectively.

Operating in negative-ion mode for PE, PS, PG and PI lipid OzID, cell line lipid extract samples were mixed 1:1 (v/v) with 5 mM methanolic ammonium acetate solution and introduced to the mass spectrometer via a TriVersa Nanomate, set to use −1.35 kV/0.35 psi spray parameters. Using the Thermo Xcalibur software package, 4 separate data independent acquisition sequences were created for each of the phospholipid subclasses. For each method, sequential OzID (AT= 2.5 s (HCD), CE = 1V) and CID (MS^2^: AT= 5 ms (LIT), NCE= 33-39) was performed for 9 deprotonated lipid precursor masses (totalling 36 phospholipids) with a maximum injection time of 100 ms, isolation window of ±0.5 Da across a scan range of 200-1000 Da. Included deprotonated precursor ion *m/z* values were: (for PE) 688.5, 716.5, 742.5, 744.5, 782.5,766.5, 768.5, 770.5 and 772.5; (for PS) 732.5, 760.5, 786.5, 788.5, 806.5, 810.5, 812.5, 814.5 and 816.5; (for PG) 719.5, 747.5, 773.5, 775.5, 793.5, 797.5, 799.5, 801.5 and 803.5; and (for PI) 807.5, 835.5, 861.5, 863.5, 885.5, 887.5, 889.5, 891.5 and 915.5. The product ions from all fragmentation experiments were detected using the orbitrap mass analyser for high-resolution mass accuracy (120,000 FWHM at 400 *m/z*), allowing for unambiguous assignment of characteristic fragments to specific lipids and not isobaric lipids or isotopes. Intensity values obtained from OzID and CID mass spectrometric experiments were averaged across 16 and 77 microscans individual scans, respectively.

Using methods similar to those developed for cell line extracts, OzID was also performed on homogenised tissue lipid extracts. Due to the small quantity of tissue from which lipids were extracted, extracts were first dried under nitrogen gas and reconstituted to ¼ the volume (4-fold increase in concentration). Samples were reconstituted in either methanolic sodium acetate (500 µM) for positive-ion mode or methanolic ammonium acetate (5 mM) for negative-ion mode. Using the parameters described previously for positive-ion mode PC acquisitions, OzID and CID/OzID was performed on three PC lipids (*m/z* 754.5, 782.6 and 810.6). Resulting OzID and CID/OzID data was detected using the orbitrap mass analyser (120,000 FWHM at 400 *m/z*) and ion intensity values were averaged across 11 and 23 microscans, respectively. Given the low sample concentration and the decrease in ion detection efficiency inherent to negative-ion mode mass spectrometry, the aforementioned method to obtain OzID double bond data for PE, PS, PG and PI lipids was modified. A data independent method was created to sequentially acquire OzID (AT=10 s; NCE=19-23) for the 32:1, 34:1 and 36:1 phospholipid compositions (12 total phospholipids) using a maximum injection time of 100 ms, isolation window of ±0.5 Da across a scan range of 200-1000 Da. Included *m/z* values were: 688.5, 716.5, 719.5, 732.5, 744.5, 747.5, 760.5, 775.5, 788.5, 807.5, 835.5 and 863.5. To further improve detection of low intensity signals, product ion fragments were analysed in the high-pressure region of the dual linear ion trap. Low resolution ion fragments were compared against standards run under the same conditions to improve reliability of assignments. Intensity values obtained from OzID experiments were averaged across 9 microscans.

#### 4.4 Direct infusion ESI-OzID of pooled fatty acid double bonds (AMPP+ derivatization)

Hydrolysed lipid extracts (including cell line lipids, LNCaP siRNA experiments and the fatty acid methyl ester standard mix) were derivatised with AMPP+ as described above (*cf*. method section 3.9) and introduced via chip-based nano-electrospray using 1.90 kV/0.5 psi spray parameters. Using the Thermo Xcalibur software package, data independent acquisition sequences were created to perform ozonolysis within the linear ion-trap (AT=5 s, NCE=25) for 24 fatty acid precursor ion masses with a maximum injection time of 100 ms, isolation window of ±0.5 Da and scan range between 105-600 Da. Included precursor ion masses for AMPP+ derivatised FAs were: 14:1 (*m/z* 393.5), 14:2 (*m/z* 391.5), 15:1 (*m/z* 407.5), 16:1 (*m/z* 421.5), 16:2 (*m/z* 419.5), 16:3 (*m/z* 417.5), 17:1 (*m/z* 435.5), 18:1 (*m/z* 449.5), 18:2 (*m/z* 447.5), 18:3 (*m/z* 445.5), 19:1 (*m/z* 463.5), 20:1 (*m/z* 477.5), 20:2 (*m/z* 475.5), 20:3 (*m/z* 473.5), 20:4 (*m/z* 471.5), 20:5 (*m/z* 469.5), 20:6 (*m/z* 467.5), 22:1 (*m/z* 505.5), 22:2 (*m/z* 503.5), 22:3 (*m/z* 501.5), 22:4 (*m/z* 499.5), 22:5 (*m/z* 497.5), 22:6 (*m/z* 495.5) and 24:1 (*m/z* 533.5)). Subsequent OzID product ions were then transferred through to the orbitrap mass analyser for unambiguous assignment by high-resolution mass detection (mass resolution 120,000 (FWHM) at *m/z* 400). Ion intensity values were averaged across 11 microscans.

#### 4.5 Conventional lipidomics for phospholipid profiles

Lipid extracts from cell lines were run through an automated lipidomics workflow using an LC-20A HPLC (Shimadzu, Kyoto, Japan) set to deliver 100 uL sample loop-injections into a mobile phase of 5 mM methanolic ammonium acetate flowing at 15 µL/min. The sample column and column oven were bypassed with Viper PEEKsil (50 µm, Thermo Fisher, Waltham, MA, USA) to maintain instrument back pressure limits. Sample lipids were then directly infused through the electrospray ionisation source of a QTRAP 6500 hybrid triple quadrupole/LIT mass spectrometer (SCIEX, Concord, ON, Canada) using a spray voltage of 5 kV, a source temperature of 150 °C and both source gasses set to 15 (arb.). Various precursor ion and neutral loss scans were employed to confirm lipid head group, with the detected *m/z* being indicative of summed-fatty-acyl composition. (PC: PIS *m/z* 184.2, CE: 39V; PE: NL *m/z* 141.1, CE: 29V; PS: NL *m/z* 185.1, CE: 29V; PG: NL *m/z* 189.1, CE: 29V; PI: NL *m/z* 275.1, CE: 29V; ChE: NL *m/z* 259.1, CE: 29V). Instrument blanks were run through-out to ensure no sample carry-over and pooled batch quality controls (PBQC) were used to gauge instrument performance over the duration of the experiment.

### 5.0 Data processing

#### 5.1 Gas chromatography – pooled fatty acyl analysis

Tabulated ion intensity data from the *m/z* 55 extracted ion chromatogram (XIC) was extracted from the data files using the native Shimadzu Post-run analysis software. The *m/z* 55 XIC was chosen due to the enhanced detection of monounsaturated fatty acids. To ensure detection biasing was minimal, 3-point calibration curved were created with the reference standard to obtain molar correction factors for all fatty acid species. Using a Python script that was developed inhouse, chromatographic peaks were compared with a temporal tolerance of 0.013 min against the reference standard for fatty acyl species assignment. For quantification, the slope of the peak and the maximum height (min. threshold: 0.01% of total ions) was used to fit a Gaussian distribution, which was subsequently integrated and normalised to the methyl-nonadecanoate internal standard. Previous analysis of the samples without an included internal standard revealed no detectable methyl-nonadecanoate and hence all integrated chromatographic signal was attributed to the internal standard. Pandas DataFrames were then created and exported to .csv format where Microsoft Excel was then used remaining for cell count normalisation, internal standard concentration factoring, statistical analysis and graphing.

#### 5.2 MALDI-MSI OzID for lipid double bond imaging

HDI software (Version 1.4, Waters, Wilmslow, England) was used for data processing and image creation by integrating positional files obtained from the µMALDI source with the Waters .raw files from the Synapt G2-*Si*. The top 100 most intense fragment ion peaks were extracted from the raw/un-normalised spectra across a mass range of 200-1000 *m/z* with an isolation window of 0.02 Da and mass resolution set to maximum (20,000). Characteristic ions for the OzID aldehyde and Criegee ions were identified and summed to show isomeric distribution of PC 34:1. Similarly for PC 36:4*n*-6, aldehyde and Criegee ions from each double bond position were summed to show the distribution across tissue. Because quantitation was not the focus of the imaging analysis, maximum gradient intensity values were individually set to assist visualisation and contrast of lipid distributions. Hence, the maximum gradient values vary for each lipid species and can be observed in the linear colour scale bars of supplementary Fig. S1A-B. Images were smoothed using linear interpolation and composite images were created using the in-built “Add” data blending mode, which blends co-localised colours together.

#### 5.3 Direct infusion ESI-OzID of lipid double bonds and pooled fatty acid double bonds (AMPP+)

Averaged mass spectral data were extracted from the Thermo .raw files using Thermo Xcalibur Qual Browser (Version 3.0.63, Thermo Scientific, Bremen, Germany). Data tables were imported to Microsoft Excel where the data was normalised to the total ion count (TIC) before product ions were located and quantified. For high resolution mass spectrometry, a minimum peak intensity threshold of 0.013% of the TIC (∼10:1 signal-to-noise) was put in place, and a *m/z* Δppm of 6 was used for product ion assignment validation. Translation to double bond assignments was made according to previously published tables for OzID aldehyde and Criegee product ion neutral loss masses.^*8*^ OzID aldehyde and Criegee ion intensities were summed and represented as a fraction of the all isomer related OzID product ion signals (*i.e*., fractional distribution). This method of representation provides a further degree of normalisation for comparison, making the fractions reflective of changes in the molar concentration of isomers.

#### 5.4 Conventional lipidomics for phospholipid profiles

Lipidview^®^ (Version 1.3 beta, SCIEX, Concord, ON, Canada) was used for data processing of Sciex data files obtained from the QTRAP 6500. Lipid assignments were based on the software lipid tables and shortlisted to include even-chain lipids with 0-6 double bonds. Odd-chain/ether-lipid data was obtained but was not included in this study due to the ambiguity in assigning isobars in low resolution mass spectrometry. Isotope correction factors were applied and MS peaks were ratioed to the isotope corrected internal standard included in each scan type. The inclusion of deuterated and odd chain fatty acids within the internal standard lipids sufficiently mass shifted internal standards away from any biological lipids, therefore allowing accurate and reliable peak intensity measurements to be discerned. Data tables were extracted from Lipidview and imported to Microsoft Excel for cell count normalisation, internal standard concentration factoring, statistical analysis and graphing.

#### 5.5 Statistics and error analysis

The 95% confidence interval was used for error analysis on column-charts throughout and was calculated using Microsoft Excel. Box-plots display conventionally accepted values for the data minimum, 1^st^ quartile, median, 3^rd^ quartile and maximum. For Fig. 1B-C, the mean and variance were calculated using Microsoft Excel to establish an independent one-tailed Welsh’s t-test and t-values were translated to statistical significance via relevant degrees of freedom and critical t-value tables. Remaining heatmaps and correlation matrices including statistical significance were calculated using R x64 3.6.1 packages and built in functions using a Pearson product-moment correlation (*i.e*., PerformanceAnalytics, prcomp(), Hmisc() and corrplot()).

## Notes

https://researchdatafinder.qut.edu.au/display/n8324

## References

1 Guillou, H. et al. The key roles of elongases and desaturases in mammalian fatty acid metabolism: Insights from transgenic mice. Prog. Lipid Res. 49, 186–199, (2010).

2 Vriens, K. et al. Evidence for an alternative fatty acid desaturation pathway increasing cancer plasticity. Nature 566, 403–406, (2019).

3 Scanferlato, R. et al. Hexadecenoic Fatty Acid Positional Isomers and De Novo PUFA Synthesis in Colon Cancer Cells. Int. J. Mol. Sci. 20, 832, (2019).

4 Lorent, J. H. et al. Plasma membranes are asymmetric in lipid unsaturation, packing and protein shape. Nat. Chem. Biol., (2020).

5 Renne, M. F. & de Kroon, A. I. P. M. The role of phospholipid molecular species in determining the physical properties of yeast membranes. FEBS Lett. 592, 1330–1345, (2018).

6 Beloribi-Djefaflia, S., Vasseur, S. & Guillaumond, F. Lipid metabolic reprogramming in cancer cells. Oncogenesis 5, e189, (2016).

7 Epand, R. Features of the Phosphatidylinositol Cycle and its Role in Signal Transduction. J. Membr. Biol. 250, 353–366, (2017).

8 Li, J. et al. Lipid Desaturation Is a Metabolic Marker and Therapeutic Target of Ovarian Cancer Stem Cells. Cell Stem Cell 20, 303-314.e305, (2017).

9 Currie, E., Schulze, A., Zechner, R., Walther, T. C. & Farese, R. V. Cellular Fatty Acid Metabolism and Cancer. Cell Metab. 18, 153–161, (2013).

10 Zadra, G., Photopoulos, C. & Loda, M. The fat side of prostate cancer. BBA-Mol. Cell Biol. L. 1831, 1518–1532, (2013).

11 Biswas, S., Lunec, J. & Bartlett, K. Non-glucose metabolism in cancer cells—is it all in the fat? Cancer Metastasis Rev. 31, 689–698, (2012).

12 Röhrig, F. & Schulze, A. The multifaceted roles of fatty acid synthesis in cancer. Nat. Rev. Cancer 16, (2016).

13 Almut, S. & Adrian, L. H. How cancer metabolism is tuned for proliferation and vulnerable to disruption. Nature 491, 364, (2012).

14 Bratton, D. L. et al. Appearance of phosphatidylserine on apoptotic cells requires calcium-mediated nonspecific flip-flop and is enhanced by loss of the aminophospholipid translocase. J. Biol. Chem. 272, 26159, (1997).

15 Hsu, L.-C. et al. Evaluation of the Anti-Inflammatory Activities of 5,8,11-<i>cis</i>-Eicosatrienoic Acid. Food Sci. Nutr. Vol.04No.09, 7, (2013).

16 Fritz, V. et al. Abrogation of de novo lipogenesis by stearoyl-CoA desaturase 1 inhibition interferes with oncogenic signaling and blocks prostate cancer progression in mice. Mol. Cancer Ther. 9, 1740, (2010).

17 Tamura, K. et al. Novel lipogenic enzyme ELOVL7 is involved in prostate cancer growth through saturated long-chain fatty acid metabolism. Cancer Res. 69, 8133, (2009).

18 Ma, X. X. et al. Photochemical Tagging for Quantitation of Unsaturated Fatty Acids by Mass Spectrometry. Anal. Chem. 88, 8931–8935, (2016).

19 Nicolaides, N. Skin Lipids: Their Biochemical Uniqueness. Science 186, 19–26, (1974).

20 Ma, X. et al. Identification and quantitation of lipid C=C location isomers: A shotgun lipidomics approach enabled by photochemical reaction. PNAS 113, 2573–2578, (2016).

21 Thomas, M. C. et al. Ozone-induced dissociation: Elucidation of double bond position within mass-selected lipid ions. Anal. Chem. 80, 303–311, (2008).

22 Poad, B. L. J. et al. Ozone-induced dissociation on a modified tandem linear ion-trap: Observations of different reactivity for isomeric lipids. J. Am. Soc. Mass Spectrom. 21, 1989–1999, (2010).

23 Ellis, S. R. et al. More from less: high-throughput dual polarity lipid imaging of biological tissues. Analyst 141, 3832–3841, (2016).

24 Cocco, L., Follo, M., Manzoli, L. & Suh, P. G. Phosphoinositide-specific phospholipase C in health and disease. J. Lipid Res. 56, 1853–1860, (2015).

25 Williams, D. S. et al. Correction: Nonsense Mediated Decay Resistant Mutations Are a Source of Expressed Mutant Proteins in Colon Cancer Cell Lines with Microsatellite Instability. PLoS One 6, (2011).

26 Enoch, H. G., Catalá, A. & Strittmatter, P. Mechanism of rat liver microsomal stearyl-CoA desaturase. Studies of the substrate specificity, enzyme-substrate interactions, and the function of lipid. J. Biol. Chem. 251, 5095–5103, (1976).

27 Lan, G., Joel, S. G., Charleen, H., Kurt, S. & Stephen, M. P. Identification of the Δ-6 Desaturase of Human Sebaceous Glands: Expression and Enzyme Activity. J. Invest. Dermatol. 120, 707, (2003).

28 Moon, Y. A., Shah, N. A., Mohapatra, S., Warrington, J. A. & Horton, J. D. Identification of a mammalian long chain fatty acyl elongase regulated by sterol regulatory element-binding proteins. J. Biol. Chem. 276, 45358, (2001).

29 Jakobsson, A., Westerberg, R. & Jacobsson, A. Fatty acid elongases in mammals: Their regulation and roles in metabolism. Prog. Lipid Res. 45, 237–249, (2006).

30 Green, C. D., Ozguden-Akkoc, C. G., Wang, Y., Jump, D. B. & Olson, L. K. Role of fatty acid elongases in determination of de novo synthesized monounsaturated fatty acid species. J. Lipid Res. 51, 1871, (2010).

31 Obukowicz, M. G. et al. Novel, selective Δ6 or Δ5 fatty acid desaturase inhibitors as antiinflammatory agents in mice. J. Pharmacol. Exp. Ther. 287, 157–166, (1998).

32 Pauter, A. M. et al. Elovl2 ablation demonstrates that systemic DHA is endogenously produced and is essential for lipid homeostasis in mice S. J. Lipid Res. 55, 718–728, (2014).

33 Mason, P. et al. SCD1 Inhibition Causes Cancer Cell Death by Depleting Mono-Unsaturated Fatty Acids. PLoS One 7, e33823, (2012).

34 Bai, Y. et al. X-ray structure of a mammalian stearoyl-CoA desaturase. Nature 524, 252, (2015).

35 Wallis, J. G. & Browse, J. The Δ8-Desaturase ofEuglena gracilis:An Alternate Pathway for Synthesis of 20-Carbon Polyunsaturated Fatty Acids. Arch. Biochem. Biophys. 365, 307–316, (1999).

36 Poad, B. L. J. et al. Combining Charge-Switch Derivatization with Ozone-Induced Dissociation for Fatty Acid Analysis. J. Am. Soc. Mass Spectrom. 30, 2135, (2019).

37 Marquardt, A., Stöhr, H., White, K. & Weber, B. H. F. cDNA Cloning, Genomic Structure, and Chromosomal Localization of Three Members of the Human Fatty Acid Desaturase Family. Genomics 66, 175–183, (2000).

38 Wenpeng, Z. et al. Online photochemical derivatization enables comprehensive mass spectrometric analysis of unsaturated phospholipid isomers. Nat. Commun. 10, 1–9, (2019).

39 Tousignant, K. D. et al. Therapy-induced lipid uptake and remodeling underpin ferroptosis hypersensitivity in prostate cancer. bioRxiv (preprint), 2020.2001.2008.899609, (2020).

## Methods References

Liebisch, G. et al. Shorthand notation for lipid structures derived from mass spectrometry. J. Lipid Res. 54, 1523–1530, (2013).

Fahy, E. et al. Update of the LIPID MAPS comprehensive classification system for lipids. J. Lipid Res. 50, S9, (2009).

Schmittgen, T. D. & Livak, K. J. Analyzing real-time PCR data by the comparative CT method. Nat. Protoc. 3, 1101, (2008).

Matyash, V., Liebisch, G., Kurzchalia, T. V., Shevchenko, A. & Schwudke, D. Lipid extraction by methyl-tert-butyl ether for high-throughput lipidomics. J. Lipid Res. 49, 1137–1146, (2008).

Bollinger, J. G. et al. Improved Sensitivity Mass Spectrometric Detection of Eicosanoids by Charge Reversal Derivatization. Anal. Chem. 82, 6790–6796, (2010).

Barré, F. et al. Faster raster matrix-assisted laser desorption/ionization mass spectrometry imaging of lipids at high lateral resolution. Int. J. Mass Spectrom. 437, 38–48, (2019).

Poad, B. L. J. et al. High-Pressure Ozone-Induced Dissociation for Lipid Structure Elucidation on Fast Chromatographic Timescales. Anal. Chem. 89, 4223, (2017).

Paine, M. R. L. et al. Mass Spectrometry Imaging with Isomeric Resolution Enabled by Ozone-Induced Dissociation. Angew. Chem. Int. Ed. 57, 10530, (2018).

Marshall, D. L. et al. Mapping Unsaturation in Human Plasma Lipids by Data-Independent Ozone-Induced Dissociation. J. Am. Soc. Mass Spectrom., (2019).

