## Supplementary Information for "Metabolic plasticity in cancer activates apocryphal pathways for lipid desaturation"

* Corresponding authors

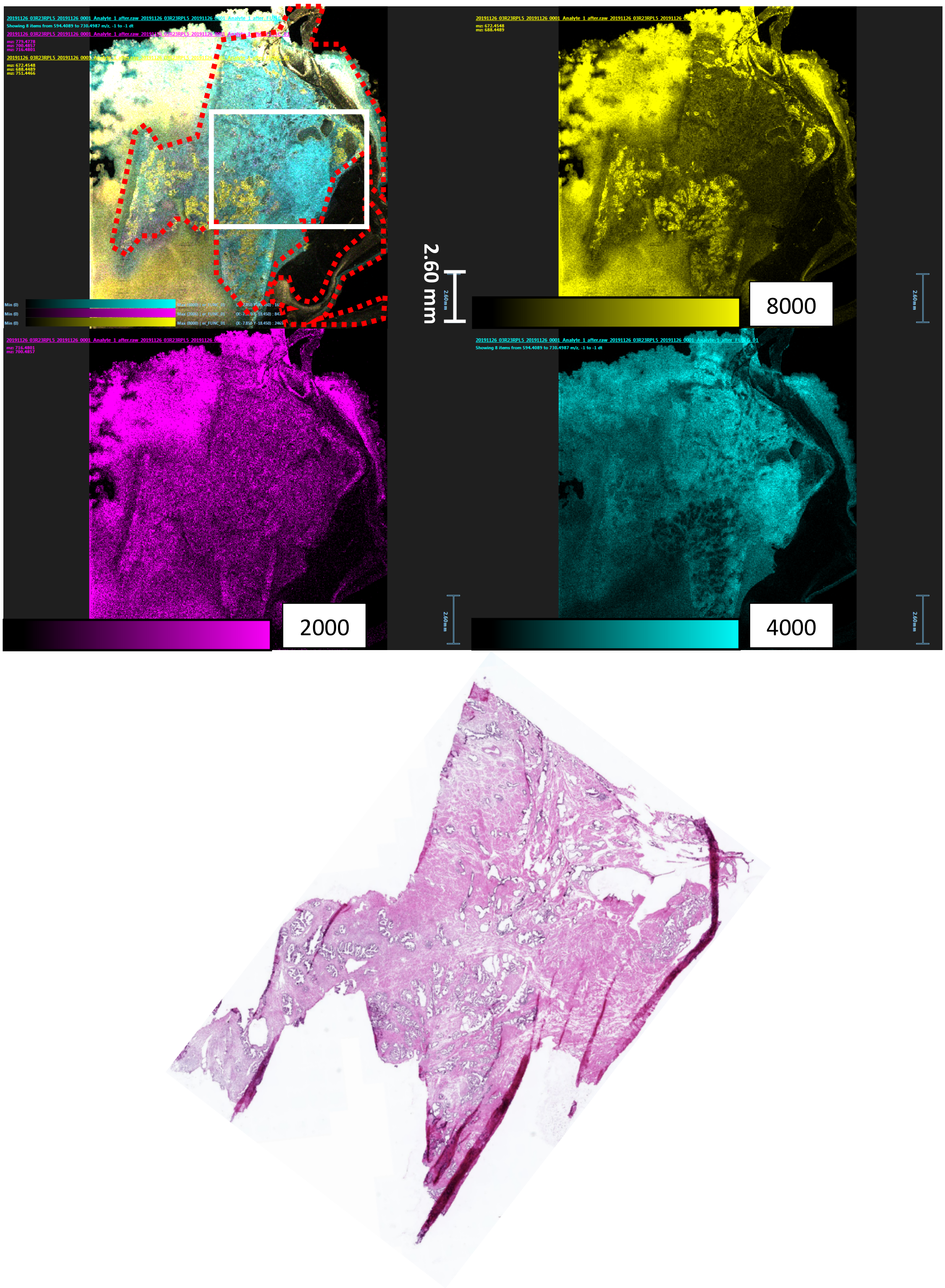

**Supplementary Fig. S1A:** Unmodified whole tissue images from MALDI-MSI OzID and H&E staining for the human prostate tissue section displayed in Fig. 1A first row. (From left to right, top to bottom) Composite image of the 3 individual lipid species from MALDI-MSI-OzID, PC 34:1*n*-9 (yellow), PC 34:1*n*-7, PC 36:4*n*-6, H&E stain of the adjacent tissue section. Tissue margin is represented with dotted red line, solid white line magnification area for Fig. 1A and max intensity counts are displayed next to colour scale bars. Lipid mobilisation to off-tissue can be observed and possibly caused by ‘smearing’ of the tissue sample during slide mounting. Mobilisation appears homogenous for all monitored lipids and therefore is only believed to minimally impact on-tissue analysis (cf. supplementary Fig. S1C). To confirm the lipid mobilisation hypothesis, an additional bio-replicate MALDI-MSI OzID image from an adjacent section can be found in supplementary Fig.1D.

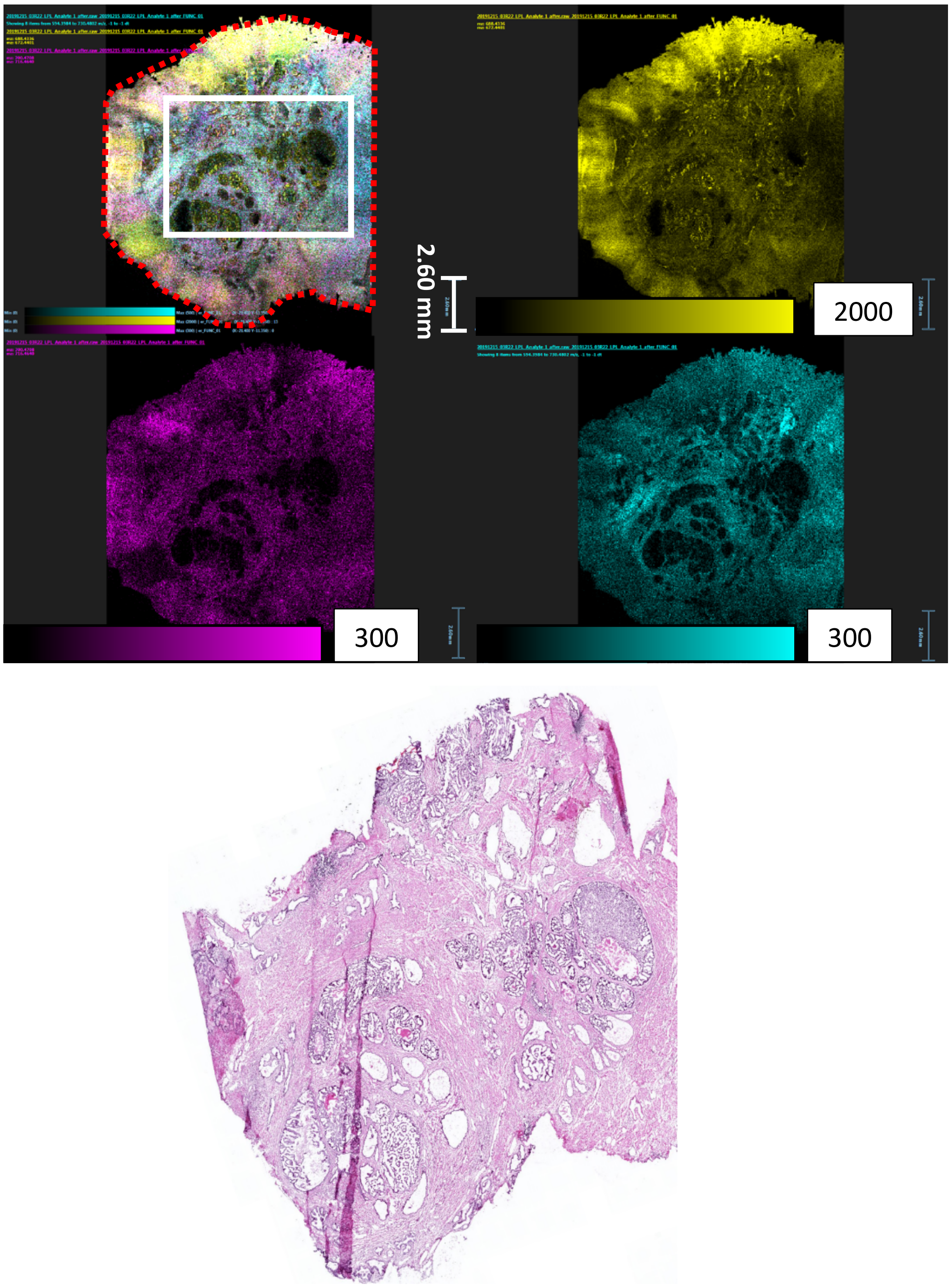

**Supplementary Fig. S1B:** Unmodified whole tissue images from MALDI-MSI OzID and H&E staining for the human prostate tissue section displayed in Fig. 1A second row. (From left to right, top to bottom) Composite image of the 3 individual lipid species from MALDI-MSI-OzID, PC 34:1*n*-9 (yellow), PC 34:1*n*-7, PC 36:4*n*-6, H&E stain of the adjacent tissue section. Tissue margin represented with dotted red line, solid white line magnification area for Fig. 1A and max intensity counts are displayed next to colour scale bars. Minimal mobilisation can be observed within this sample section, however H&E tissue shape appears skewed due to section folding.

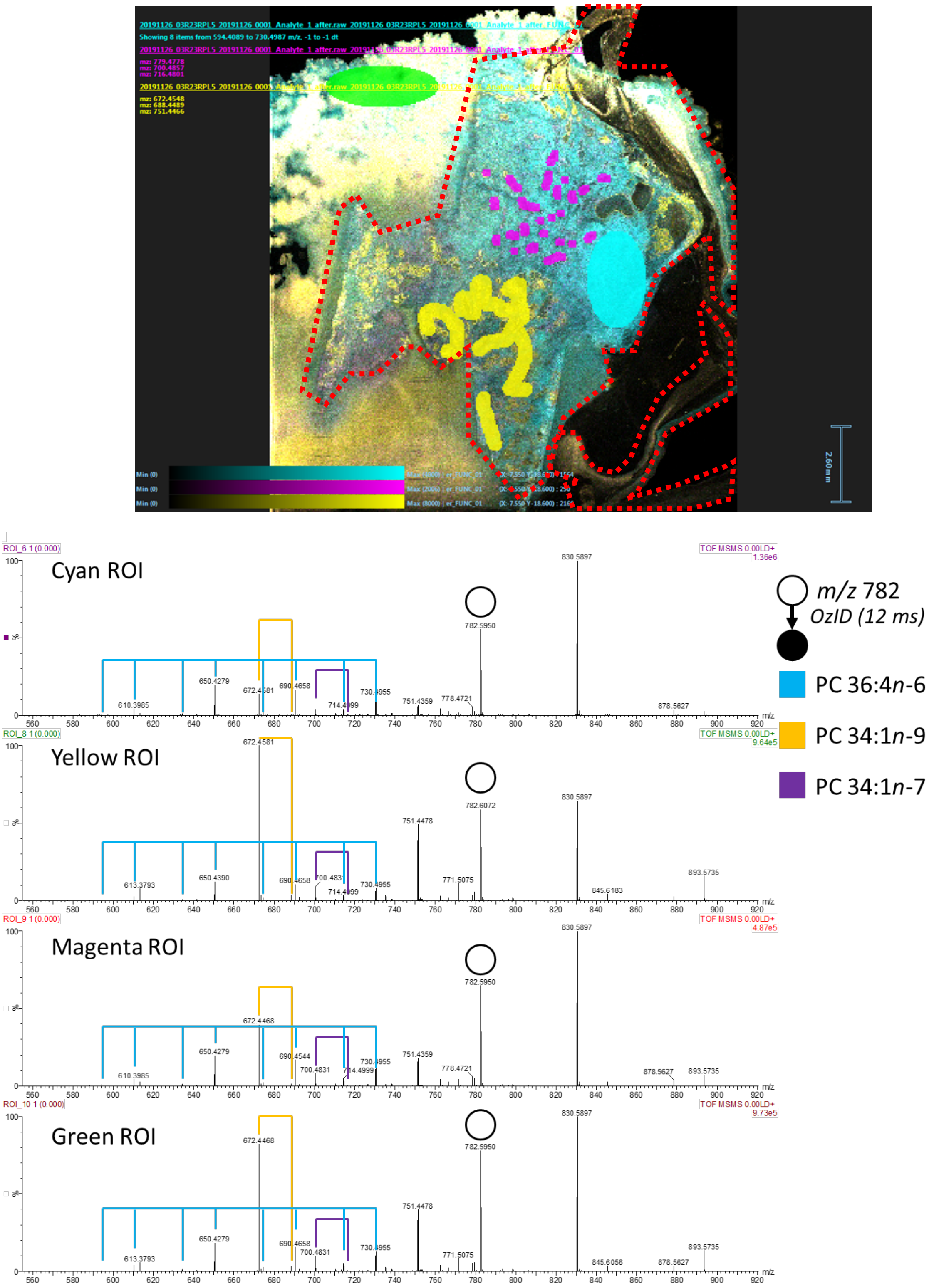

**Supplementary Fig. S1C:** MALDI-MSI OzID derived images displaying lipid distribution across tissue for human prostate lobe section and mass spectra from indicated regions of interest (ROI) indicated as colour block overlays. Product ions within the mass spectra confirm that there is homogenous mobilisation to off-tissue (green ROI) of the three monitored ions and distribution differences within the on-tissue ROIs (cyan, yellow, magenta).

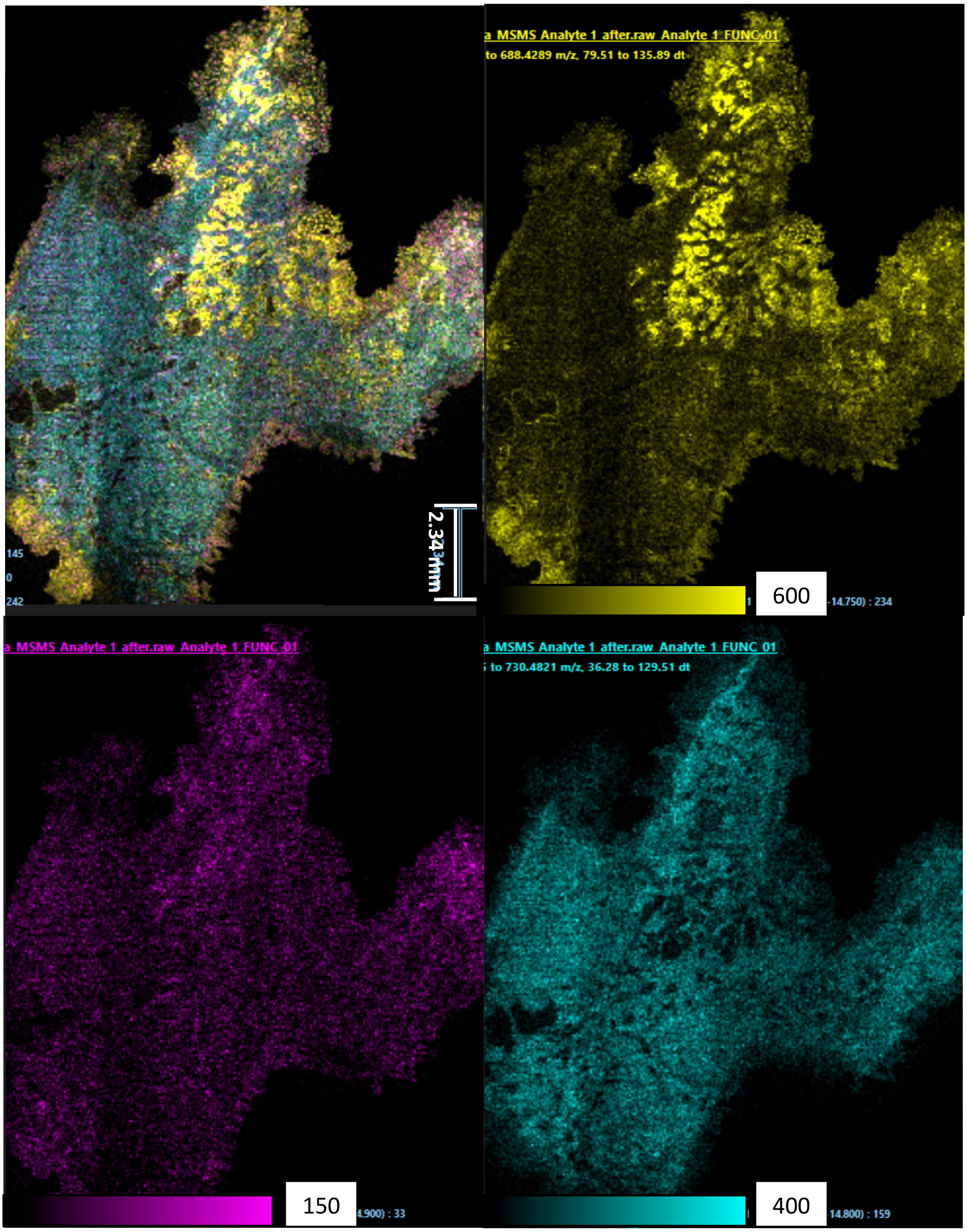

**Supplementary Fig. S1D:** Unmodified whole tissue images from MALDI-MSI OzID for a bio-replicate section of the human prostate tissue displayed in supplementary Fig. 1A (images approximately 180° rotated). (From left to right, top to bottom) Composite image of the 3 individual lipid species from MALDI-MSI OzID, PC 34:1*n*-9 (yellow), PC 34:1*n*-7, PC 36:4*n*-6, H&E stain of the adjacent tissue section. Max intensity counts are displayed next to colour scale bars. No lipid mobilisation to off-tissue can be observed within this replicate which may point to confirmation of the slide mount 'smearing' hypothesis observed in supplementary Fig. 1A.

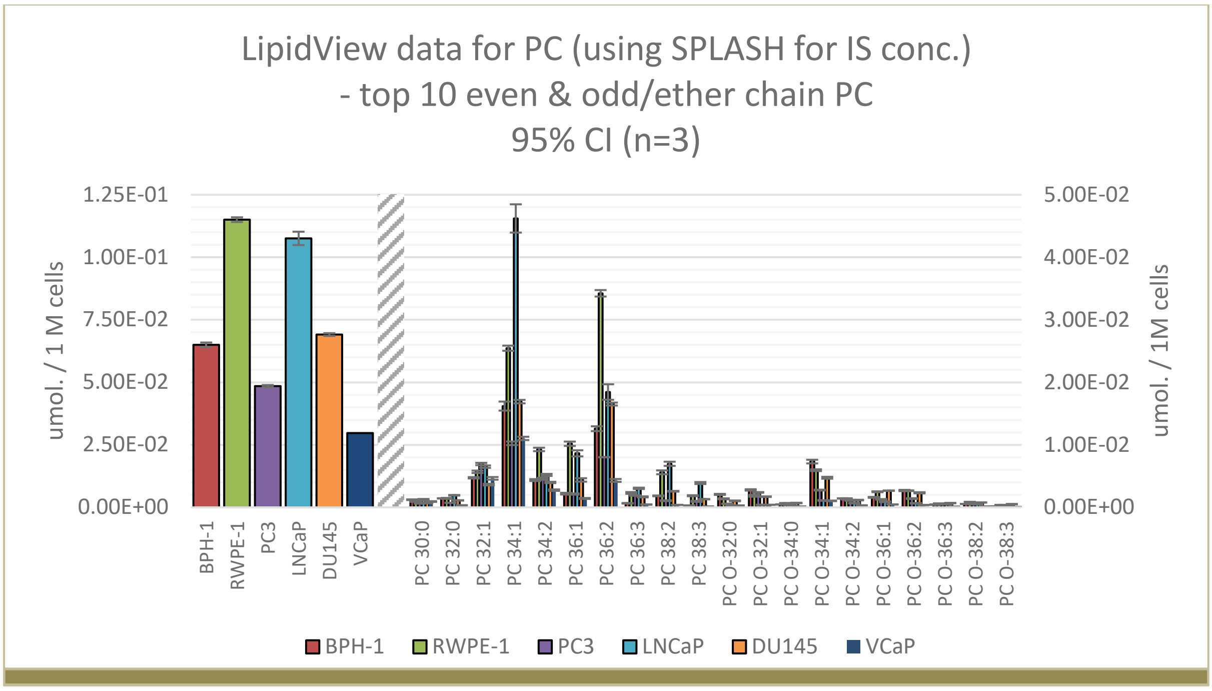

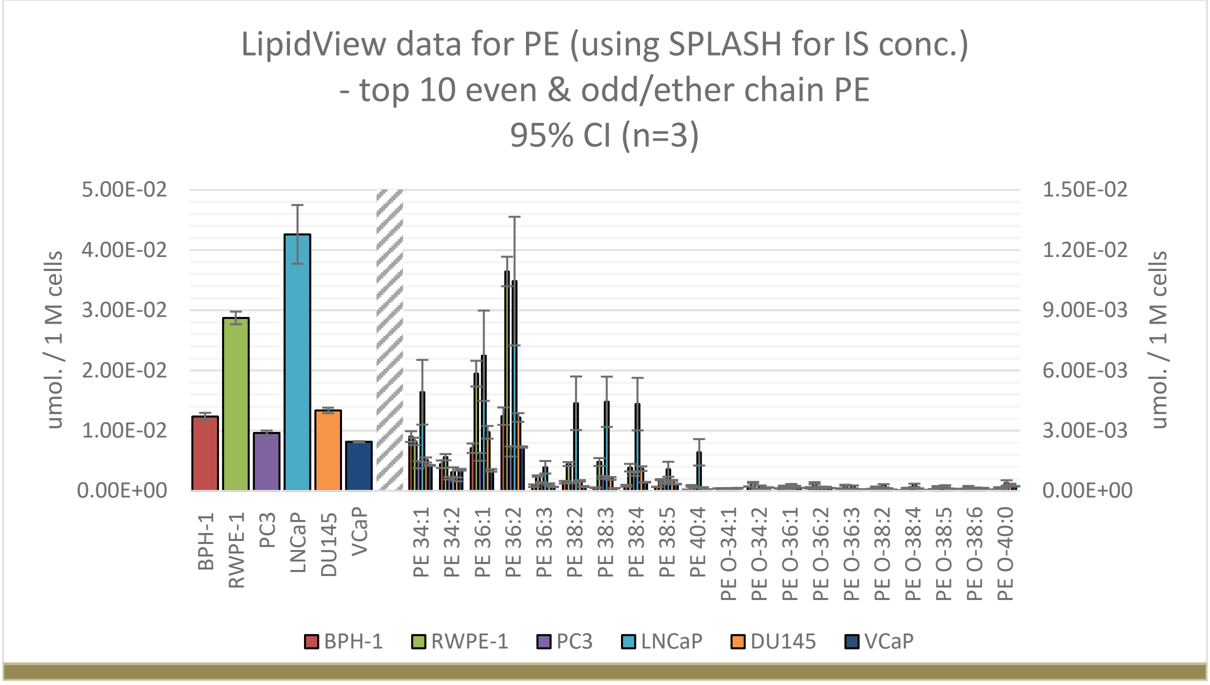

**Supplementary Fig. S2A:** Quantitative complex lipidomics profile from PCa cell lines for: (top) phosphatidylcholine (PC) and (bottom) phosphatidylethanolamine (PE). Left panels display total subclass lipid abundance and right panels display the top 10 most abundant even chain fatty acyl sum compositions and top 10 most abundant odd/ether chain fatty acyl sum compositions. (n=3; 95% confidence interval error)

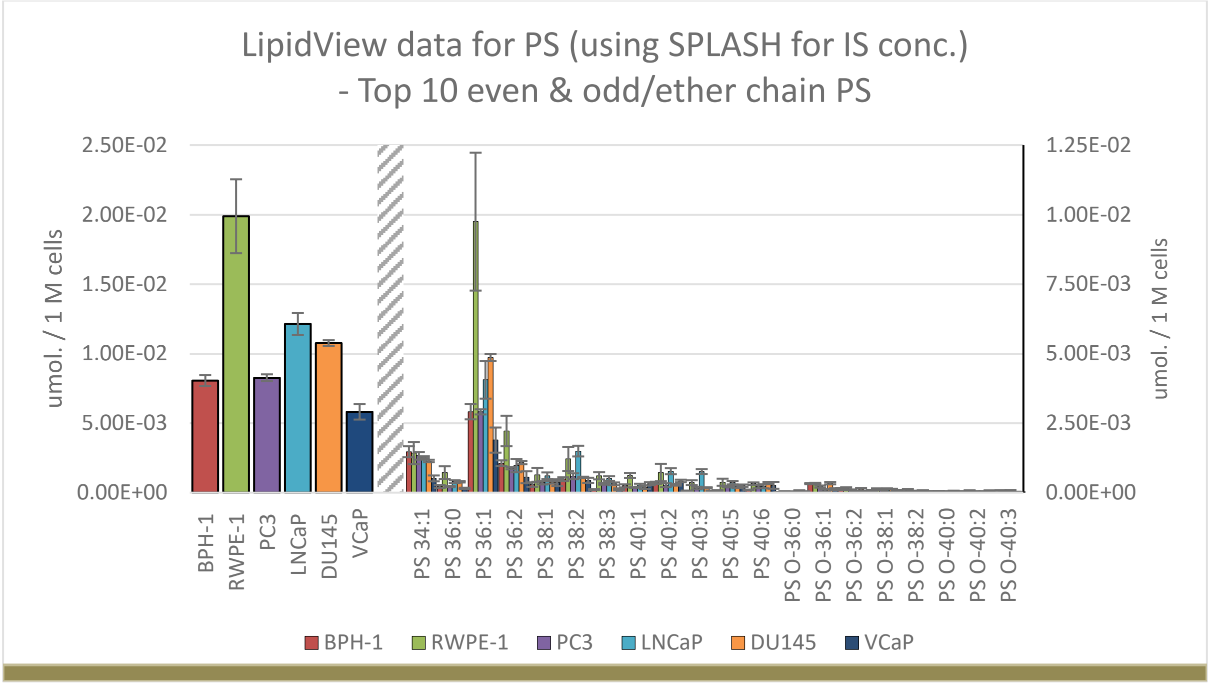

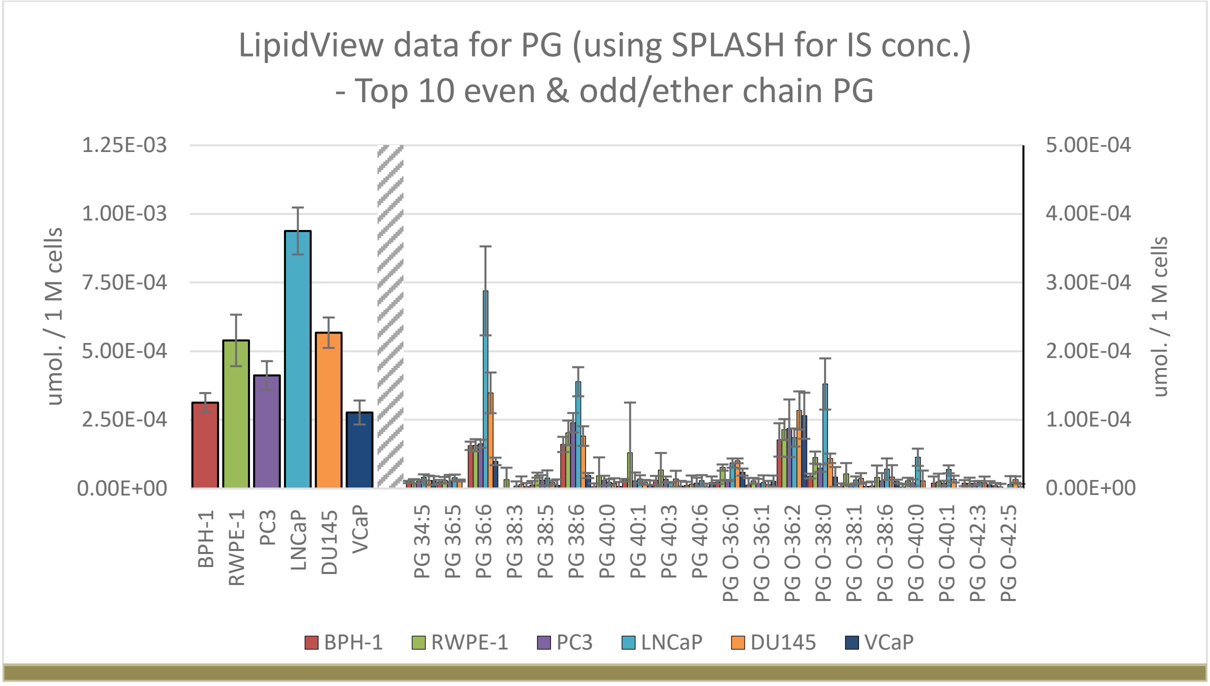

**Supplementary Fig. S2B:** Quantitative complex lipidomics profile from PCa cell lines for: (top) phosphatidylserine (PS) and (bottom) phosphatidylglycerol (PG). Left panels display total subclass lipid abundance and right panels display the top 10 most abundant even chain fatty acyl sum compositions and top 10 most abundant odd/ether chain fatty acyl sum compositions. (n=3; 95% confidence interval error)

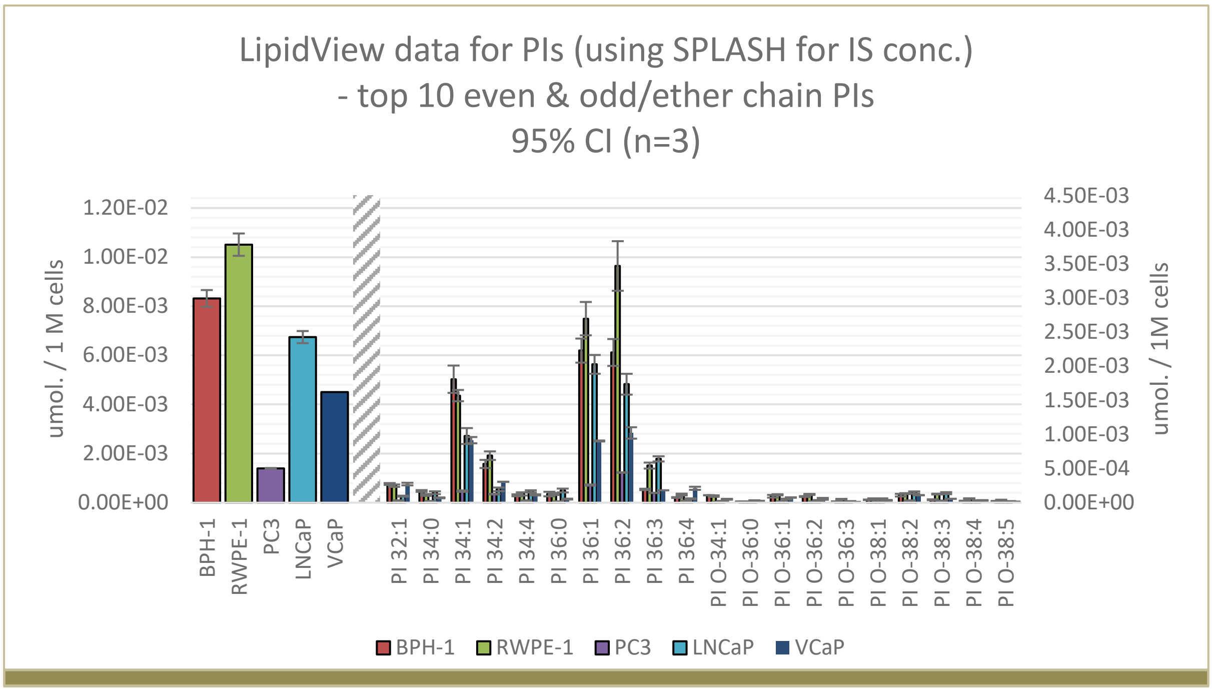

**Supplementary Fig. S2C:** Quantitative complex lipidomics profile from PCa cell lines for: (top) phosphatidylinositol (PI). Left panel displays total subclass lipid abundance and right panel displays the top 10 most abundant even chain fatty acyl sum compositions and top 10 most abundant odd/ether chain fatty acyl sum compositions. (n=3; 95% confidence interval error). DU145 removed from analysis due to technical error.

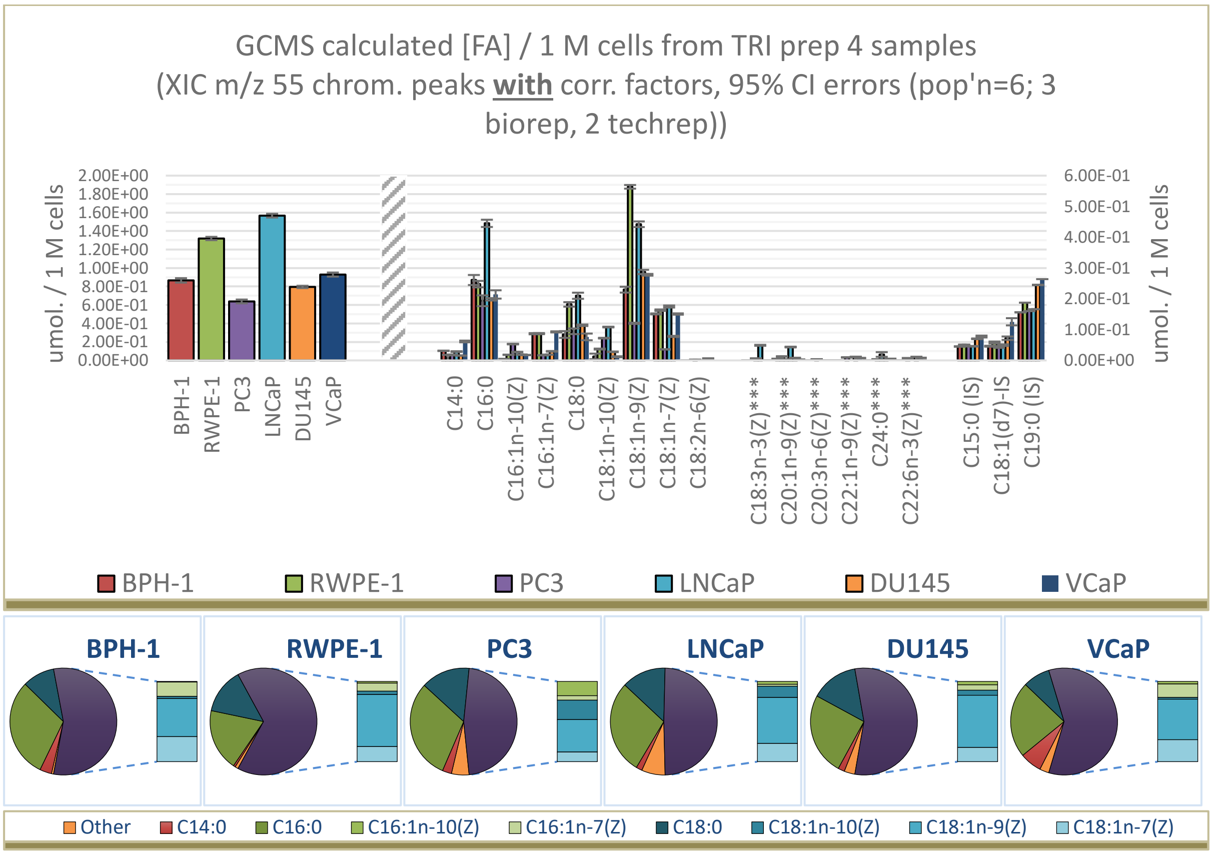

**Supplementary Fig. S3:** Quantitative fatty acyl double bond analysis (GC-MS FAMEs) for PCa cell lines. Top left panel displays total fatty acid abundance per 1 M cells, top right displays quantified individual fatty acyl species (reference material matched), bottom pie charts display relative totals of the individual fatty acyl species. *** indicates hesitancy in assignment due to multiple samples peaks existing around where the reference standard should elute.

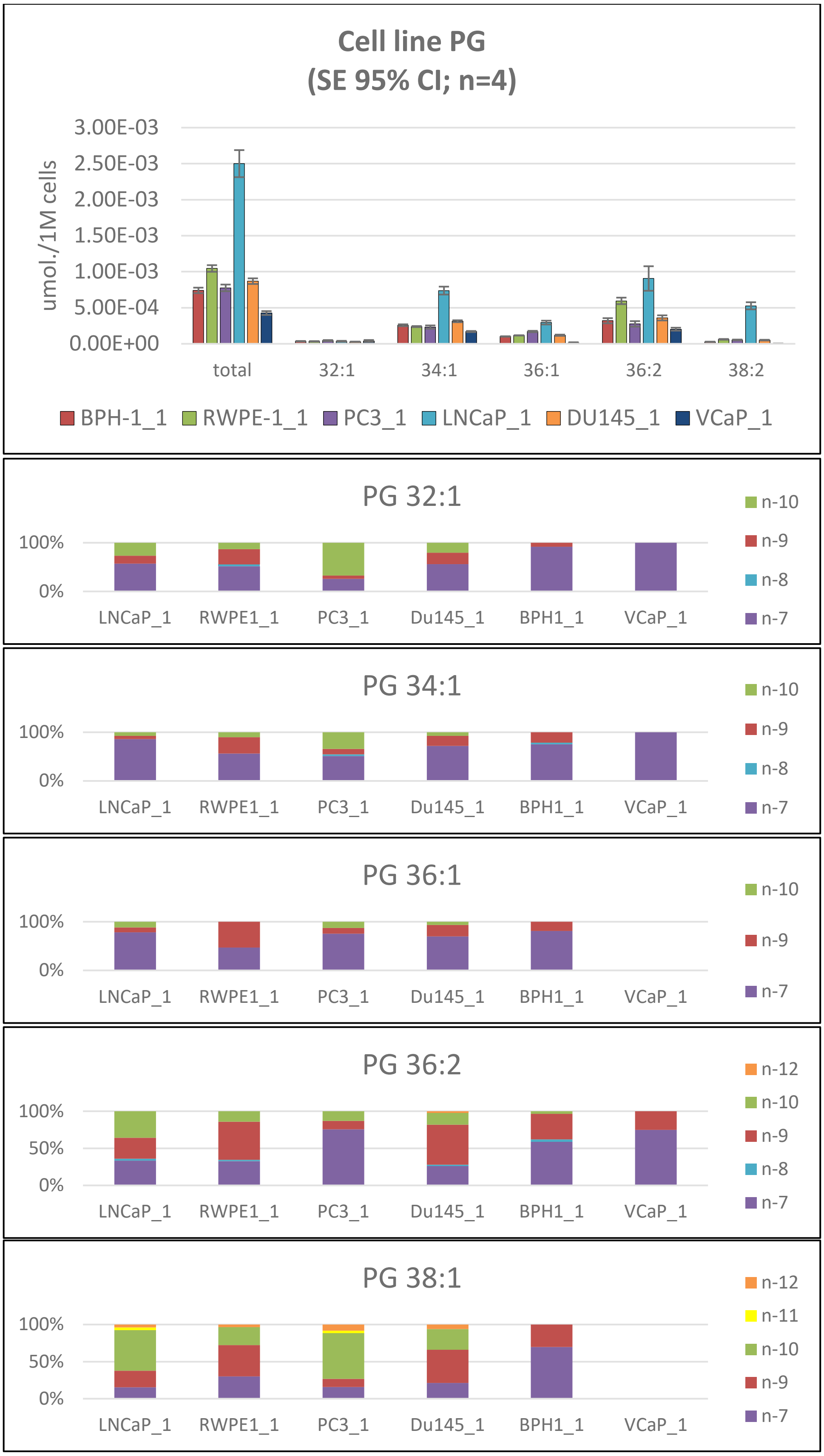

**Supplementary Fig. S4A:** Quantitative lipid abundance from complex lipidomics for PG mono and di unsaturated species and fractional distribution of double bonds. mean and 95% confidence interval displayed (n=3, mean displayed).

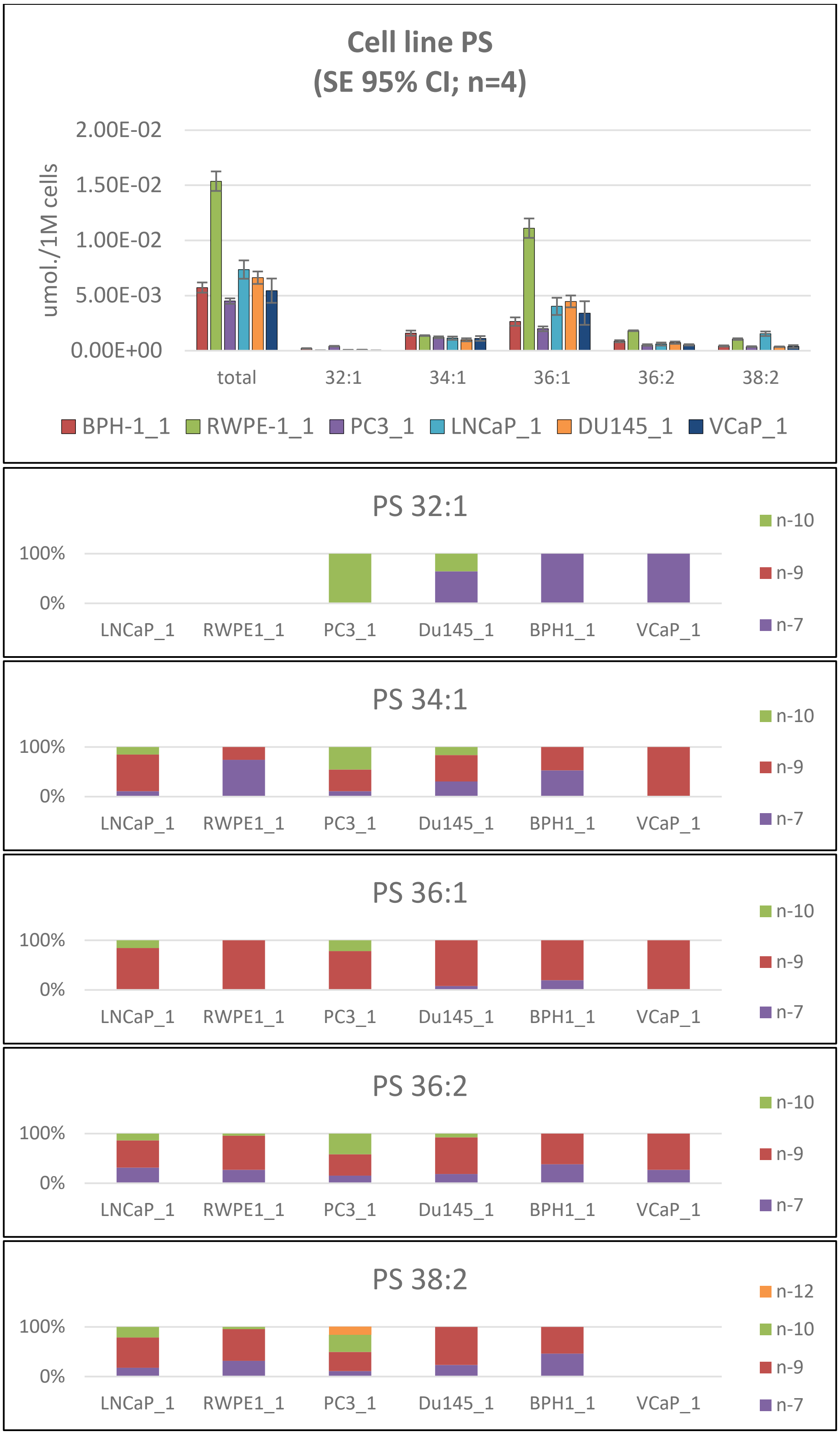

**Supplementary Fig. S4B:** Quantitative lipid abundance from complex lipidomics for PS mono and di unsaturated species and fractional distribution of double bonds. mean and 95% confidence interval displayed (n=3, mean displayed).

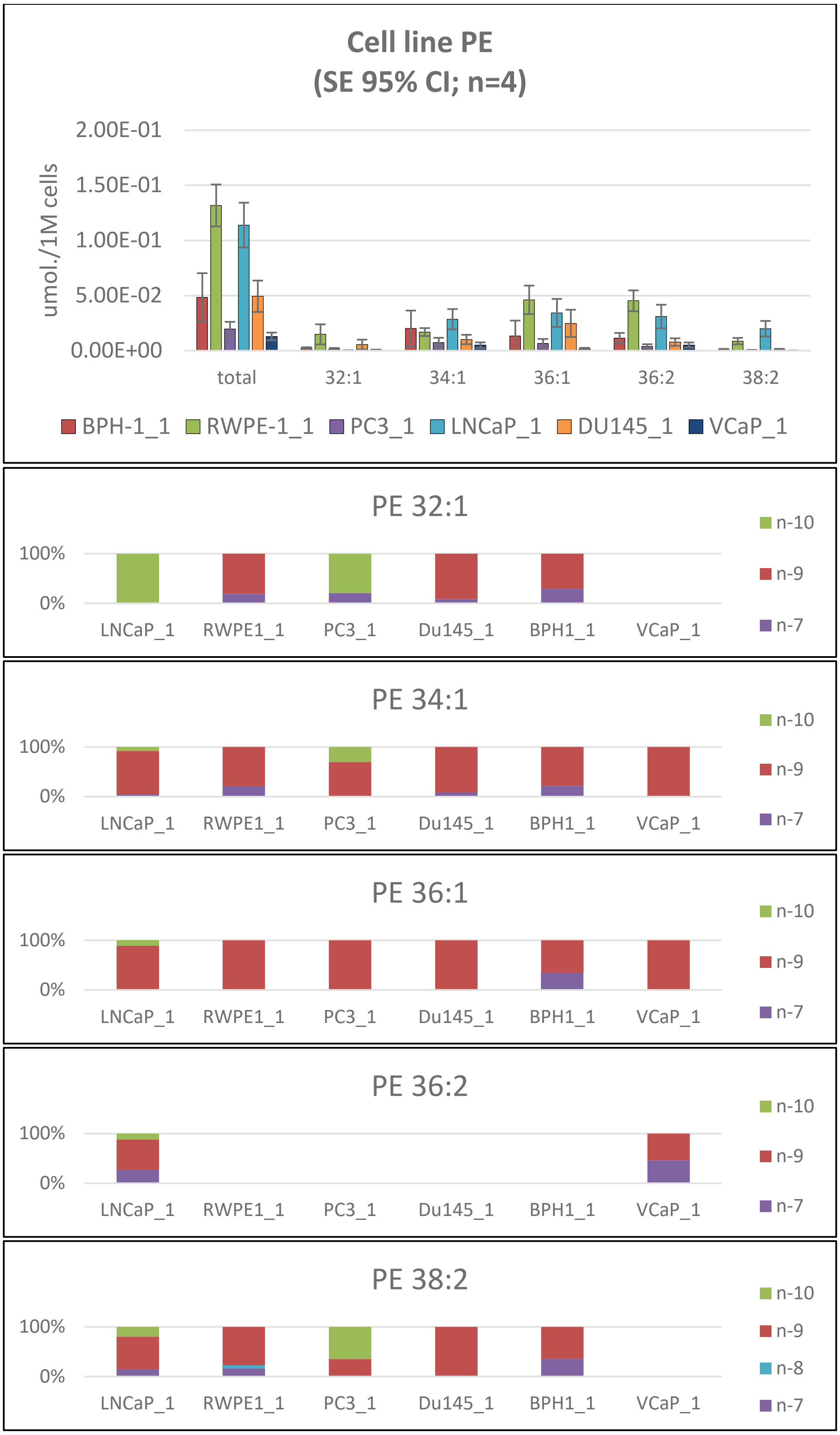

**Supplementary Fig. S4C:** Quantitative lipid abundance from complex lipidomics for PE mono and di unsaturated species and fractional distribution of double bonds. mean and 95% confidence interval displayed (n=3, mean displayed).

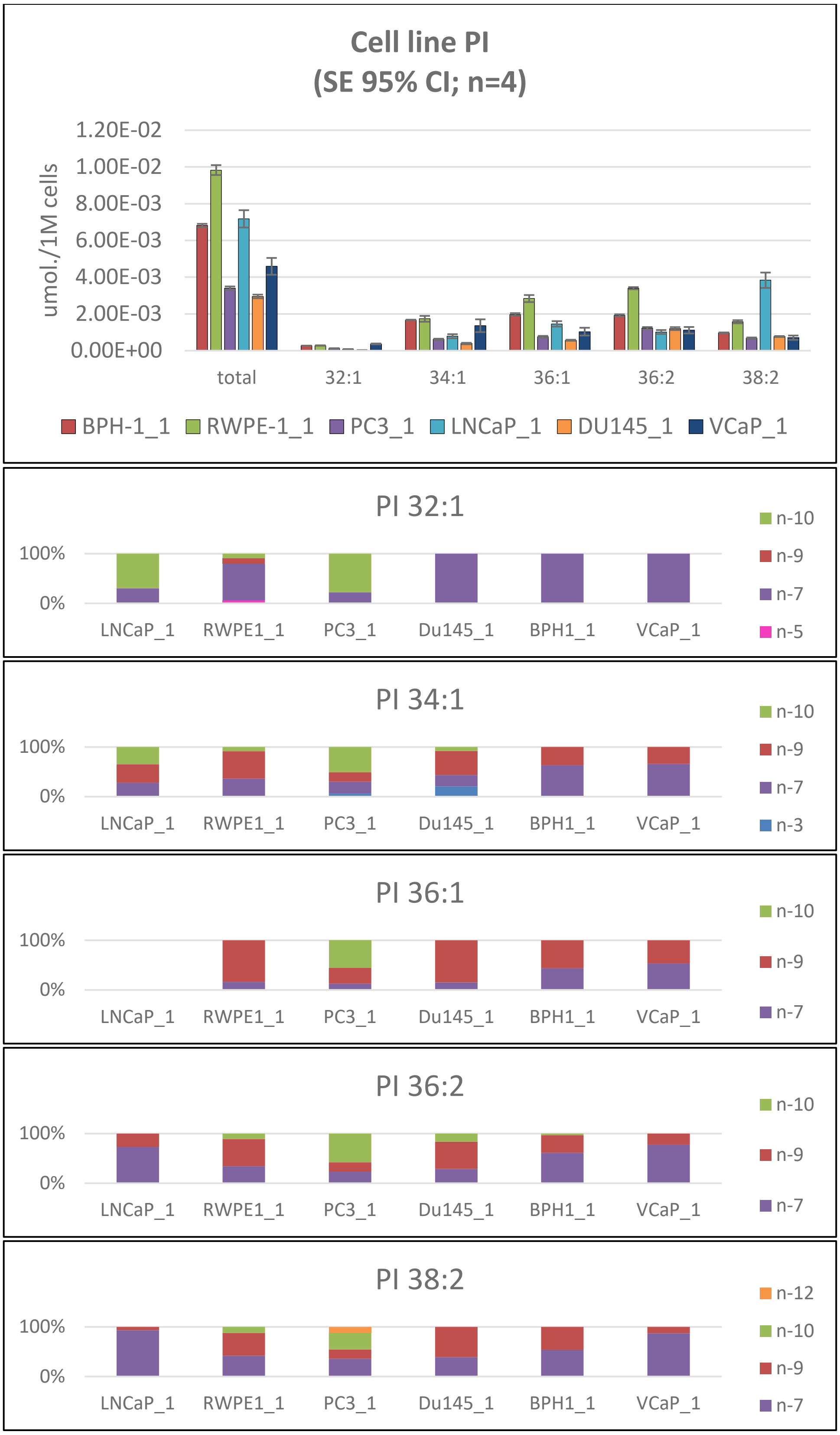

**Supplementary Fig. S4D:** Quantitative lipid abundance from complex lipidomics for PI mono and di unsaturated species and fractional distribution of double bonds. mean and 95% confidence interval displayed (n=3, mean displayed).

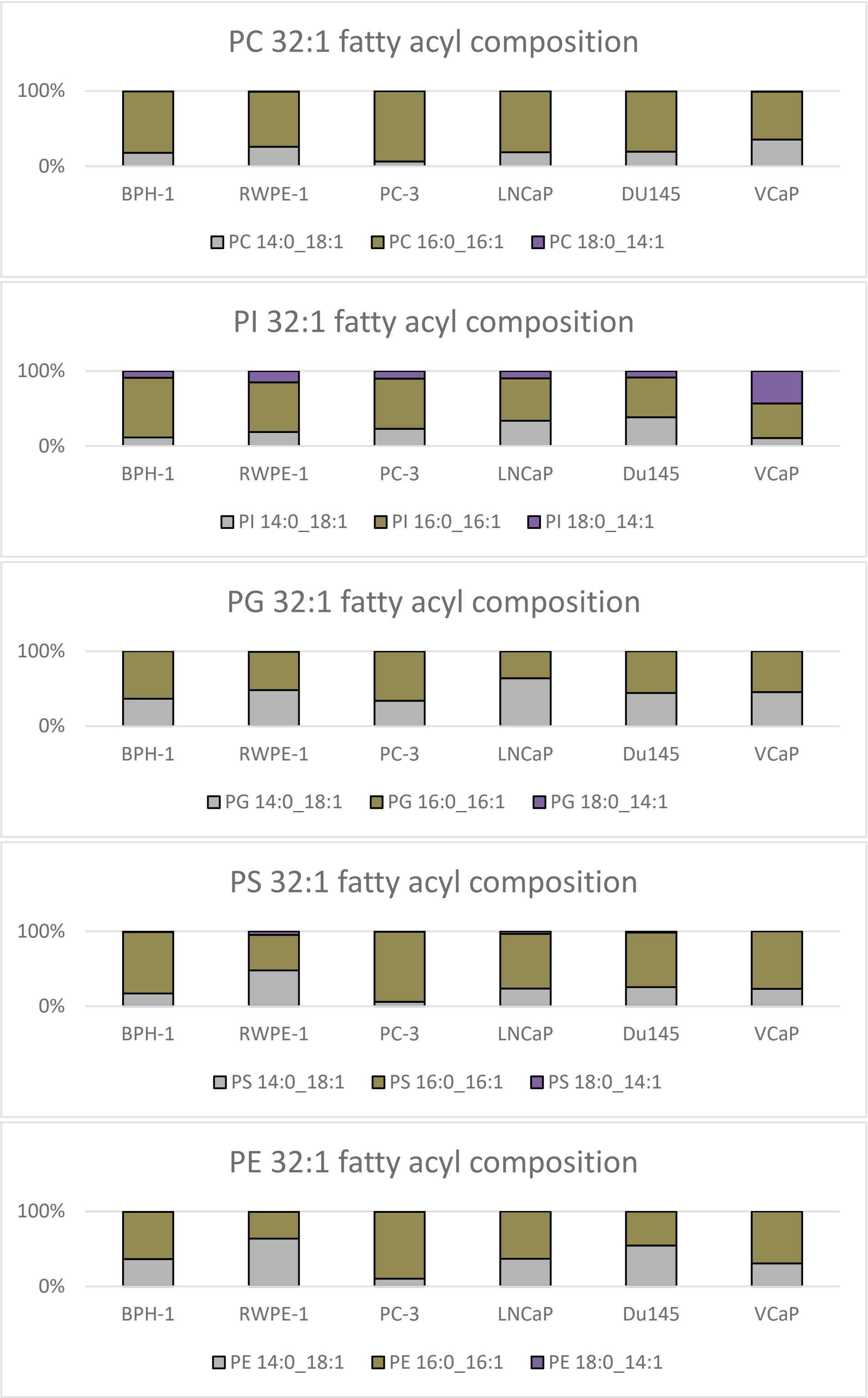

**Supplementary Fig. S5A:** Intact-lipid fatty-acyl compositional analysis for 32:1 glycerophospholipids derived from PCa cell lines. CID/OzID of the $[M+Na]^+$ precursor ion was used for PC analysis and negative $[M-H]^-$ anion MS analysis was used for the remaining lipid sub classes.

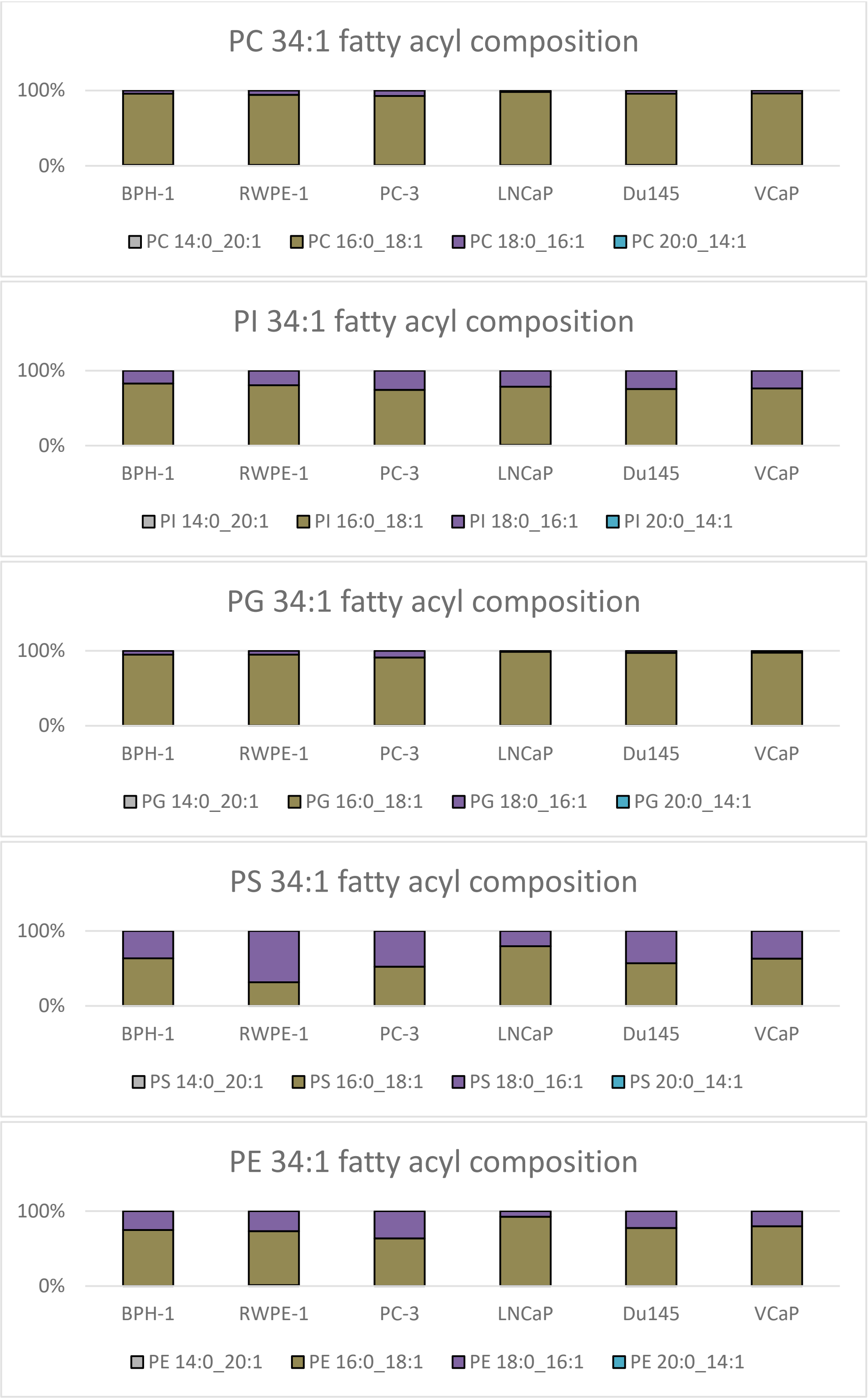

**Supplementary Fig. S5B:** Intact-lipid fatty-acyl compositional analysis for 34:1 glycerophospholipids derived from PCa cell lines. CID/OzID of the $[M+Na]^+$ precursor ion was used for PC analysis and negative $[M-H]^-$ anion MS analysis was used for the remaining lipid sub classes.

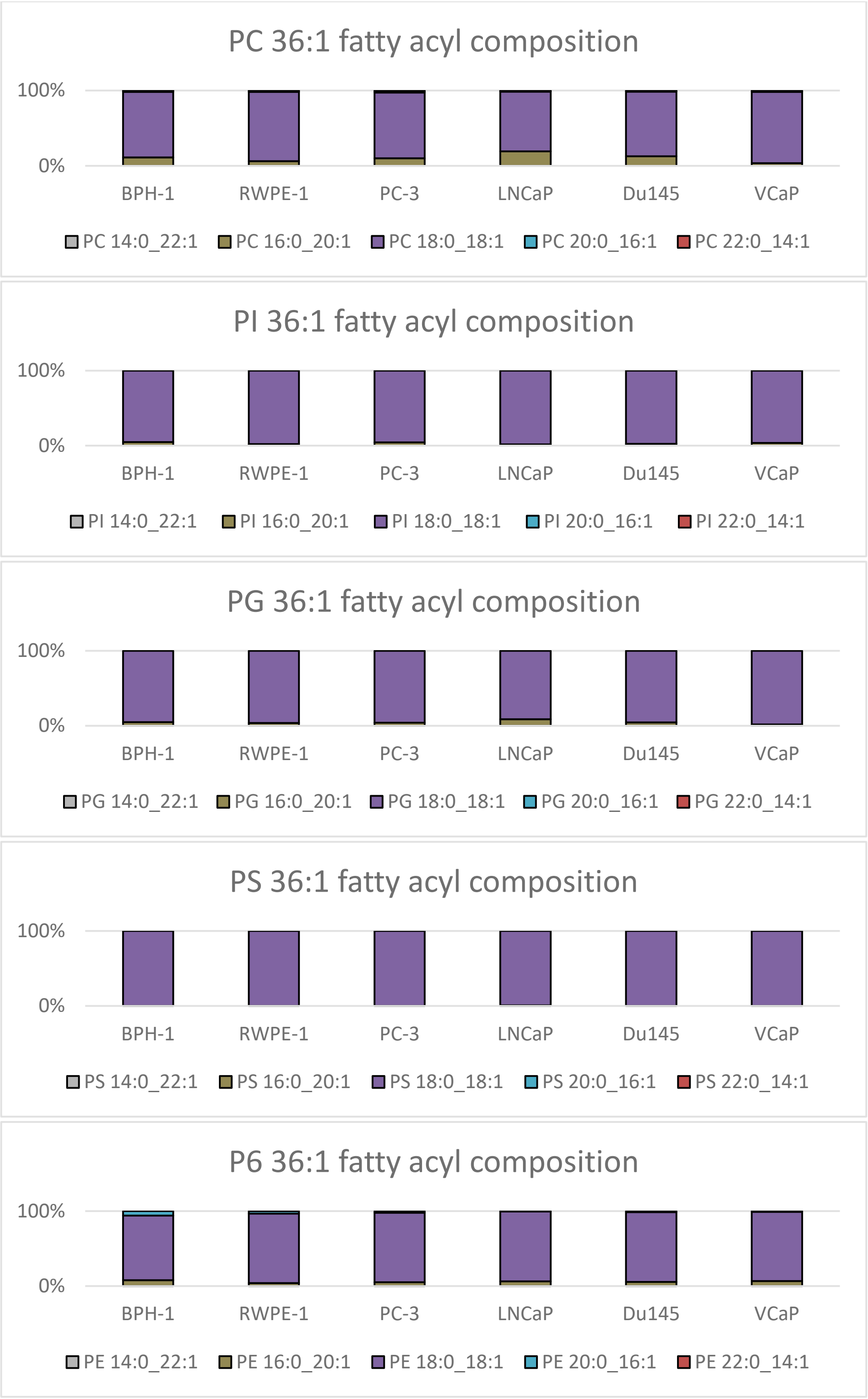

**Supplementary Fig. S5C:** Intact-lipid fatty-acyl compositional analysis for 36:1 glycerophospholipids derived from PCa cell lines. CID/OzID of the $[M+Na]^+$ precursor ion was used for PC analysis and negative $[M-H]^-$ anion MS analysis was used for the remaining lipid sub classes.

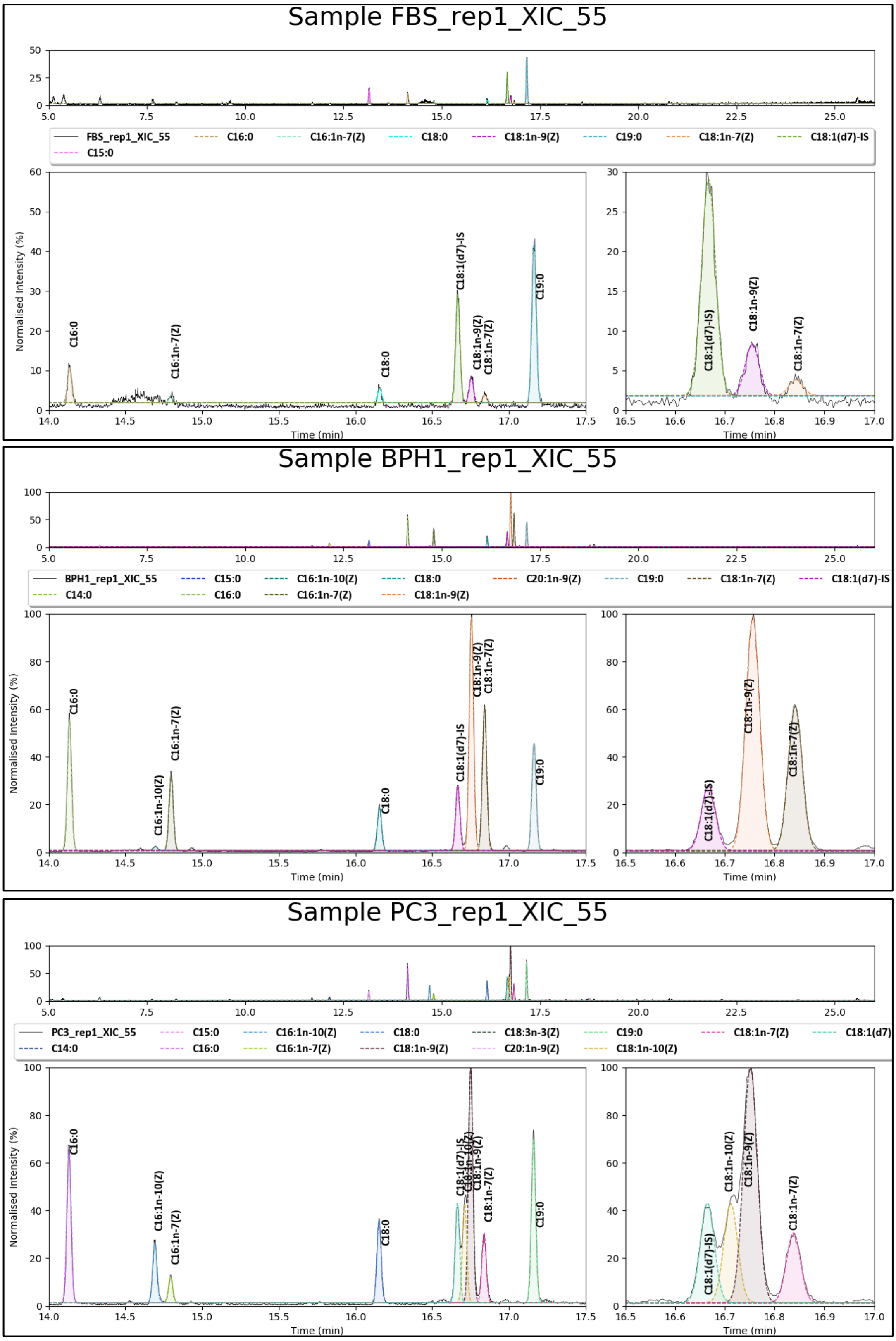

**Supplementary Fig. S6A:** GCMS FAMEs double bond profiles from (top) FBS media, (mid) BPH-1 and (bottom) PC-3. Each individual panel shows the integrated and assigned chromatographic peaks for the full elution profile (top insert), the magnified elution region for the 16-length and 18-length fatty acyl chains (bottom left insert, and a further magnification for the elution region of the 18-length fatty acyl chains (bottom right). As can be seen the chromatographic shoulder-feature for the 18:1*n*-10 is absent in the FBS media (top panel; bottom right insert) showing no presence of FA 18:1*n*-10. Similarly, the 18:1*n*-10 feature of BPH-1 (mid) appears to be fit by the python script, however it is below threshold limitations and more likely to be peak overlap between 18:1*n*-9(d7) and 18:1*n*-9. In contrast, PC-3 (bottom) displays 18:1*n*-10 at ~50% of 18:1*n*-9.

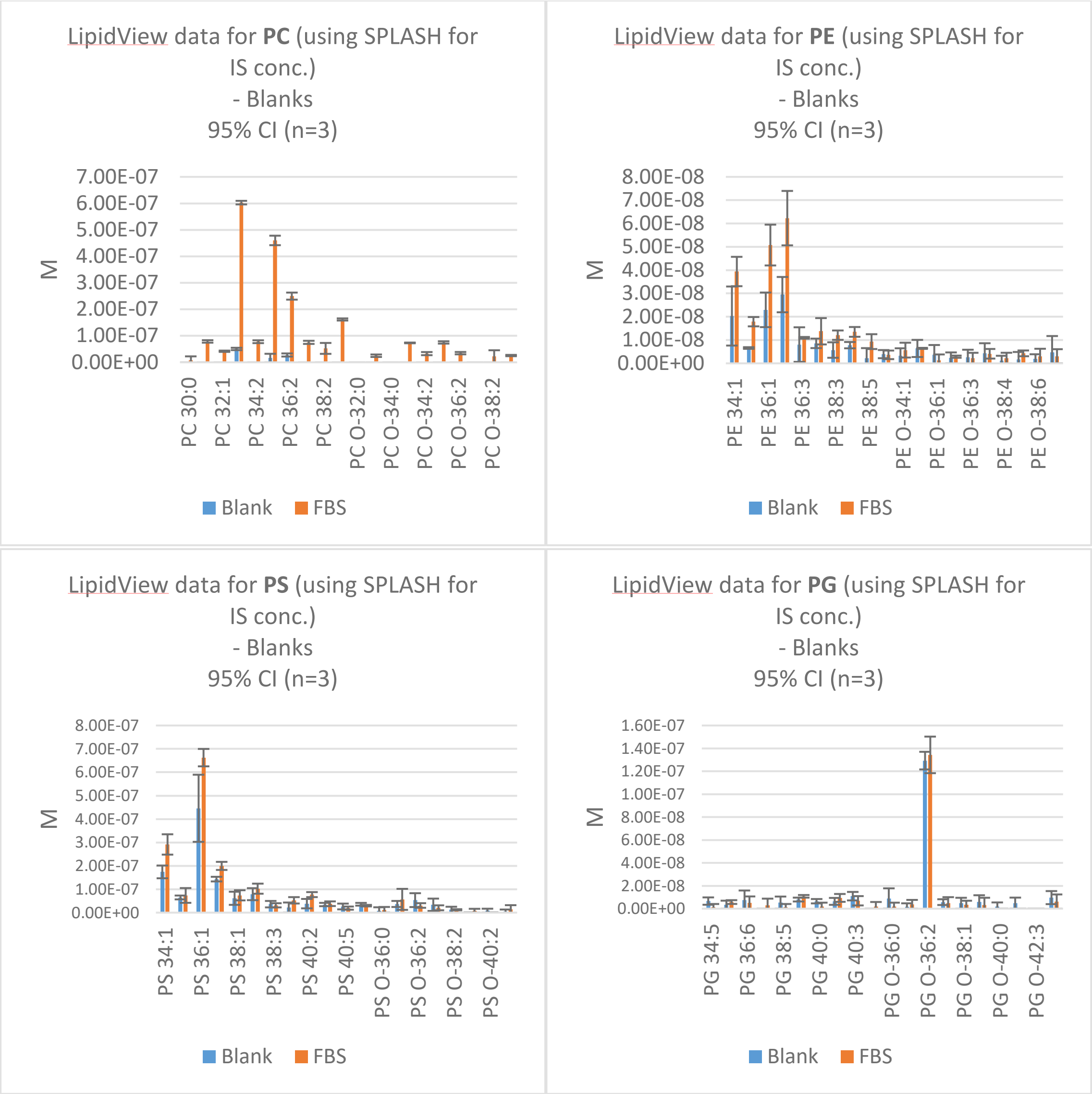

**Supplementary Fig. S6B:** Quantitative complex lipidomics profile from FBS media (orange) and lipid extraction method blank (blue). Complex lipidomics for: PC (top left), PE (top right), PS (bottom left) and PG (bottom right). Panels display the top 5 most abundant even chain fatty acyl sum compositions and top 5 most abundant odd/ether chain fatty acyl sum compositions. (n=3; mean and 95% confidence interval error).

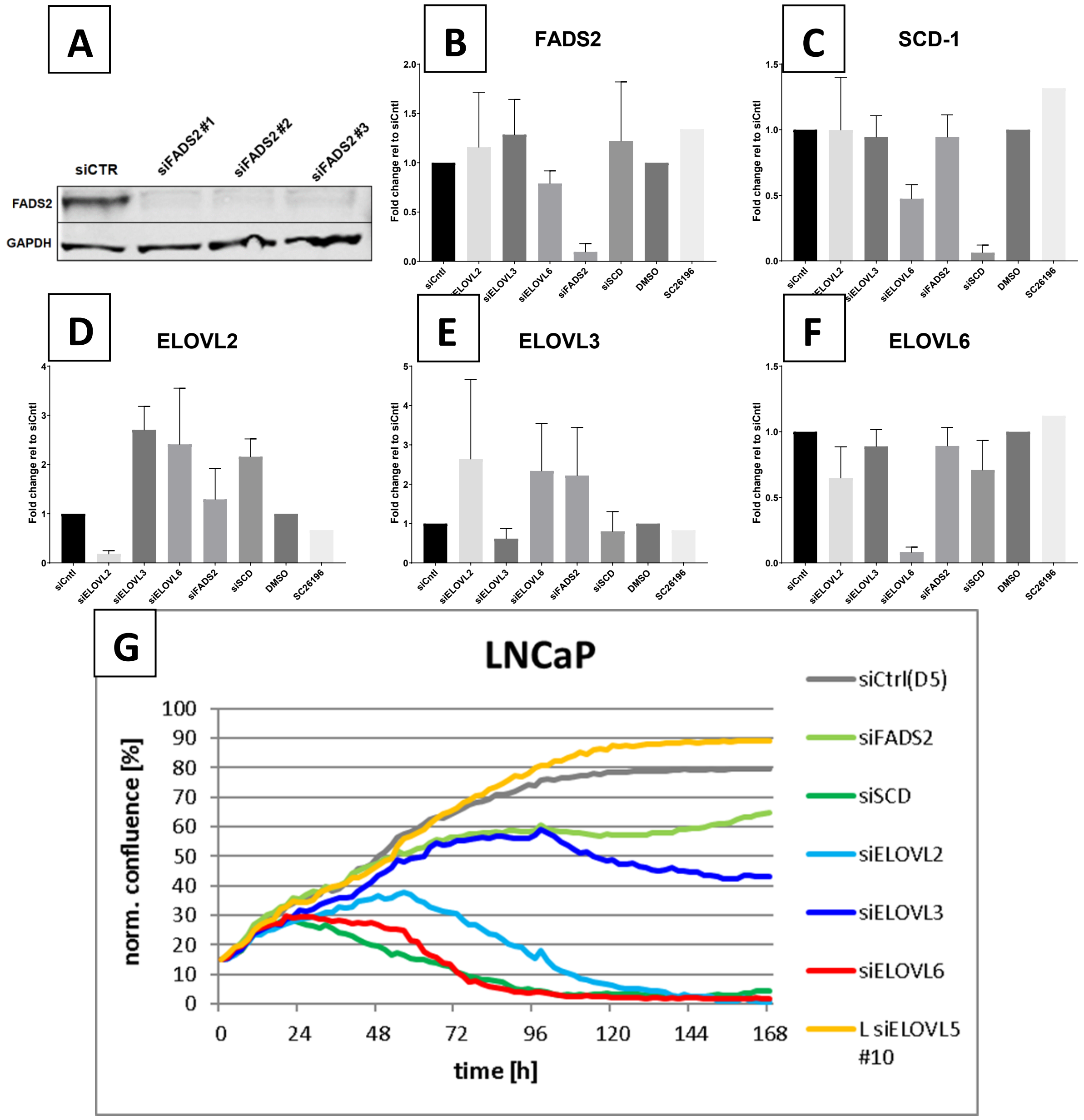

**Supplementary Fig. S7**: Cellular studies confirming the depletion of the FADS2 protein (Western blot), post siRNA treatment LNCaP mRNA expression (qRT-PCR) and post siRNA treatment cell confluence by Incucyte. A) Western blot analysis confirming the depletion of FADS2 protein after FADS2 siRNA-mediated gene silencing with three different siRNA sequences (lane 2= , lane 3= , lane 4= ) for 72 hours. The position of FADS2 protein (~52 kDa) is indicated. Even protein loading was controlled by Ponceau S staining and immunoblotting of GAPDH. B-F) qRT-PCR derived mRNA profiles of LNCaP cells treated with siFADS2, siSCD-1, siELOVL2, siELOVL3 and siELOVL6 (for primer sequences cf. supplementary Fig. 12). G) Normalised LNCaP cell confluence after siRNA treatment from 0-168 hours.

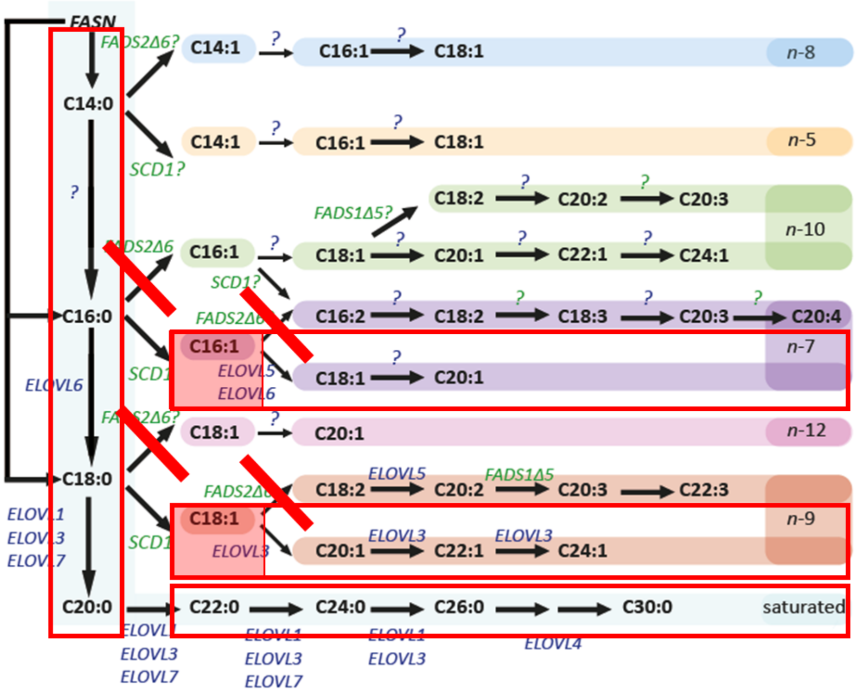

**Supplementary Fig. S8A:** Theoretical fatty acyl profile after siFADS2 treatment. Solid-lines indicate knockdown effect, hollow boxes indicate increased species abundance and shaded boxes represent further increases in species abundance due to subsequent modifications being rate limiting.

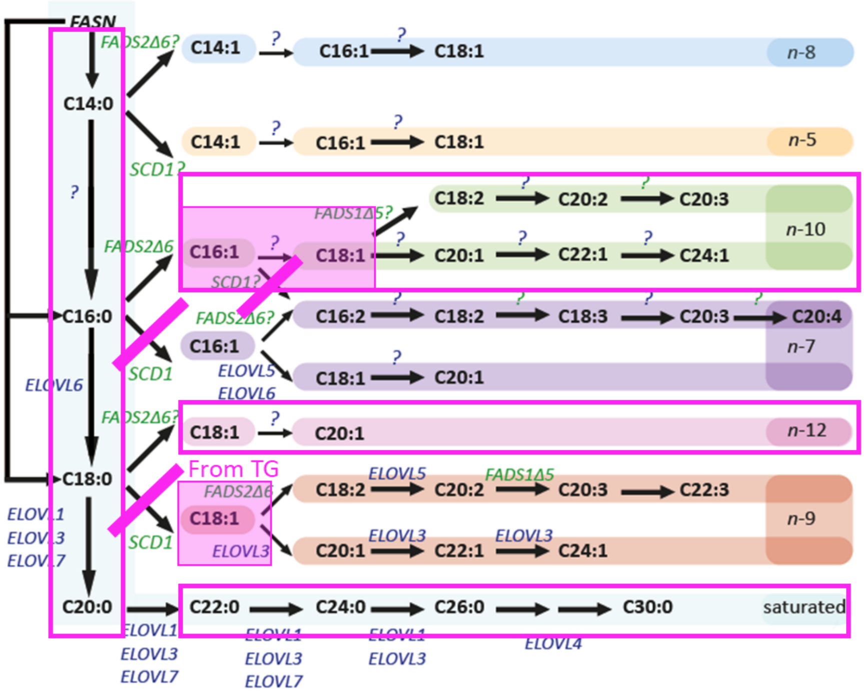

**Supplementary Fig. S8B:** Theoretical fatty acyl profile after siSCD-1 treatment. Solid-lines indicate knockdown effect, hollow boxes indicate increased species abundance and shaded boxes represent further increases in species abundance due to subsequent modifications being rate limiting. A significant population of 18:1*n*-9 is known to exist in the triacylglycerols and it is speculated that these will supply the fatty acyl pool under treatment conditions. Based on visual comparison siSCD-1 should present negative correlations with siFADS2 (S. Fig. 6 A) and can be seen to do so in Fig. 4 B.

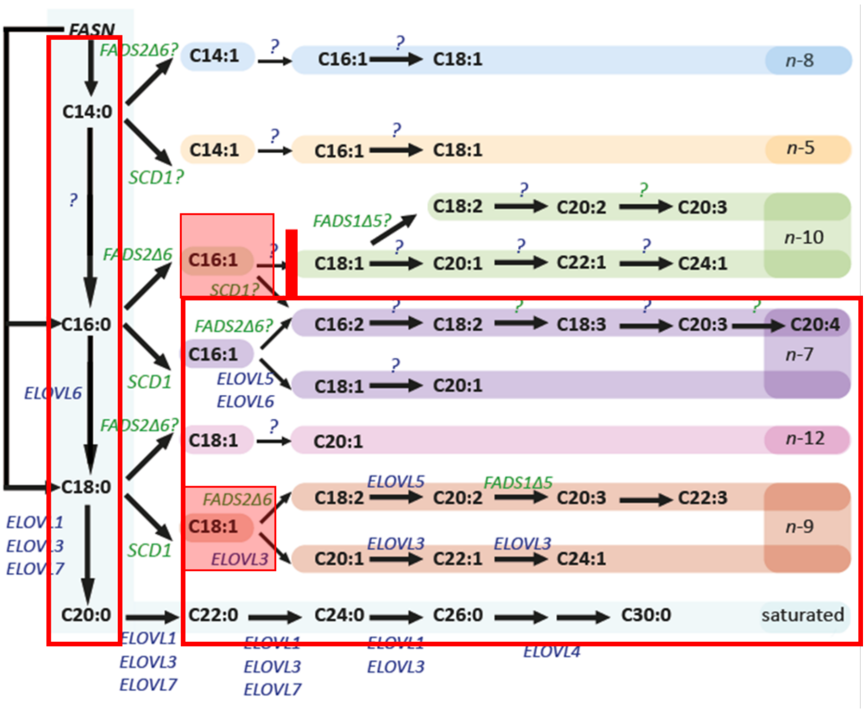

**Supplementary Fig. S8C:** Theoretical fatty acyl profile after siELOVLx treatment, (proposed to be ELOVL2). Solid-lines indicate knockdown effect, hollow boxes indicate increased species abundance and shaded boxes represent further increases in species abundance due to subsequent modifications being rate limiting. Visual comparison of siELOVLx against siSCD-1 (S. Fig. 6B) presents that unsaturation profiles should be positively correlated due to similarity in major contributors. Data observations from Fig. 4 B and C suggest that siSCD-1 only has strong positive correlations with siELOVL2 and siELOVL6.

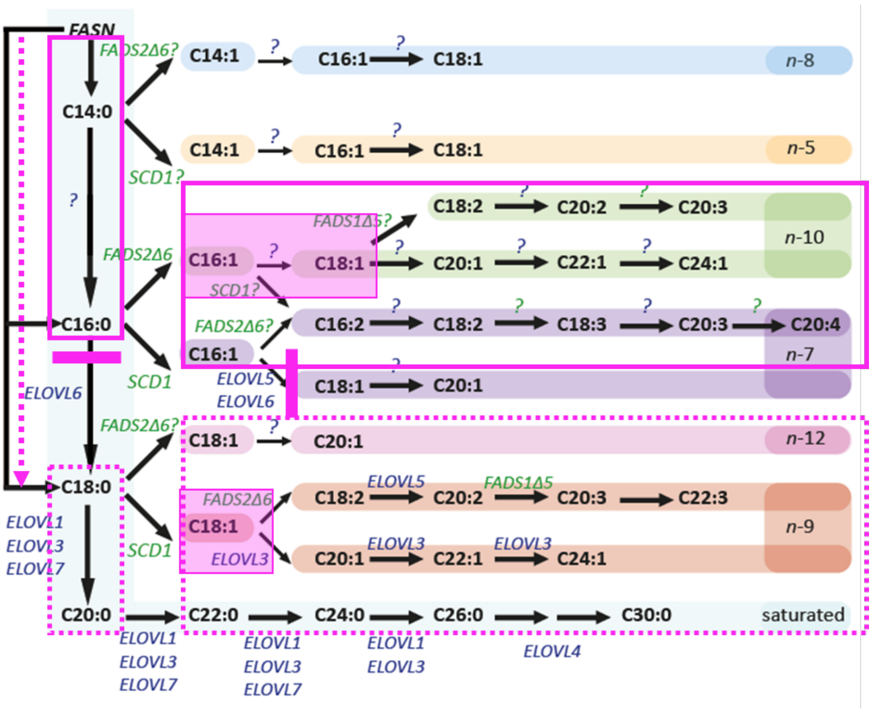

**Supplementary Fig. S8D:** Theoretical fatty acyl profile after siELOVL6 treatment. Solid-lines indicate knockdown effect, hollow boxes indicate increased species abundance and shaded boxes represent further increases in species abundance due to subsequent modifications being rate limiting. Dotted boxes present potential alternative routes for synthesis. Visual comparison of siELOVL6 against siSCD-1 (S. Fig. 6B) presents that fatty acyl unsaturation profiles should be positively correlated due to similarity in major contributors. Similarly, strong positive correlations can be seen with S. Fig. 6 C (siELOVL-x). However, the data observations of Fig. 4 C shows that ELOVL2 is the only tested elongase to be strongly correlated to ELOVL6. This again suggests that ELOVLx is in fact ELOVL 2 and responsible for the elongation between 16:1*n*-10→18:1*n*-10.

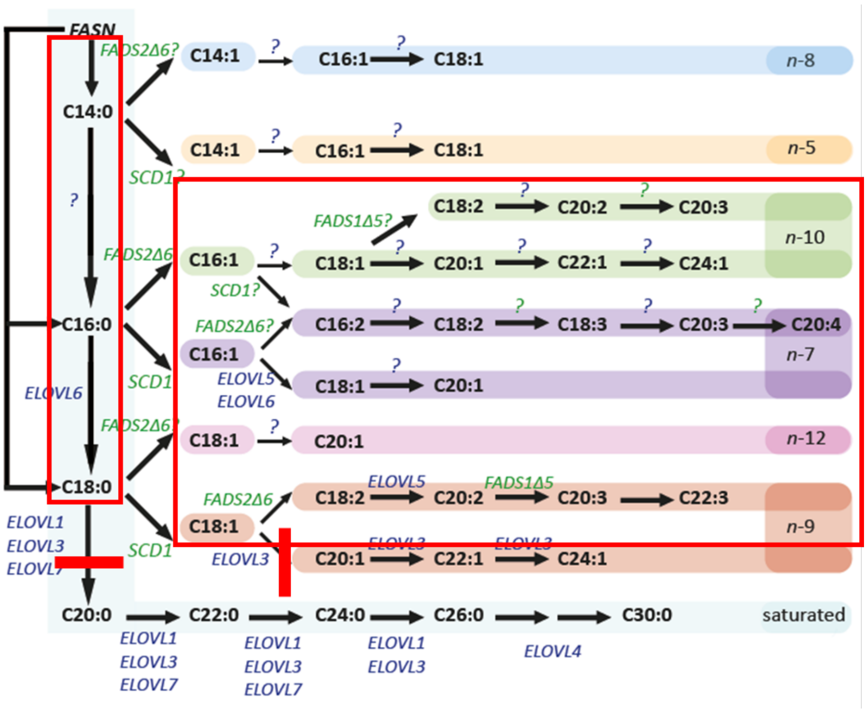

**Supplementary Fig. S8E:** Theoretical fatty acyl profile after siELOVL3 treatment. Solid-lines indicate knockdown effect, hollow boxes indicate increased species abundance and shaded boxes represent further increases in species abundance due to subsequent modifications being rate limiting. Aside from minor correlations with other ELOVL knockdowns because of an overall decrease in FA chain length, siELOVL3 profiles shouldn't display significant correlation to other treatments.

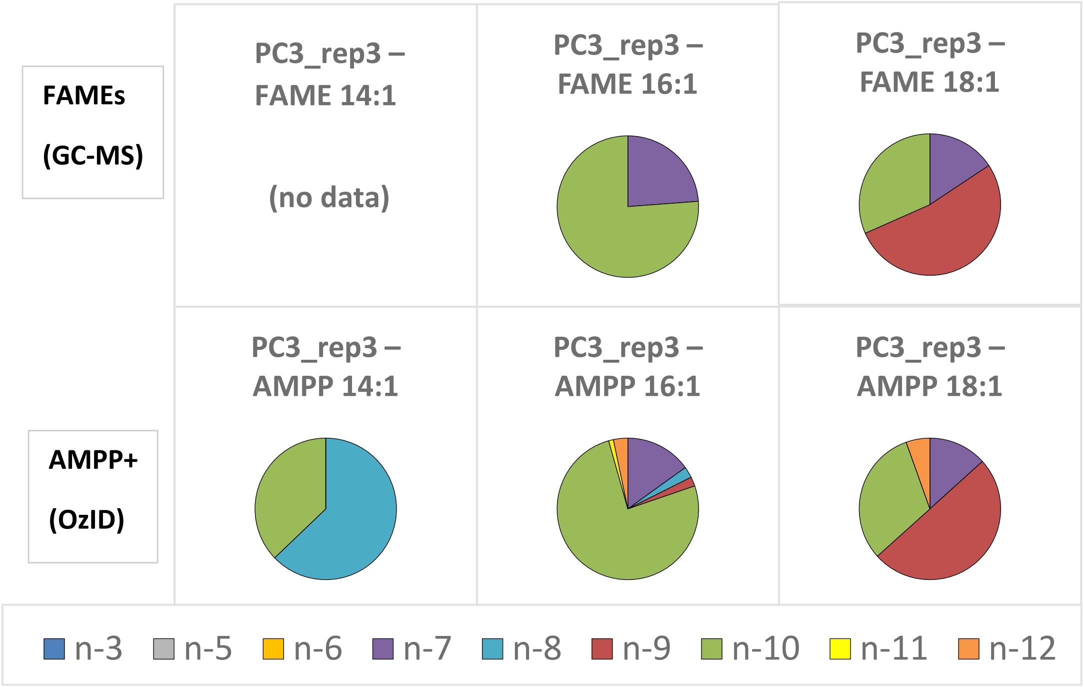

**Supplementary Fig. S9**: Comparison between fatty acyl double bond analysis derived from GC-MS FAMEs and AMPP+ OzID. Because the AMPP+ derivatisation products have a fixed cationic charge, anolyte detection is not reliant on ESI. This means that less abundant FA species, such as the FA 14:1 and the minute isomers of FA 16:1 and FA 18:1, can be detected. Secondly, OzID double bond analysis of AMPP+ derivatives is achieved via mass spectrometric fragmentation principals and therefore requires no reference standard for identification. We recognise that there may be some minor fragmentation biases arising from OzID – the fractional distribution of each isomer is hence reflective of the molar % contribution to each fatty acyl composition.

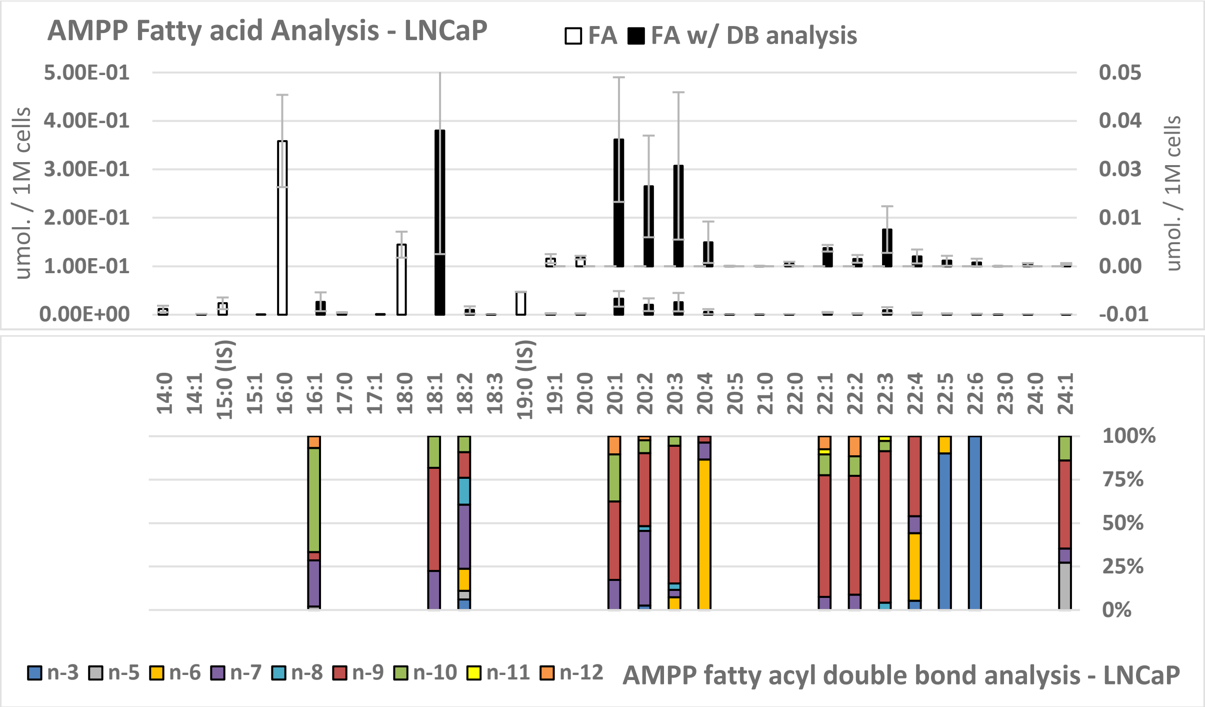

**Supplementary Fig. S10A:** Hydrolysed fatty acyl quantitative abundance and double bond analysis of metastatic-LNCaP using AMPP derivatisation and OzID. (n=3; mean and 95 % confidence interval error)

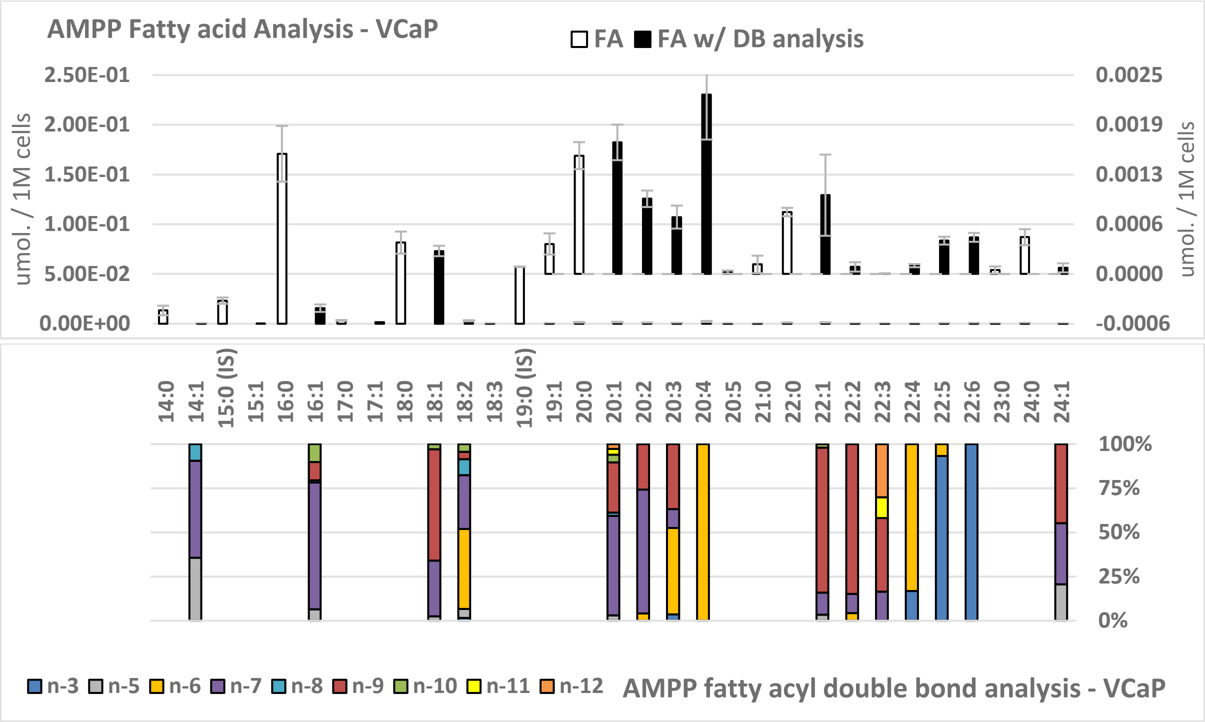

**Supplementary Fig. S10B** Hydrolysed fatty acyl quantitative abundance and double bond analysis of metastatic-VCaP using AMPP derivatisation and OzID. (n=3; mean and 95 % confidence interval error)

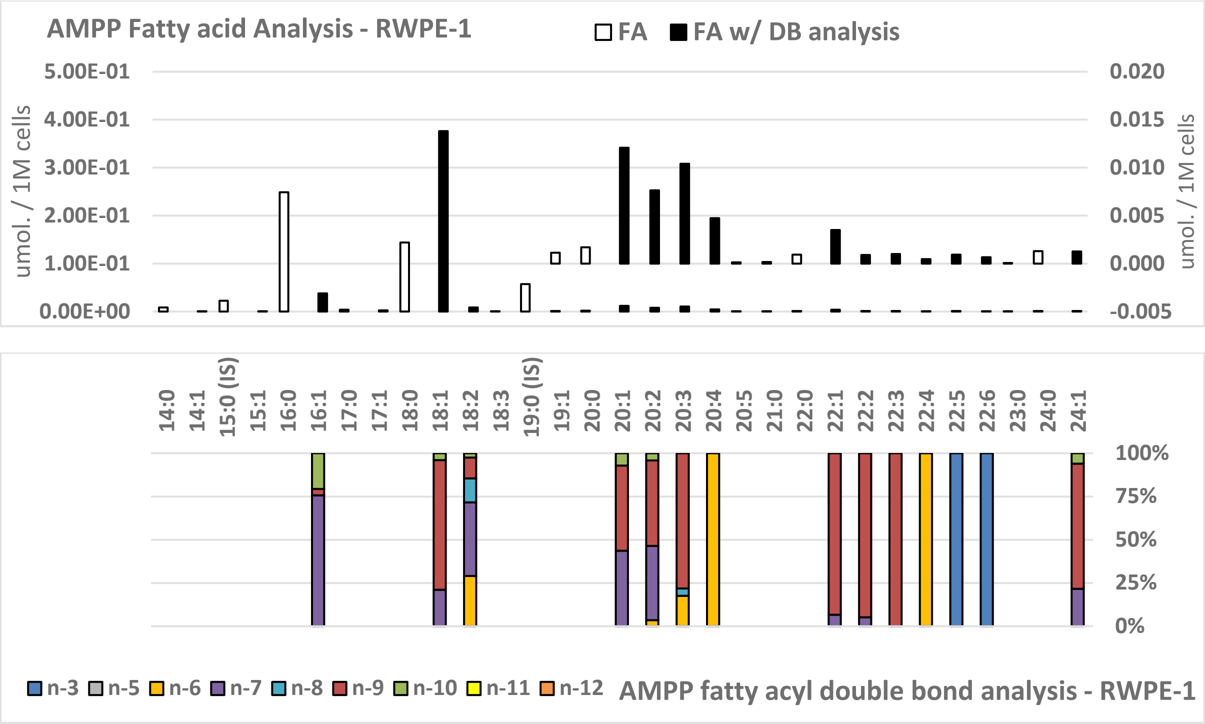

**Supplementary Fig. S10C:** Hydrolysed fatty acyl quantitative abundance and double bond analysis of non-metastatic-RWPE-1 using AMPP derivatisation and OzID. (n=1; no error analysis).

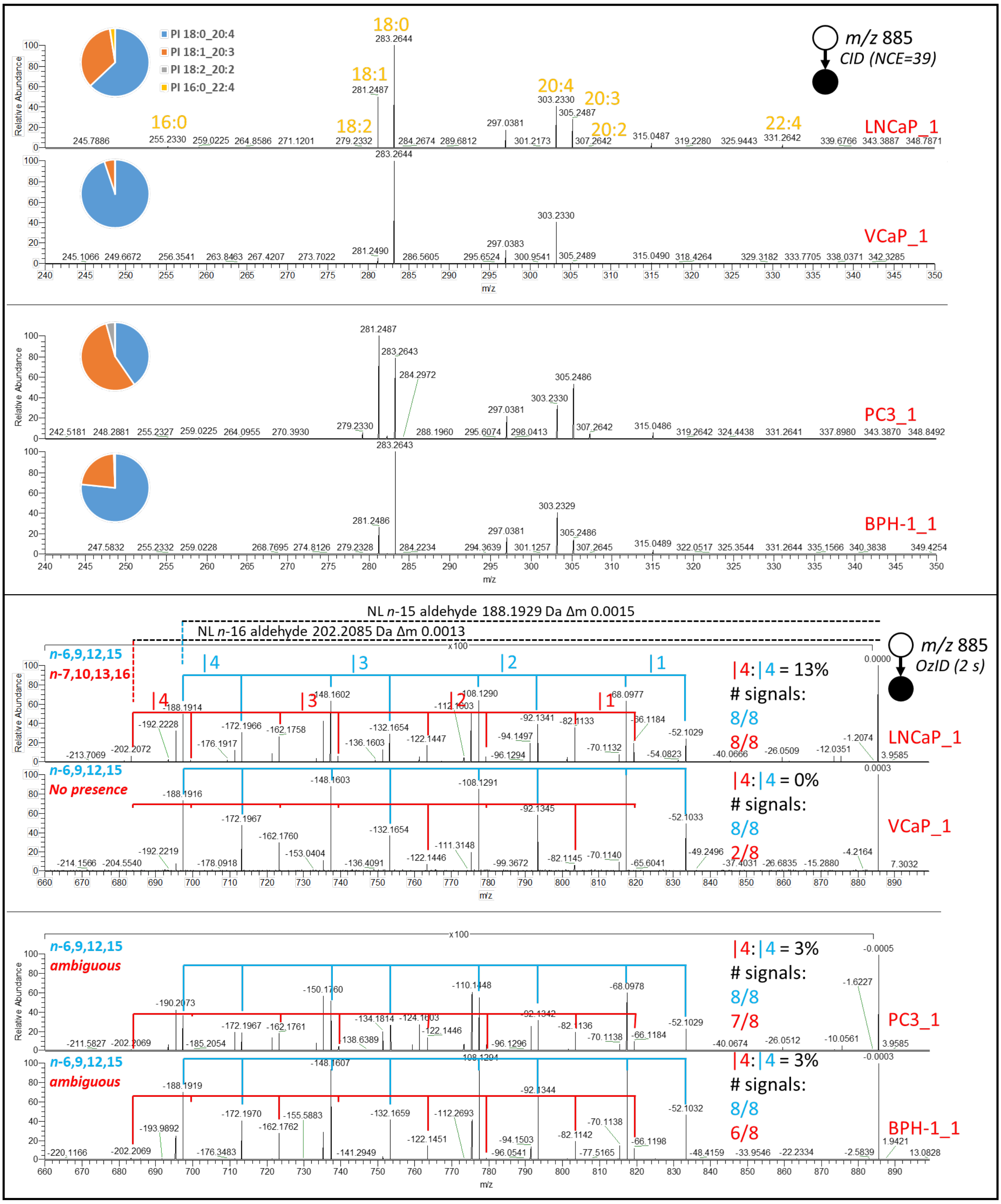

**Supplementary Fig. S11A:** Conventional fatty acyl anion analysis of PI 38:4 and OzID lipid double bond analysis of PI 38:4 within 4 cell lines. Top panel shows that PI 18:0_20:4 is the most abundant species for LNCaP, VCaP and BPH-1, while PC-3 displays a slightly higher abundance of PI 18:1_20:3 (as represented in pie charts). Other species that are present include PI 16:0_22:4 and PI 18:2_20:2. Bottom panel displays the fatty acyl double bond analysis for the cell lines and reveals that LNCaP unambiguously contains the unusual 20:4*n*-7 polyunsaturated fatty acid, while PC-3 and BPH-1 display ambiguous assignment with PC-3 being slightly more confirmatory based on signal analysis. VCaP on the other hand definitively displays absence of the 20:4*n*-7 species in this analysis. Signal analyses were based on comparisons of the 4th sequential double bonds (|4) and presence of the 8 total characteristic product ions. Other unusual double bond sequences that can be discerned from this analysis include the *n*-12,15; *n*-10,13; *n*-8,11,14; *n*-9,12,15 and *n*-10,13,16 but cannot be assigned to specific lipid fatty acyl combinations.

| n- (\|1) | Aldehyde | Criegee | n- (\|2) | Aldehyde | Criegee | n- (\|3) | Aldehyde | Criegee | n- (\|4) | Aldehyde | Criegee |
|---|---|---|---|---|---|---|---|---|---|---|---|
| 3 | 26.0520 | 10.0571 | 3 | 24.0364 | 8.0415 | 3 | 22.0207 | 6.0258 | 3 | 20.0051 | 4.0102 |
| 4 | 40.0677 | 24.0728 | 4 | 38.0520 | 22.0571 | 4 | 36.0364 | 20.0415 | 4 | 34.0207 | 18.0258 |
| 5 | 54.0833 | 38.0884 | 5 | 52.0677 | 36.0728 | 5 | 50.0520 | 34.0571 | 5 | 48.0364 | 32.0415 |
| 6 | 68.0990 | 52.1041 | 6 | 66.0833 | 50.0884 | 6 | 64.0677 | 48.0728 | 6 | 62.0520 | 46.0571 |
| 7 | 82.1146 | 66.1197 | 7 | 80.0990 | 64.1041 | 7 | 78.0833 | 62.0884 | 7 | 76.0677 | 60.0728 |
| 8 | 96.1303 | 80.1354 | 8 | 94.1146 | 78.1197 | 8 | 92.0990 | 76.1041 | 8 | 90.0833 | 74.0884 |
| 9 | 110.1459 | 94.1510 | 9 | 108.1303 | 92.1354 | 9 | 106.1146 | 90.1197 | 9 | 104.0990 | 88.1041 |
| 10 | 124.1616 | 108.1667 | 10 | 122.1459 | 106.1510 | 10 | 120.1303 | 104.1354 | 10 | 118.1146 | 102.1197 |
| 11 | 138.1772 | 122.1823 | 11 | 136.1616 | 120.1667 | 11 | 134.1459 | 118.1510 | 11 | 132.1303 | 116.1354 |
| 12 | 152.1929 | 136.1980 | 12 | 150.1772 | 134.1823 | 12 | 148.1616 | 132.1667 | 12 | 146.1459 | 130.1510 |
| 13 | 166.2085 | 150.2136 | 13 | 164.1929 | 148.1980 | 13 | 162.1772 | 146.1823 | 13 | 160.1616 | 144.1667 |
| 14 | 180.2242 | 164.2293 | 14 | 178.2085 | 162.2136 | 14 | 176.1929 | 160.1980 | 14 | 174.1772 | 158.1823 |
| 15 | 194.2398 | 178.2449 | 15 | 192.2242 | 176.2293 | 15 | 190.2085 | 174.2136 | 15 | 188.1929 | 172.1980 |
| 16 | 208.2555 | 192.2606 | 16 | 206.2398 | 190.2449 | 16 | 204.2242 | 188.2293 | 16 | 202.2085 | 186.2136 |

**Supplementary Fig. S11B:** OzID lipid double bond neutral loss table used to assign double bond positions. In the case of polyunsaturated species, bar nomenclature (*i.e.,* |1) is used to indicate the placement ordering in a sequence of double bonds. *e.g.,* 20:4*n*-6 (*i.e.,* 20:4*n*-6,9,12,15); where |1 = *n*-6, |2 = *n*-9, |3 = *n*-12 and |4 = *n*-15.

| Enzyme | Direction | Sequence |
|---|---|---|
| FADS2 | FADS2fwd | CCCGGCACAACTTACACA |
| | FADS2rev | CCATGCTTGGCACATAGACACTT |
| SCD1 | SCD1fwdexon4 | CCAGCTGTCAAAGAGAAGG |
| | SCD1frevexon5 | AAATACCAGGGCACAAGC |
| ELOVL2 | ELOVL2fwdexon5 | GTGTGTCTTGAACTGGATACC |
| | ELOVL2revexon6 | TCCACCAAAGATACTTGTGC |
| ELOVL3 | ELOVL3fwdexon2 | CTACATGAAGGAACGCAAGG |
| | ELOVL3revexon3 | ACACGGTTTGCTTTAGGC |
| ELOVL6 | ELOVL6fwdexon2 | GAAGCCATTAGTGCTCTGG |
| | ELOVL6revexon3 | ACAAACTGACTGCTTCAGG |

**Supplementary Fig. S12:** Primer sequences used for qRT-PCR experiments through-out. (cf. Methods section 3.6 for qRT-PCR experimental method)

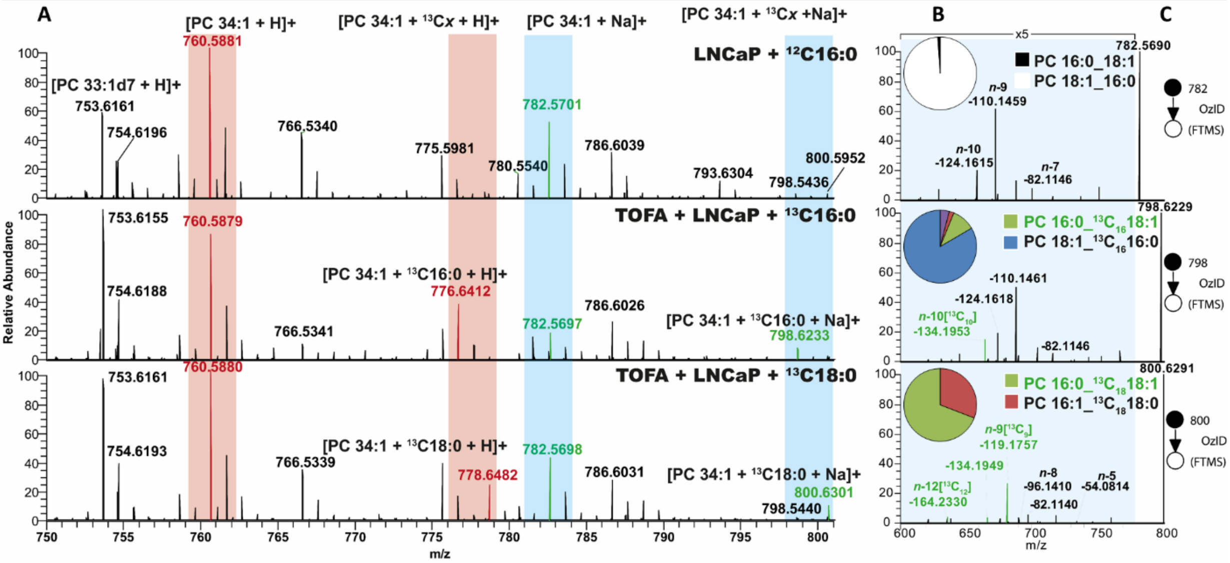

**Supplementary Fig. S13A:** Incorporation of labelled fatty acids into PC 34:1 and determined fatty acyl composition and double bonds of labelled PC 34:1. **A)** High resolution full mass spectrum of: (top) LNCaP supplemented with unlabelled palmitic acid; (mid) TOFA (SCD1/ACC1 inhibitor) treated LNCaP supplemented with labelled palmitic acid; and (bottom) TOFA treated LNCaP supplemented with labelled stearic acid. Spectra show unlabelled PC 34:1 as protonated (*m/z* 760) and sodiated (*m/z* 782) species, which likely would be derived from denovo lipogenisis. Incorporation of the labelled fatty acids can be seen as [M+16]+ (*m/z* 776 & 798) or [M+18]+ (*m/z* 778 & 800) isotopologues for labelled palmitic and stearic acids, respectively. High resolution mass accuracy was used to confirm assignments and eliminate incorrect assignment from isobaric interference (<Δppm 5). **B)** Data derived from CID/OzID (not shown). Pie charts showing the distribution of fatty acyl compositions present within either (top) unlabelled PC 34:1 or (mid and bottom) label incorporated PC 34:1. Within the labelled palmitic acid species (mid) it can be seen that the major lipid species is PC 18:1_(13C16)16:0, whereby the labelled fatty acid chain is saturated. In contrast, the labelled stearic acid species (bottom) is majority comprised of PC 16:0_(13C18)18:1 and shows that the labelled fatty acid has undergone cellular desaturation. **C)** The OzID mass spectra for the (top) unlabelled PC 34:1 or (mid and bottom) label incorporated PC 34:1. The product ions (shown as neutral losses (NL) from the precursor ion) are displayed in black if the OzID double bond fragmentation is associated with the unlabelled fatty acyl chain, or green if it is associated with the labelled chain (cf. Supplementary Fig. S13B for fragment assignment table). While the presence of the labelled *n*-10 product ion fragment (NL *m/z* 134) in the heavy palmitic acid lipid species (mid) suggests desaturation of a 16:0 to 16:1*n*-10 is subsequently elongated to 18:1*n*-10, the presence of this same *n*-10 product ion fragment in the labelled stearic acid species (bottom) reveals that stearic acid can be directly desaturated at the Δ8 position to yield an 18:1*n*-10 metabolite, which is being incorporated into phosopholipids.

| n- (\|1) | Unlabelled OzID fragment (NL) Aldehyde | Criegee | Isotop labelled OzID fragment (NL) Aldehyde | Criegee |
|---|---|---|---|---|
| 3 | 26.05204 | 10.05712 | 29.0621 | 13.06719 |
| 4 | 40.06769 | 24.07277 | 44.08111 | 28.08619 |
| 5 | 54.08334 | 38.08842 | 59.10011 | 43.1052 |
| 6 | 68.09899 | 52.10407 | 74.11912 | 58.1242 |
| 7 | 82.11464 | 66.11972 | 89.13812 | 73.14321 |
| 8 | 96.13029 | 80.13537 | 104.1571 | 88.16221 |
| 9 | 110.1459 | 94.15102 | 119.1761 | 103.1812 |
| 10 | 124.1616 | 108.1667 | 134.1951 | 118.2002 |
| 11 | 138.1772 | 122.1823 | 149.2141 | 133.2192 |
| 12 | 152.1929 | 136.198 | 164.2331 | 148.2382 |
| 13 | 166.2085 | 150.2136 | 179.2522 | 163.2572 |
| 14 | 180.2242 | 164.2293 | 194.2712 | 178.2762 |
| 15 | 194.2398 | 178.2449 | 209.2902 | 193.2952 |

**Supplementary Fig. S13B:** OzID neutral loss (NL) fragment masses for unlabelled monoisotopic unsaturated fatty acids (left) and isotope labelled unsaturated fatty acids (right). As the neutral loss *m/z* from OzID is relative to the methyl terminus and hence reflective of the number of carbons being lost from the lipid precursor, the neutral loss from labelled fatty acids differ from their unlabelled counterparts by the mass of a neutron multiplied by the number of carbons. This difference in *m/z* allows one to assign both the sites of unsaturation and the specific fatty acid substrates used during desaturation to confirm biosynthetic pathways.
